# Discovery of RNA-Reactive Small Molecules Guides Design of Electrophilic Modules for RNA-Specific Covalent Binders

**DOI:** 10.1101/2025.04.22.649986

**Authors:** Noah A. Springer, Patrick R. A. Zanon, Amirhossein Taghavi, Kisu Sung, Matthew D. Disney

**Affiliations:** Department of Chemistry, The Herbert Wertheim UF Scripps Institute for Biomedical Innovation & Technology, 130 Scripps Way, Jupiter, FL 33458, USA; Department of Chemistry, The Scripps Research Institute, 130 Scripps Way, Jupiter, FL 33458, USA

## Abstract

RNA is a key drug target that can be modulated by small molecules, however covalent binders of RNA remain largely unexplored. Using a high-throughput mass spectrometry screen of 2,000 electrophilic compounds, we identified ligands that react with RNA in a binding-dependent manner. RNA reactivity was influenced by both the reactive group and the RNA-binding scaffold. Electrophilic modules such as 3-chloropivalamide, bis(2-chloroethyl)amine, chloroacetamide, and N-acylimidazole that react with proteins also cross-linked to RNA, especially when paired with aromatic heterocycles, particularly those with a thieno[3,2-c]pyridinium core. These results suggest that electrophiles commonly used for protein targeting can also covalently modify RNA, potentially contributing to both on- and off-target effects. This insight enabled the design of an RNA-specific covalent compound by modifying a Hoechst scaffold, originally identified to bind DNA, to react selectively with the expanded triplet repeat RNA, r(CUG)^exp^, that causes myotonic dystrophy type 1 (DM1). Selectivity appears to arise from binding to the RNA major groove near the reactive site. Overall, this study highlights the potential of rationally designing covalent RNA-targeting small molecules.

**TOC Graphic:** 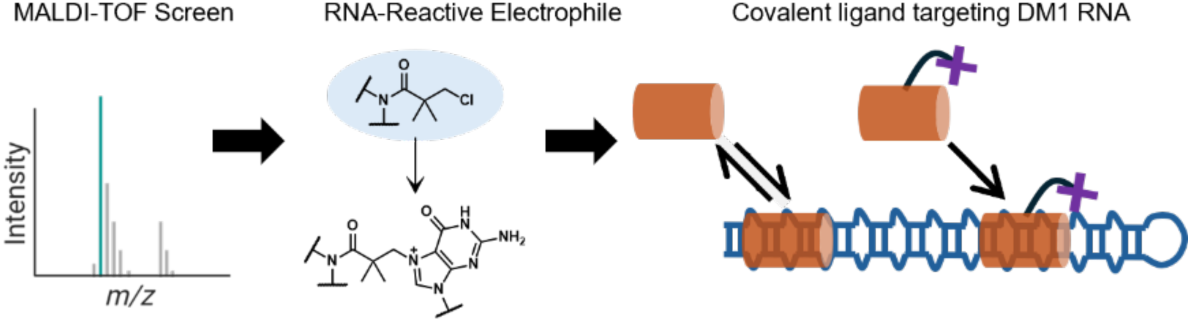

## INTRODUCTION

RNA structure, both in noncoding (nc) and coding transcripts, plays a crucial role in its function and dysfunction.^1, 2^ Thus, one way to modulate RNA function is by targeting its structure with small molecules. Various strategies,^3^ such as sequence-based design,^4, 5^ structure-based design,^6–11^ and high-throughput screening (HTS),^12–17^ have been employed to identify and optimize small molecules for RNA targeting. While most bioactive compounds act as simple RNA binders, heterobifunctional compounds such as degraders (direct^18–20^ or enzyme-mediated^21, 22^) and covalent ligands^23, 24^ have also emerged as RNA-targeted modalities. Previous efforts to develop covalent RNA-targeting compounds have primarily focused on well-known nucleic acid-reactive electrophiles, such as nitrogen mustards^23, 24^ or *N*-acylimidazoles,^25, 26^ attached to binding elements that confer affinity for specific RNAs. This study develops a methodology to rapidly identify small molecules that form stable, covalent adducts with RNA structures (Figure 1A).

**Figure 1.**
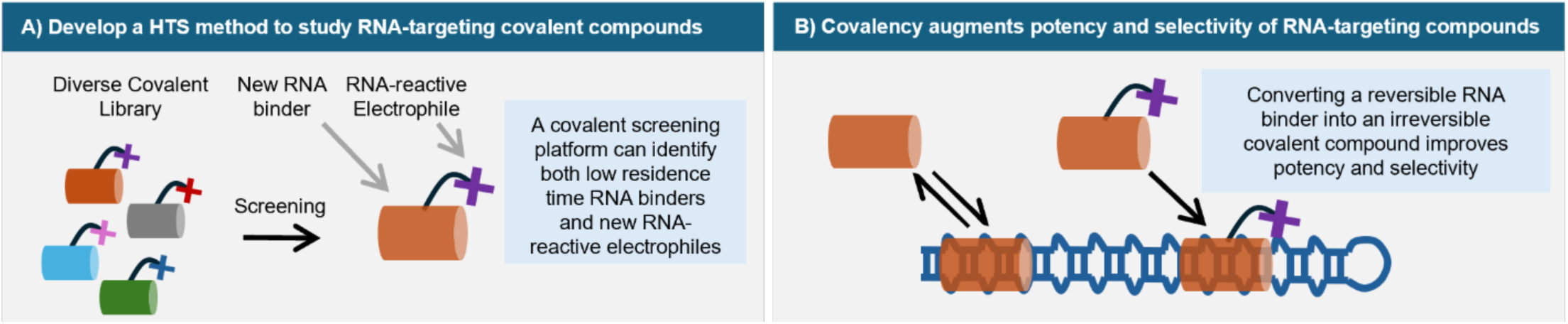
Identification and application of compounds covalently targeting RNA. **A)** This work seeks to develop a screening methodology to identify both RNA-reactive electrophiles and specific RNA-binding moieties. **B)** Irreversible covalent bond formation of small molecules targeting RNA confers improved potency and selectivity.

A matrix-assisted laser desorption/ionization-time of flight (MALDI-TOF)-based mass spectrometry (MS) screening approach was established, enabling the screening of a covalent compound library. This library included diverse electrophiles (e.g. α,β-unsaturated amides, chloroacetamides, imines) and potential RNA-binding elements. For this proof-of-concept study, the structure of a trinucleotide repeat expansion, r(CUG)^exp^ (where “exp” denotes an expanded repeat), associated with myotonic dystrophy type I (DM1),^27–29^ was chosen as a test case. In DM1 patients, r(CUG)^exp^ is found in the 3’ untranslated region (UTR) of the dystrophia myotonica protein kinase (*DMPK*) mRNA.^29, 30^ This repeat expansion forms an array of 1×1 nucleotide UU internal loops (5’C<u>U</u>G/3’G<u>U</u>C; where the loop nucleotides are underlined) that bind with high affinity to RNA-binding proteins (RBPs), such as muscleblind-like protein 1 (MBNL1).^31–33^ Sequestration of MBNL1 in toxic nuclear foci prevents it from performing its canonical function in alternative pre-mRNA splicing, leading to widespread dysregulation of alternative splicing in genes implicated in DM1 pathology, including the muscle-specific chloride ion channel 1 (*CLCN1*) and insulin receptor (*IR*).^34–36^

Herein, the reactivity of 2,000 small molecules was comprehensively evaluated, identifying several compounds that bind to and react with validated structural models of r(CUG)^exp^.^29–33^ A mildly reactive electrophile capable of alkylating guanosine was discovered, and its nucleic acid reactivity as well as its reactivity with thiols were profiled, which demonstrated specific reaction with the RNA repeat. Attaching this covalent module to a r(CUG)^exp^-binding small molecule imparted a significant improvement in potency and selectivity.

## RESULTS

### Development of Methods to Identify Small Molecule-RNA Adducts

Two methods amenable for high-throughput screening were developed to detect adducts between RNA and small molecules. To benchmark these approaches, a series of commercially available compounds (**1 – 9**) was selected based on the known reactivity of nitrogen mustards towards nucleobases (Figure 2A).^37–39^ These compounds contain a (2-chloroethyl)amine moiety that generates a reactive aziridinium electrophile *in situ* that reacts with biological nucleophiles.^37–39^ The small molecule panel includes the cancer chemotherapeutic chlorambucil (**1**), which has previously been shown to react with RNA *in vitro* and in cells when attached to an RNA-binding compound,^23^ uramustine (**2**), bendamustine (**3**), melphalan (**4**), and estramustine (**5**). Additionally, phenoxybenzamine (**6**), Astrazon Red 6B (**7**), Astrazon Pink FG (**8**), and a benzofuran mustard derivative (**9**), which have no known nucleic acid-alkylating ability, were also studied. The different aromatic or heterocyclic substructures found in this panel could bind to RNA and facilitate a proximity-induced reaction. In particular, **2** has a uracil moiety, **3** has an *N*-methyl benzimidazole, and **7** and **8** have a 1,3,3-trimethyl-3*H*-indol-1-ium, each of which could bind to RNA, likely with modest to weak affinity, as these nitrogen-containing heteroaromatic moieties are similar to known RNA binders.^40^ Further, the varying substituents can affect the formation and reactivity of the reactive aziridinium species.^41^ Each compound was tested for binding to and reacting with a r(CUG)^exp^ mimic, in which two 1×1 nucleotide UU internal loops were placed into a stem loop structure (**RNA1**). This minimized design allows the use of this construct in a variety of assays.

**Figure 2.**
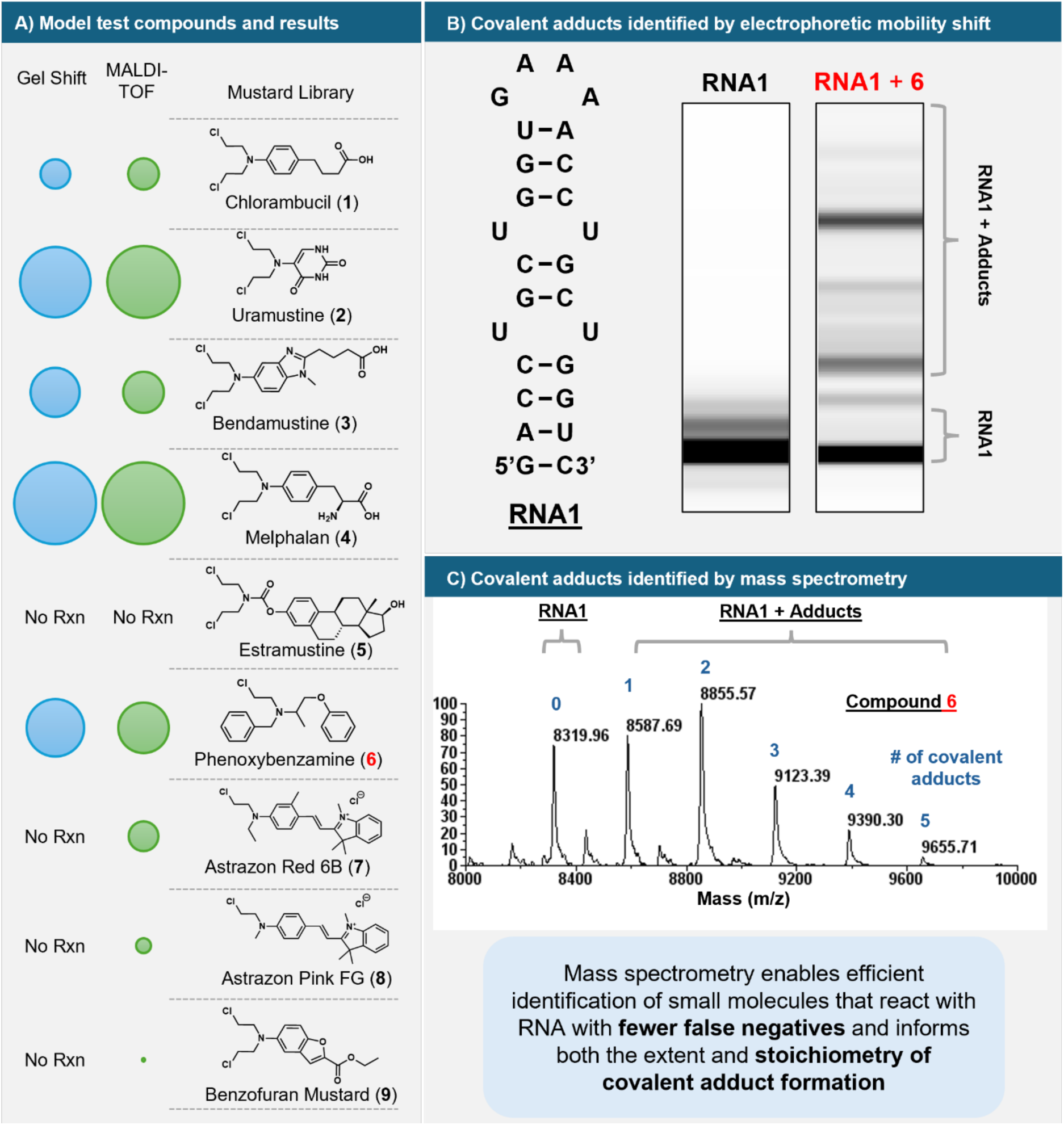
Development of methodologies to identify small molecules that react with RNA. **A)** Structures of the nine nitrogen mustard derivatives tested in both the electrophoretic mobility shift and MALDI-TOF mass spectrometry assays. Circles next to the compounds are sized according to their activity in each assay, a proxy for reactivity, where larger circles represent higher activity. “No Rxn” is used to denote compounds that showed no adduct formation in the corresponding assay. **B)** *Left*: Secondary structure of **RNA1** used for screening. The RNA contains two 1×1 nucleotide UU internal loops found in the trinucleotide repeat expansion r(CUG)^exp^, as well as base pairs at the 5’ and 3’ ends and a GNRA tetraloop for stable folding. *Right*: Results from screening using the gel shift assay by capillary electrophoresis for one of the hit compounds (**6**). Hits were defined as compounds that reduced the electrophoretic mobility of **RNA1** with signal intensities >4-fold higher than a DMSO (vehicle) control. **D)** MALDI-TOF mass spectrum for the reaction of **6** with **RNA1**. Each peak is labeled with its corresponding mass (m/z) and the stoichiometry of adduct formation with **6**.

The first method explored to assess covalent adduct formation with the r(CUG) repeat RNA was a electrophoretic mobility shift assay, where covalent modification could change the electrophoretic mobility of the RNA target, akin to a previously reported method used to identify and study reactive compounds targeting a riboswitch.^42^ After incubating **RNA1** (20 µM) with each nitrogen mustard (2 mM; 4 h at 37 °C), the change in electrophoretic mobility was assessed by capillary electrophoresis (CE) on a fragment analyzer. Of the nine compounds tested, five (**1 – 4**, **6**) significantly changed the electrophoretic mobility of the RNA by more than 4-fold above background signal from a DMSO control, with **2** and **4** showing the greatest extent of modification (Figure 2B & S1A). To provide a secondary assessment of reaction, the samples were also studied by denaturing polyacrylamide gel electrophoresis (dPAGE) and subsequent SYBR staining (Figure S1B), which identified the same five compounds as RNA-reactive. Interestingly, phenoxybenzamine (**6**), an FDA-approved α-adrenergic receptor inhibitor, induced shifts in the mobility of **RNA1** (Figure 2B), despite no prior direct evidence of nucleic acid alkylation for this drug.^43^ Neither the dPAGE nor CE methods detected adduct formation for some compounds that have an expected RNA-reactive moiety and a module that should confer at least some level of RNA binding, particularly at the high concentration of the small molecules (2 mM) present in the assays. For example, **7** and **8**, which each have a 1,3,3-trimethyl-3*H*-indol-1-ium, **9**, and **5** did not induce a gel mobility shift and were therefore, in these assays, scored as unreactive with **RNA1**.

As an orthogonal method to assess covalent RNA adducts, the reaction mixtures were subjected to MALDI-TOF mass spectrometry (Figure 2C). This analysis allows for the direct detection of adduct formation, reducing the potential for false negatives or false positives. All compounds that induced gel shifts also formed detectable adducts by MALDI-TOF. The most potent alkylators, **2** and **4**, formed between one to seven adducts per RNA molecule, with no observable unmodified RNA remaining (Figure S2). The other three hits from the CE and dPAGE assays – **6**, **3**, and **1** – resulted in stoichiometries of adduct formation of up to five, four, and three adducts per RNA, respectively (Figure S2). Interestingly, three additional nitrogen mustards (**7 – 9**) that were inactive in the dPAGE and CE assays alkylated **RNA1** as determined by MALDI-TOF MS (Figures 2A & S2). Indeed, eight of the nine nitrogen mustards alkylated **RNA1**, with **5** being the only exception (Figure 2A). The electron-withdrawing carbamate group of **5** likely prevents the formation of the reactive aziridinium intermediate and therefore diminished the likelihood it will react with RNA. Previous studies suggest that the mechanism of action of estramustine (**5**), unlike other anti-cancer nitrogen mustards, is based on anti-tubulin activity rather than DNA interstrand cross-linking.^44–48^ Collectively, these data show that MALDI-TOF analysis is superior to electrophoretic mobility shift assays to discover small molecules that react with RNA, particularly in its ability to eliminate false negatives and to assess the stoichiometry of modification.

### Development and Implementation of a High-Throughput Screen of Small Molecules that React with RNA Using MALDI-TOF

In proteins, the pK_a_ values of nucleophilic amino acids such as cysteines are perturbed due to their local environment, rendering them more reactive.^49^ Through specific interactions of the covalent ligand with the binding pocket, its electrophile can be adequately positioned to covalently engage the protein nucleophile.^50–52^ Further, covalency is commonly used to identify fragments that bind biomolecules with low affinity.^53–55^ We envisioned that covalent compounds might also act similarly for RNA targets and that by screening a collection of such small molecules, new RNA-reactive modules as well as RNA-binding elements might be discovered.

To adapt the MALDI-TOF method for high-throughput screening, an efficient, reproducible, and small-scale method for removing salt and other contaminants from RNA samples was required. Here, solid-phase reverse immobilization (SPRI)-based magnetic beads, commonly used for the preparation of sequencing libraries, were chosen. These beads use polyethylene glycol (PEG) and high salt concentrations to precipitate RNA onto carboxy-coated paramagnetic beads. Optimization of the method—including the addition of isopropanol to facilitate precipitation of small RNA constructs, increasing the number of washes to thoroughly remove salt and PEG contaminants, and miniaturizing the sample volume—enabled screening and RNA purification from 5 µL reactions in 384-well plates (Figure S3). Full details for this method can be found in the Supporting Information.

A 2,000-compound electrophile library (Extended Data Table 1) was carefully designed from both the perspective of the electrophile and potential RNA-binding element. In particular, each compound harbors a module with potential to react with biomolecules and binding elements to facilitate interaction with RNA (Figures 3A & S4). Various electrophiles are represented in the covalent library, ranging from known nucleic-acid reactive *N*-acylimidazoles^56^ and bis(2-chloroethyl)amines^39^ to traditionally protein-targeted chloroacetamides^57, 58^ and α,β-unsaturated amides (Figures 3A & S4).^59^ Various small molecules that bind to r(CUG)^exp^ have been reported in the literature,^17, 60–66^ and the library designed herein shares structural similarities with them. The library contains 98 unique nitrogen-containing heteroaromatic ring systems, commonly found in RNA-binding small molecules,^40^ with benzothiazole, pyridine, pyrazole, benzoxazole, and indole among the most common in the library (Figure S4). Thus, these molecules might induce covalency via induced proximity, rather than non-specific reactivity. The physicochemical properties of the library overlap with a subset of the chemical space occupied by small molecules in the DrugBank database,^67^ as assessed by Uniform Manifold Approximation and Projection (UMAP) analysis (Figure 3A).^68^ Additionally, most compounds in the library (97%) follow Lipinski’s Rule of 5^69^ and have a topological polar surface area (TPSA) of less than 140 Å^2^ (99.8%) (Figures 3A & S4).^70, 71^

**Figure 3.**
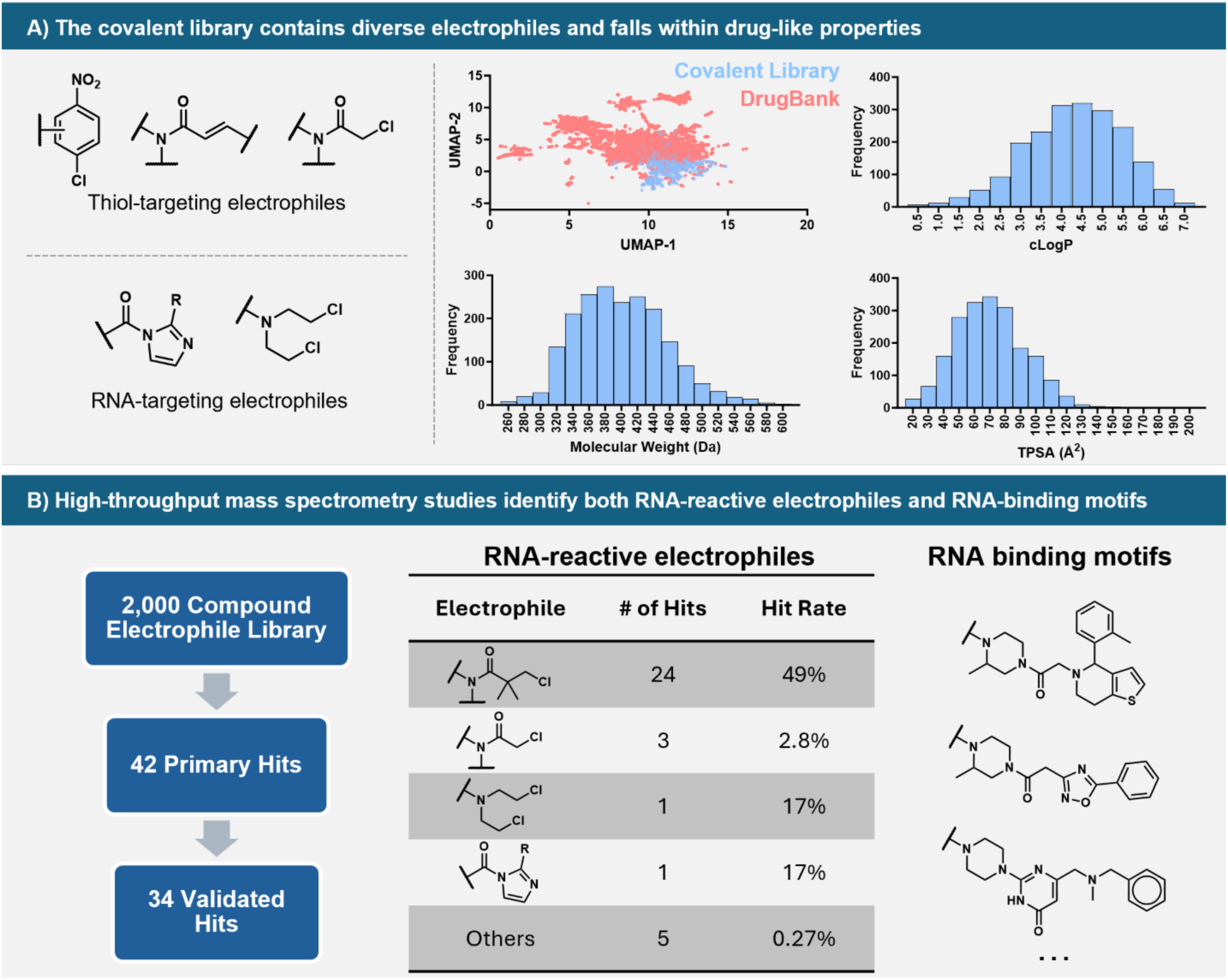
Composition and screening of a 2,000-compound covalent library. **A)** *Left*: The library is composed of both traditionally protein-targeted electrophiles and those known to react with RNA. *Right*: Uniform Manifold and Projection (UMAP) structural analysis of the library^68^ shows that the electrophile library (blue) overlaps with the properties of a subset of the DrugBank library.^67^ Histograms for several physicochemical properties of the electrophile library show that most compounds fall within desirable drug-like values of molecular weight <500 Da, cLogP <5, and TPSA <140 Å^2^. **B)** Screening of the electrophile library for reacting with **RNA1** identified 34 replicable, validated hits. Of these, 24 compounds contained a 3-chloropivalamide electrophile as the reactive component. While the RNA-binding motifs varied, several of the most potent covalent compounds contained heteroaromatic rings common among RNA-binding modules, with their structures shown to the right.

Adduct formation was assessed by incubating each compound (1 mM) with **RNA1** (20 µM; 12 h at 37°C). Long incubation times were used to maximize adduct formation. After clean-up with SPRI magnetic beads, covalent adducts of 42 small molecules with **RNA1** were detected by MALDI-TOF, affording a hit rate of 2.1% (Extended Data Table 1 & Figure S5). Of these, 34 were validated by replicating the assay (81% validated; Extended Data Table 2, Figures 3B & S5). To identify the small molecules that react with **RNA1** to the greatest extent, the 34 validated hits were studied in a dose response at concentrations of 10, 100, and 1000 µM. From this collection, 18 compounds formed covalent adducts at doses lower than 1 mM (Figure S5).

Of the 34 validated hits, three were chloroacetamides, which were previously unknown to react with RNA,^42, 72^ although chloroacetamides have been used to chemically cross-link interactions formed between RNA and RBPs.^73^ Two compounds containing known RNA-reactive electrophiles – one nitrogen mustard and one *N*-acylimidazole – reacted with **RNA1**. Most strikingly, 24 of the small molecules that formed covalent adducts with **RNA1** contained a 3-chloropivalamide which underwent substitution of the chloride substituent (Figure 3B). Only 49 of the 2000 small molecules in the covalent library contained this reactive module; the difference between the hit rate of 3-chloropivalamides targeting **RNA1** (24/49; 49%) as compared to other electrophiles in the library (10/1951; 0.5%) is statistically significant (p <0.0001). A subset of compounds with the 3-chloropivalamide electrophile contained a central 4-(piperazin-1-yl)pyrimidine motif (14/49) and the extent of reactivity towards **RNA1** appeared to depend upon the substituents on the pyrimidine ring of the RNA-binding element. Bulky aromatic substituents at the 2-position eliminated reactivity towards the RNA (Figure S6). This structure-activity relationship suggests that the RNA-binding module conjugated to the 3-chloropivalamide electrophile is important for covalent bond formation with **RNA1**.

Compound **10** showed the greatest extent of adduct formation to **RNA1** in the primary screen (1.03 for **10,** 0.14 ± 0.15 on average for the other hits; Figure S5), secondary validation (1.74 for **10**, 0.21 ± 0.23 on average for other validated hits; Extended Data Table 2), and dose-response studies (1.52 for **10**, 0.15 ± 0.17 on average for other validated hits at 1 mM; Figure S5), as determined by the ratio in peak intensities of RNA containing the adduct to unmodified RNA. For this reason, **10** was carried forward to additional studies. Further examination of the screening and validation data indicated that the detected adduct with **RNA1** was nearly 3 Da less than expected (Extended Data Table 2). Analysis of the stock used for screening revealed that it was a mixture of the four diastereomers of the intended compound (**10a – 10d**) and racemic mixtures of two oxidized forms: iminium (**10e, 10f**) and pyridinium (**10g, 10h**) (Figures 4 & S7), suggesting that one of the oxidized forms (**10e – 10h**) might be responsible for the observed activity. Compound **10a-d, 10g,** and **10h** were synthesized and studied for covalent adduct formation with **RNA1** by MALDI-TOF analysis; synthetic difficulties prevented analysis of **10e** and **10f**. Indeed, the oxidized derivatives **10g** and **10h** formed adducts with the RNA while **10a – 10d** showed no reactivity (Figures 4 & S7). Both **10g** and **10h** showed time-dependent reaction with **RNA1** over the span of 23 h, with the (*R*) enantiomer (**10h**) resulting in a modestly higher extent of reaction at extended time points (0.97±0.08 ratio of peak intensities vs. 0.67±0.02; Figure S7).

**Figure 4.**
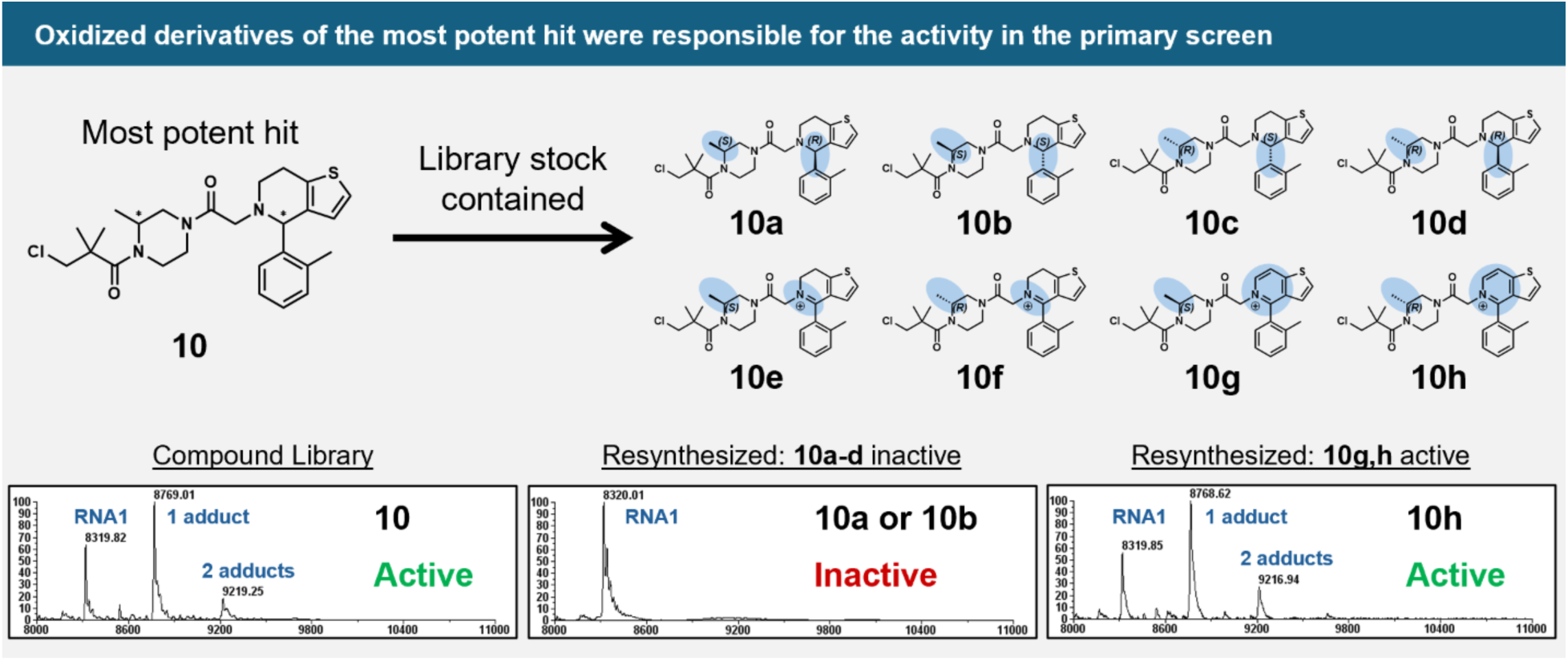
Identification of active component of the most potent hit, 10. The stock of **10** in the compound library was a mixture of its four diastereomers **10a – 10d** as well as two oxidized forms **10e – 10h**. While **10a – 10d** showed no reactivity upon resynthesis, **10g** and **10h** both recapitulated the screening data, with little stereoselectivity.

Enzymatic digestion of **10h**-modified **RNA1** to nucleosides revealed that the compound reacts with the guanine base (Figure S8). Although unequivocal assignment could not be made from the mass spectra, the N-7 position is known to be intrinsically prone to modification by many electrophiles including nitrogen mustard derivatives,^74^ dimethyl sulfate (DMS),^75, 76^ and diethyl pyrocarbonate (DEPC).^76^ Furthermore, guanosines in the r(CUG)^exp^ structure are engaged in Watson-Crick pairing, leaving N7 accessible in the major groove. As many ligands that bind to RNA recognize the major groove,^15^ a proximity-induced reaction of **10h** with N7-G could be rationalized. To support the N-7 of guanine as the reactive site, **RNA1** modified with **10g** was reduced with sodium borohydride, which induces an abasic site at guanines alkylated at the N-7.^77, 78^ MALDI-TOF spectra of the reduced RNA showed the disappearance of peaks corresponding to the covalent adducts of **10g** and the appearance of a previously unobserved peak corresponding to an abasic site generated from the loss of a guanine (Figure S8). This same peak was not observed when **RNA1** lacking the covalent adducts was treated identically (Figure S8; DMSO-treated samples), suggesting the reduction-induced abasic site requires covalent modification of the RNA.

### Studying Structure-Reactivity Relationships of the 3-Chloropivalamide Electrophile with RNA

The reactivity profile of the 3-chloropivalamide electrophile was more thoroughly characterized by studying its reaction with various nucleophiles, including thiols and nucleobases. Analysis of the attachment points of the electrophile to the putative RNA-binding modules in the library suggested that reactivity depended on the amide substitution. For example, none of the eight anilides reacted with **RNA1**, while all seven of the methyl-piperazine derivatives formed adducts (Figure S9). Although these differences could be due to variations in the RNA-binding components, it is likely that the amide properties influence the electrophile’s reactivity, as has been reported for other electrophiles such as chloroacetamides.^54^

To assess effects on the reactivity of 3-chloropivalamides, a set of eight electrophiles lacking an RNA-binding element (**11 – 18**) was synthesized with varying amide substructures, including primary and secondary alkyl amines and aniline (Figure 5A). To study the rate of reaction of the electrophiles with guanosine, a colorimetric model system with 4-(4-Nitrobenzyl)pyridine (NBP) was utilized (Figures 5B & S10).^79^ NBP is an aromatic nitrogen nucleophile commonly used to detect and assess alkylating agents due to its similar reactivity with N7-G, as indicated by their nearly identical Swain-Scott nucleophilicity values.^80^

**Figure 5.**
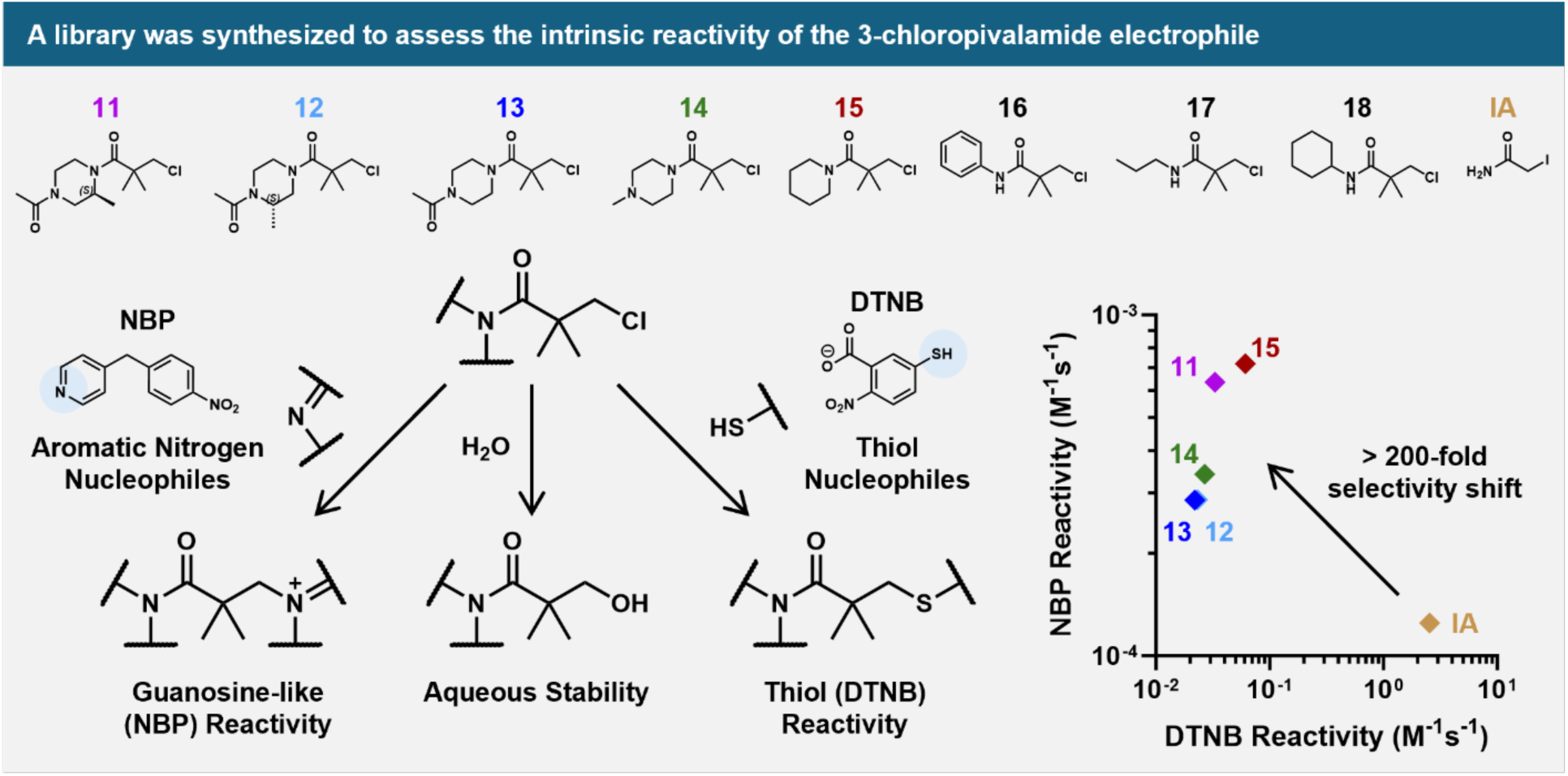
Assessing the intrinsic reactivity of the 3-chloropivalamide electrophile. *Top*: Eight compounds containing cyclic secondary amines, primary amines, and aniline were coupled to the 3-chloropivalamide electrophile to assess the influence of the amide substitution on the reactivity of the electrophile. Their intrinsic reactivity was compared to the promiscuously reactive electrophile iodoacetamide (IA). *Left*: The reactivity of the compounds was measured by three assays: (i) a colorimetric assay with NBP was used to assess reactivity towards guanosine-like nucleophiles; (ii) the aqueous stability of the compounds was assessed using LC/MS; and (iii) the reactivity towards thiols was assessed using a colorimetric assay with Ellman’s reagent (DTNB). *Right*: The cyclic tertiary amides (**11 – 15**) were 200-fold more selective towards guanosine-like nucleophiles (NBP) than thiols (DTNB) relative to iodoacetamide.

Reactions between the electrophiles **11 – 18** as well as iodoacetamide and NBP showed that the cyclic tertiary amides (**11 – 15**) were the most reactive, while other derivatives (**16 – 18**) showed negligible alkylation after 28 h (Figures 5B & S10). To calculate second order rate constants for the reaction with NBP, the hydrolytic stability for each electrophile (as measured by LC-MS; Figures 5B & S11) was determined, where half-lives ranged from 2 h to >4 d (Table S1). After incorporating hydrolytic stability, rate constants for the reactions of **11 – 15** with NBP ranged from 2 × 10^-4^ M^-1^s^-1^ to 7 × 10^-4^ M^-1^s^-1^, which is 2-6-fold more reactive than iodoacetamide (1.2 ± 0.1 × 10^-4^ M^-1^s^-1^).

Nucleophilic thiols such as glutathione and reactive cysteines are other potential targets for the 3-chloropivalamide electrophile in cells. To assess intrinsic thiol reactivity, a colorimetric assay with 5,5-dithiobis-(2-nitrobenzoic acid) (DTNB) was used^54^ (Figures 5B, S12 & Table S1). To benchmark reactivity, studies were first conducted with the promiscuous thiol-reactive compound iodoacetamide, affording a second order rate constant of 2.55 ± 0.04 M⁻¹s⁻¹, in agreement with a previously reported value (2.6 ± 0.1 M^-1^s^-1^).^81^ The amide substituents within **11 – 18** significantly influenced thiol reactivity by more than 60-fold, with second-order rate constants ranging from 6.1 ± 0.1 × 10⁻² M⁻¹s⁻¹ to <1 × 10^-3^ M⁻¹s⁻¹ (Figures 5B, S12 & Table S1). In line with the NBP reactivity assay, cyclic tertiary amides (**11 – 15**) were the most reactive, followed by the anilide (**16**) and secondary amides (**17, 18**), where **18** showed no detectable reactivity. These trends are largely consistent with those observed for chloroacetamides^54^ and correlate with the reactivity of the hits that emerged from the reactivity profiles of the 2,000-member covalent compound library toward **RNA1** (Figure S9). Furthermore, a previous screen of more than 750 chloroacetamides revealed that chloroacetamides are, on average, 20 to 25-fold less reactive than iodoacetamide.^54^ The 3-chloropivalamide electrophiles are 40- to >2,000-fold less reactive towards thiols than iodoacetamide (Figure S12 & Table S1); thus, they are less reactive towards thiols than the average chloroacetamide. When the relative reactivity of iodoacetamide and **11 – 15** towards both guanosine-like (NBP) and thiol (DTNB) nucleophiles are compared, a more than 200-fold shift in selectivity toward guanosine-like nucleophiles was observed for the 3-chloropivalamide electrophile (Table S1 & Figure 4B). Collectively, these studies suggest that the identified electrophiles are unlikely to be promiscuously reactive with thiols in the proteome or other cellular components and have high potential to be used as a module on an RNA-binding compound to induce specific reaction with an RNA target.

### Using Covalent Reactivity to Inform Design of Non-covalent r(CUG)^exp^ Binders

The covalent screen revealed not only RNA-reactive electrophiles, but also RNA-binding small molecules, suggesting that the extent of reaction with **RNA1** depended on both the reactivity of the electrophile and the non-covalent affinity of the binding component. To assess the contribution of the RNA-binding component of **10g** and **10h** to **RNA1** modification, the stoichiometry of adduct formation between **10h** and a control compound lacking the RNA-binding moiety, **11**, was compared. At equivalent concentrations (2 mM), **10h** showed approximately 5-fold greater adduct formation than **11** (0.83 vs. 0.18 adducts per RNA, respectively); when **11** was used at a 5-fold higher concentration (10 mM), the stoichiometry of adduct formation with **RNA1** was comparable (0.93 adducts per RNA) to that observed for 2 mM of **10h** (0.83 adducts per RNA; Figure S13) (n = 1). This modest 5-fold increase in adduct formation may be attributed to the weak binding affinity of the non-covalent portion for **RNA1** or improper orientation of the electrophile toward the RNA target. The binding of a non-covalent derivative of **10g**, **10i**, was studied by nuclear magnetic resonance (NMR) spectroscopy, monitoring changes in the imino protons present in **RNA1**, particularly in the 1×1 UU internal loops, whether chemical shift perturbation or peak broadening; however, no significant binding (very weak) was observed (100 μM of **10i** and 50 μM of **RNA1**; Figure S14).

Knowledge of the reactivity of the 3-chloropivalamide electrophile with the guanine base, presumably at N-7, informed design of higher affinity RNA-binding modules. That is, the electrophile in **10g/10h** (reacted to the greatest extent with **RNA1**) was replaced with moieties capable of facilitating hydrogen bonding with the Hoogsteen face of guanine, with which the electrophile reacts (Figure S14). These moieties include histidine (**10j**, **10k**) or 4-oxoproline (**10l**, **10m**), which could form hydrogen bond interactions to the Hoogsteen face in a similar way to base triple interactions that occur naturally by interacting with the major groove^82, 83^ and have been used to engineer triple helical structures.^84–87^ Binding of **10j – 10 m** to **RNA1** was monitored by NMR spectroscopy, as completed for the original core scaffold (**10i**), which induced limited changes in the imino proton spectrum of **RNA1**. In contrast to **10i**, more significant changes were observed upon addition of both **10l** and **10m** at the same concentration but not **10j** or **10k** (Figure S14). Small molecule **10m** induced the most significant changes, as assessed by the largest decrease in peak intensity for the imino protons in 1×1 nucleotide UU internal loops (Figure S14).

Collectively, these studies enabled the identification of a low-affinity binding core through covalent modification of its RNA target. Information about the positioning of an electrophilic moiety within the compound and its site of reactivity within an RNA target enabled design of a higher affinity RNA-binding module; that is, the electrophile was replaced with non-reactive moieties capable of forming interactions with guanosine residues. Additional studies of this approach and this compound set are required to better understand how to utilize this scaffold to target r(CUG)^exp^ and provide potentially bioactive ligands.

### Enhancing Reactivity Using Known r(CUG)^exp^-Binding Ligands

Given the high concentrations (>1 mM) and low extent of reactivity (incomplete modification after 24 h) of **10g** and **10h** with **RNA1**, a previously validated r(CUG)^exp^-binding compound, **H**^88^ (a derivative of the DNA-binding small molecule Hoechst), was conjugated to the 3-chloropivalamide electrophile to improve the extent of covalent adduct formation (Figure 6A). A high affinity nucleic acid binder, Hoechst recognizes the minor groove of DNA^89, 90^ as well as non-canonically paired internal loop regions in RNA, particularly the 1×1 nucleotide UU internal loop found in r(CUG)^exp^ amongst a variety of other RNA targets.^88^ **H** was used as a module in a series of multivalent ligands that specifically bind r(CUG)^exp^ with high affinity and specificity.^91^ While an **H** homodimer alleviates DM1-associated phenotypes in a cellular model, **H** itself is biologically inert.^92^

**Figure 6:**
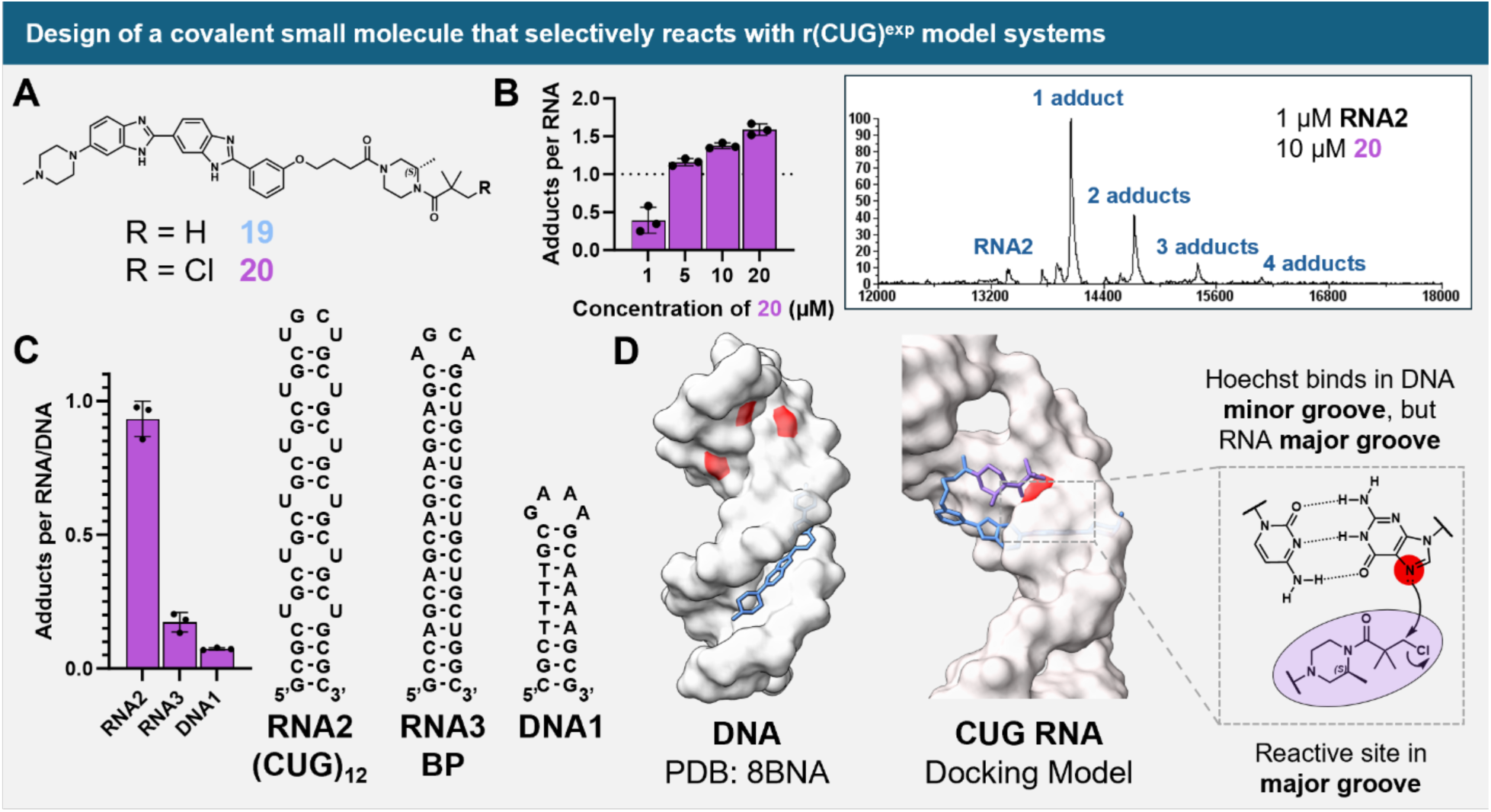
Developing a more potent covalent r(CUG)^exp^ binder. **A)** A previously validated r(CUG)^exp^ binder was appended to both the identified electrophile **11** and a non-covalent control lacking the chloride to generate the r(CUG)^exp^-targeting compounds, **20** and **19**, respectively. **B)** *Left*: Compound **20** dose-dependently formed adducts with **RNA2** (1 µM). *Right*: A representative mass spectrum showing near-complete reaction of **20** (10 µM) with the **RNA2** (1 µM). **C)** Compound **20** (20 µM) reacted to a much lesser extent with **RNA3** (20 µM), which lacks the 1×1 UU loops, and **DNA1** (20 µM) than **RNA2** (20 µM). **E)** Low reactivity towards DNA may be explained by differential modes of binding. The **H** parent compound is a known minor groove binder in DNA (*Left*),^94^ but molecular docking of **19** to a model of r(CUG)^exp^ (*Right*) suggests that the electrophile can be presented to the major groove, where the proposed reactive N7-guanine resides.

Due to the favorable reactivity profile of **11** in the NBP alkylation assay (Figure S10), a conjugate between **H** and **11**, compound **20**, as well as a non-covalent control **19**, were synthesized (Figure 6A). A longer r(CUG) repeat construct (r(CUG)_12_; **RNA2**) was chosen to better represent the biologically relevant r(CUG)^exp^ structure (Figure 6). [Note: this RNA was not used in the library screen because it requires more stringent washing necessary for clear detection by MALDI-TOF MS due to its larger molecular weight]. Compound **20**, like **10g**, specifically modified the guanine bases of **RNA2** (Figures S8 & S15A). Compound **20** also showed dose-dependent labeling of **RNA2**; in particular, at 10 µM of **20**, nearly complete alkylation of **RNA2** (1 µM) was observed (Figures 6B & S15B), a higher extent of adduct formation than observed with 1 mM of **10g** or **10h** with **RNA1** (Figure 4). The greater extent of adduct formation at a 100-fold lower concentration of compound represents a more than 100-fold improvement of *in vitro* potency of the r(CUG)-targeting compounds. The RNA-binding component of **20** was necessary to facilitate these covalent adducts, as no reaction of **11**, which lacks the RNA-binding **H** component, with **RNA2** was observed until at least 2 mM of the small molecule was added (Figure S15C). Additionally, this reaction was selective for RNAs containing the 1×1 UU motif, as the fully base-paired **RNA3** (0.17 ± 0.04 adducts per RNA) showed significantly attenuated reactivity towards **20** than **RNA2** (0.93 ± 0.07 adducts per RNA, Figure 6C). Further, when **20** was incubated with a mixture of **RNA2** and **RNA3**, only covalent adducts to **RNA2** were observed (Figure S15D). The selectivity of the reaction of **20** with **RNA2** was examined in the presence of 1 or 10 mM glutathione, a highly abundant thiol nucleophile present in cells. Under these conditions, only a modest (∼14%) decrease in reaction with **RNA2** was observed (Figure S15E), suggesting that these compounds could be valuable for studying cellular models of DM1 by specifically targeting r(CUG)^exp^ and potentially improving DM1-associated defects.

Because **20** contains a binding core derived from the DNA-binding Hoechst dye, the ability of the compound to react with DNA was studied. Two DNA constructs (**DNA1** and **DNA2**) were designed which contain both the preferred Hoechst binding sites (A/T stretches) and adjacent guanines for potential covalent adduct formation (Figures 6A & S15F). The binding of **19** (non-reactive to prevent potential interference of reactivity in the assay) to **DNA1**, **DNA2**, and **RNA2** was evaluated by monitoring change in the intrinsic fluorescence of **19**, resulting in binding EC_50_s of 250 nM (95% confidence interval (CI): 205-295 nM), 74 nM (95% CI: 53-98 nM), and >20 µM, respectively (Figure S15F). Despite the higher EC_50_ for **19** binding to **RNA2** compared to both DNA oligonucleotides, the propensity of **20** to form covalent adducts with **RNA2** (0.93 ± 0.06 adducts per RNA) is 13 times higher than with **DNA1** (0.073 ± 0.004 adducts per DNA) (Figure 6C). This difference in reactivity is likely not due to differences in intrinsic reactivity of the electrophile towards DNA or RNA, as the electrophile lacking the RNA binder (**11**) reacts with **DNA1** only ∼2-fold less than with **RNA2**, despite **RNA2** containing 15 potentially reactive guanine residues and **DNA1** only having six (Figures S15C & S15G).

To gain insight into these observations, molecular modelling was used. These studies revealed that **19** binds to the 1×1 nucleotide UU internal loops and can display the electrophilic site into the major groove of r(CUG) repeats, in contrast to binding in the DNA minor groove^89, 90^ (Figure 6D). The lowest free energy pose of **19** bound to a r(CUG) repeat containing RNA places the electrophilic site in proximity to the N-7 position of guanine at the base pair adjacent to the binding site (Figure 6D). Out of docked poses generated using AutoDock-GPU,^93^ the electrophile is positioned into the major groove for half of them, including 6 out of the 8 lowest free energy poses (Figure S16A). In contrast, when **19** is docked to a previously solved structure of DNA-bound Hoechst 33258 (PDB: 8BNA),^94^ all docked structures position the molecule and electrophilic site within the minor groove (Figure S16B). Further, the predicted binding free energies are ∼8 kcal/mol lower for docking of **19** to DNA (−18.92 kcal/mol) than the model r(CUG) RNA (−10.45 kcal/mol), supporting the measured EC_50_s (Figure S15F). Thus, despite higher affinity, non-covalent binding with DNA, the lack of reactivity for **20** with DNA is likely due to the positioning of the reactive module in the minor groove, away from the reactive site, presumably the N-7 of guanine. In contrast, the binding of **20** to the RNA positions the reactive module nearby the guanine N-7 to facilitate reactivity. These results suggest that the specificity of a reactive, RNA-targeting small molecule is influence by both its affinity and binding mode, that is proper position of the reactive group, or positional reactivity.

## DISCUSSION

Covalent approaches to target biomolecules have transformed chemical biology and medicinal chemistry. A wide variety of protein-targeted covalent binders have expanded the druggable space, informed biology, and garnered US Food & Drug Administration (FDA) approval.^95, 96^ Developing such approaches broadly for RNA targets could therefore have significant potential.^3^ Yet, there have been relatively few studies of small molecules that covalently target RNA.^23, 97, 98^ One of the challenges with specific reactivity with RNA targets is that the bases have relatively similar reactivity profiles, unlike amino acid side chains which vary significantly.^96^ The development and optimization of a MALDI-TOF MS screening methodology allowed a broader view of the potential for RNA to be covalently targeted. Indeed, many compounds that have been used for specific protein reactivity also react with RNA, suggesting that RNA should be considered an on- or off-target in these and other screens, as has been previously put forward for non-covalent ligands.^99^

A 3-chloropivalamide electrophile was identified as a common motif among hit small molecules that react with RNA. Using these results coupled with chemical synthesis and reactivity assessment elucidated characteristics of a molecule that can provide covalent ligands for RNA targets. Importantly, attachment of this electrophile to a dual DNA- and RNA-binding compound afforded selective reaction with RNA *in vitro*, particularly the 1×1 nucleotide UU internal loop that is present in r(CUG)^exp^. This conversion was possible by exploiting the differences in binding mode between the targets, where the electrophile is positioned nearby an N-7 of guanosine in the RNA but not in the DNA target. Thus, positional reactivity could be a general strategy for improving the selectivity of binding small molecules, as observed for targeted degradation approaches, either directly^100^ or via enzymatic recruitment.^101^

Covalent screening in the format presented herein could have broader implications. By studying a larger and more diverse covalent compound library, it is possible to identify not only reactive molecules but also compounds that bind RNA, including those of low affinity or short residence times. Moreover, information about binding sites and the positioning of reactive molecules can guide the design of compounds that recognize the target noncovalently, leading to improved activity. There are likely many additional approaches that can emerge based on these studies and earlier ones on covalent chemistry for RNA.^23, 24^ The development of covalent binders for RNA targets not only could provide bioactive compounds but could also be used to study the molecular recognition of RNA structures in cells in an unbiased way, which could be lead optimized into bioactive ligands targeting RNA.^102–104^

## ACKNOWLEDGMENTS

This work was funded by National Institutes of Health (R35 NS116846 to M.D.D.), The Muscular Dystrophy Association (Grant ID 1069959 to M.D.D.) and the German Research Foundation (DFG) through a Walter Benjamin fellowship (#515396515 to P.R.A.Z.). This material is based upon work supported by the National Science Foundation Graduate Research Fellowship under Grant No. 2235200 (to N.A.S.). Any opinion, findings, and conclusions or recommendations expressed in this material are those of the authors(s) and do not necessarily reflect the views of the National Science Foundation.

