## Supporting Information for "Discovery of RNA-Reactive Small Molecules Guides Design of Electrophilic Modules for RNA-Specific Covalent Binders"

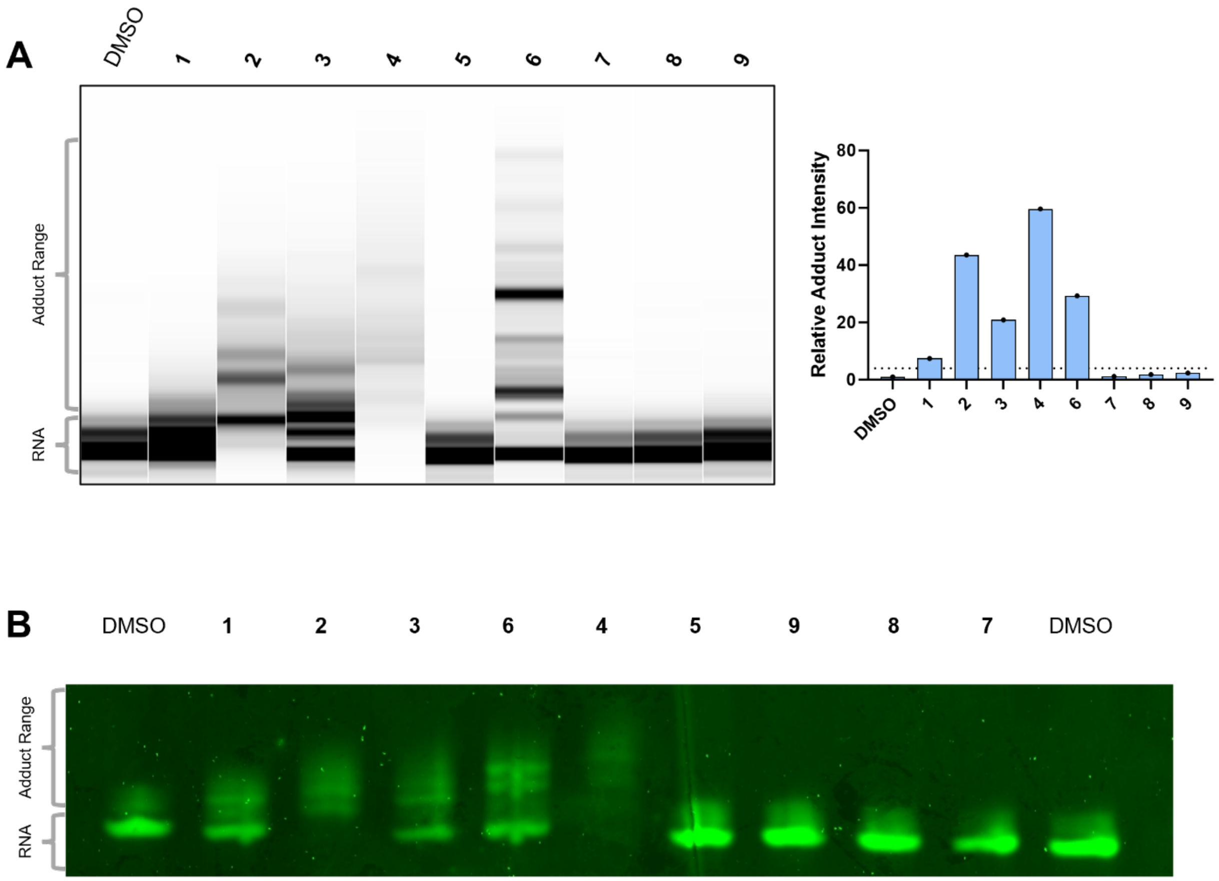


**Figure S1**. **Electrophoretic mobility shift assays to monitor covalent adduct formation**. **A)** Following covalent adduct formation between 20 μM **RNA1** with 2 mM of each nitrogen mustard (numbered according to Figure 1) or a DMSO-treated sample (10% (v/v) DMSO was present in all samples), samples were assessed for electrophoretic mobility shifts via capillary electrophoresis (CE) using a fragment analyzer. *Left*: An electropherogram showing the gel-shift induced following covalent adduct formation for each nitrogen mustard (n = 1 independent experiment). *Right*: Each lane was quantified in ImageJ by taking the ratio of the signal above the primary band for **RNA1** ("Adduct Range") to the signal for corresponding to **RNA1** ("RNA"). Hits were defined as compounds capable of inducing at least 4-fold signal relative to DMSO (dashed line). **B)** Gel image showing an electrophoretic mobility shift induced for the same samples in **(A)**, as assessed by denaturing urea polyacrylamide gel electrophoresis (dPAGE) followed by SYBR staining. Samples were not quantified, but assessed qualitatively, due to inconsistent background staining influencing the measurement.

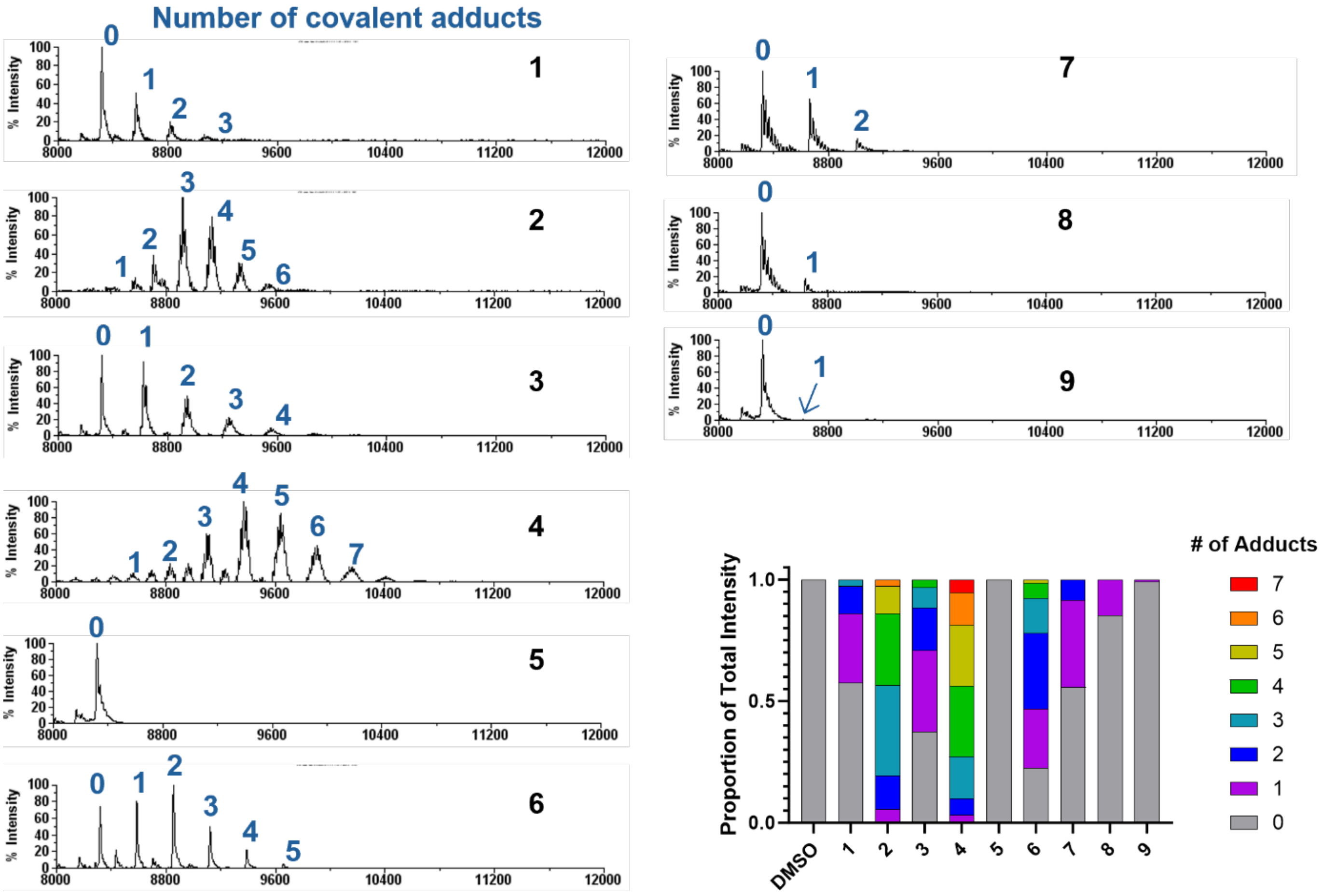


**Figure S2. MALDI-TOF analysis to assess covalent adduct formation**. MALDI-TOF spectra for covalent adducts to **RNA1** by the nitrogen mustard library (**1-9**) showed adduct peaks for all compounds except **5** (estramustine). Peaks are labeled with the corresponding number of covalent adduct to a single RNA. Broadening with multiple peaks was observed for divalent mustards (**1-4**), which corresponds to varying hydrolysis states of the second chloroethyl group on the molecule, as observed previously.[1] The abundance of each adduct was quantified by peak height in the mass spectrum, and the proportion of the total intensity in each spectrum that corresponds to each stoichiometry of adduct is plotted in the bottom right.

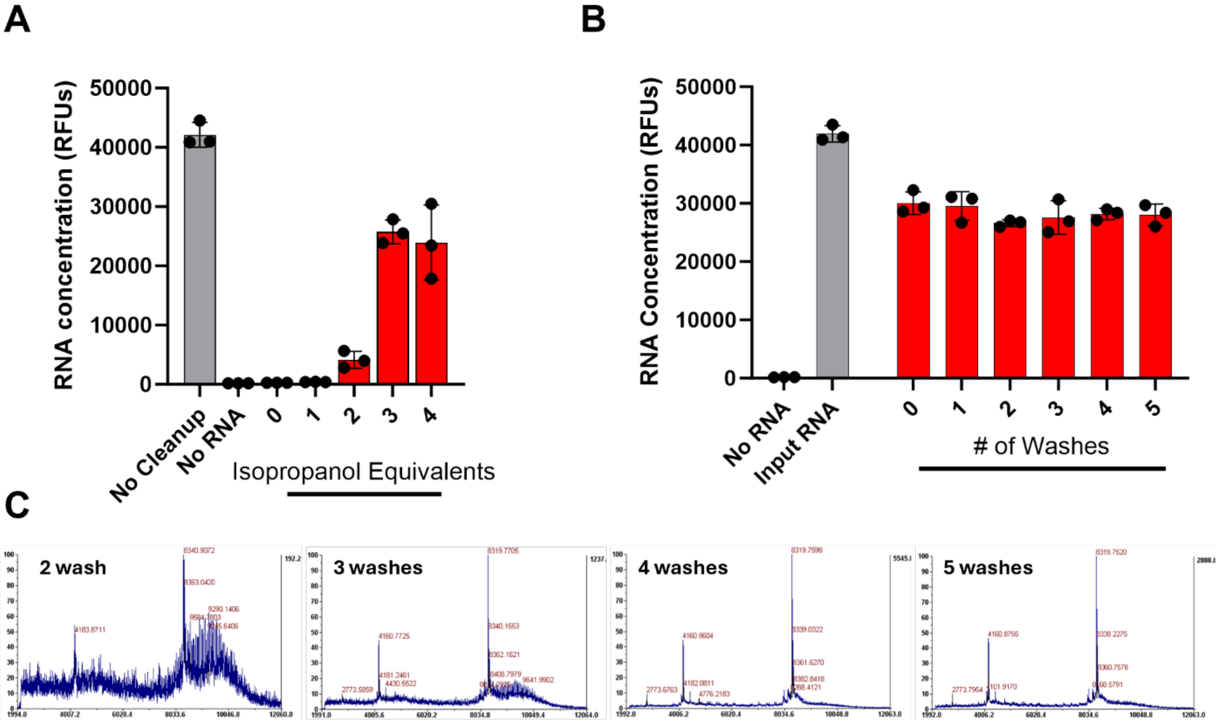


**Figure S3**. **Optimization of bead clean-up of RNA for MALDI-TOF analysis**. **A)** For small RNAs such as **RNA1**, isopropanol (at least three volume equivalents) was required for efficient precipitation of RNA onto the beads. RNA concentration was measured using the Quant-it RiboGreen assay (ThermoFisher), and the fluorescence in the assay is plotted (n = 3 independent measurements). **B)** Sequential washes with 85% (v/v) ethanol did not reduce the recovery of RNA. RNA yield was quantified and plotted as in **(A)** (n = 3 independent measurements). **C)** For MALDI-TOF spectra with high signal-to-noise, at least 4 washes were required. No measurable signal was obtained for samples with one or fewer washes (n = 1 independent measurements).

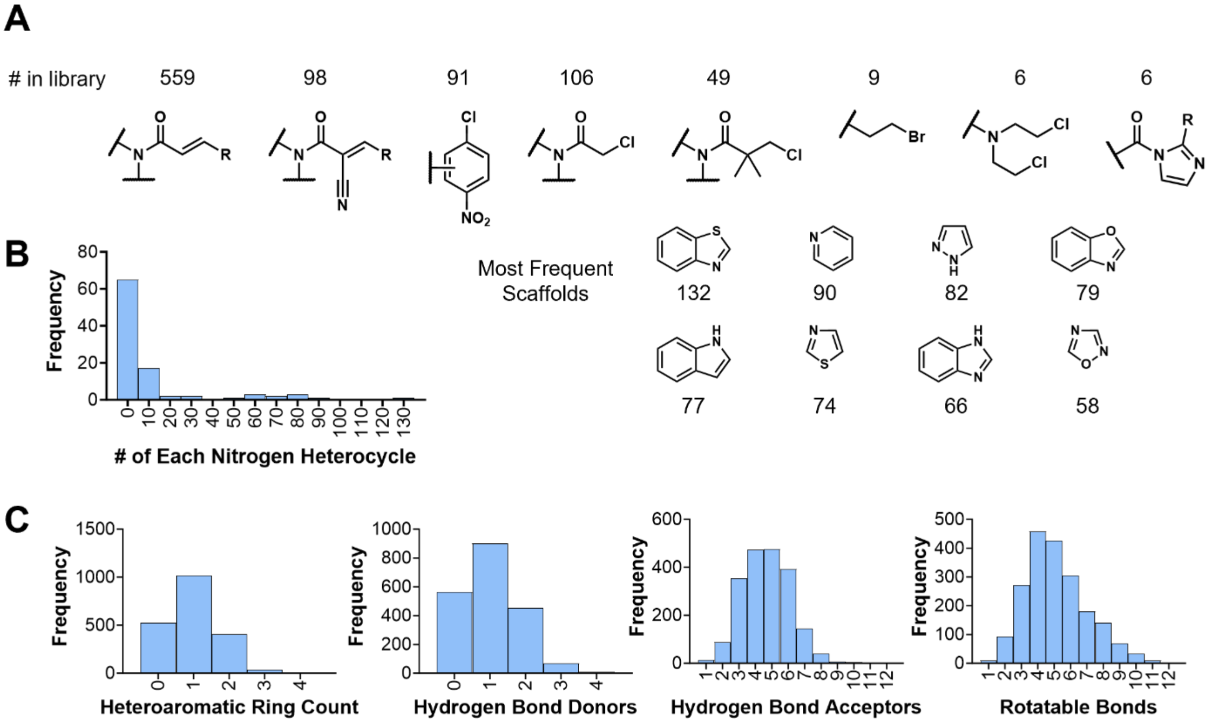


**Figure S4**. **Structural analysis of the 2,000-member electrophile library**. **A)** Electrophiles in the library were largely composed of known cysteine-reactive electrophiles such as acrylamides, cyanoacrylamides, chloronitrobenzenes, and chloroacetamides. Other known nucleic acid-targeting electrophiles such as alkyl bromides, nitrogen mustards, and *N*-acylimidazoles were included as well. **B)** OpenMolecules DataWarrior[2] was used to extract ring system scaffolds present within the library. Filtering for nitrogen-containing aromatic ring systems (common among RNA-binding small molecules) revealed 98 unique scaffolds. A histogram for the frequency of each scaffold (*Left*) and a selection of the eight most common scaffolds (*Right*) are shown. **C)** A selection of physicochemical property descriptors, calculated by RDKit,[3] for the electrophile library are shown, highlighting those potentially impacting RNA-binding, such as heteroaromatic rings, hydrogen bond donors and acceptors, and rotatable bonds.

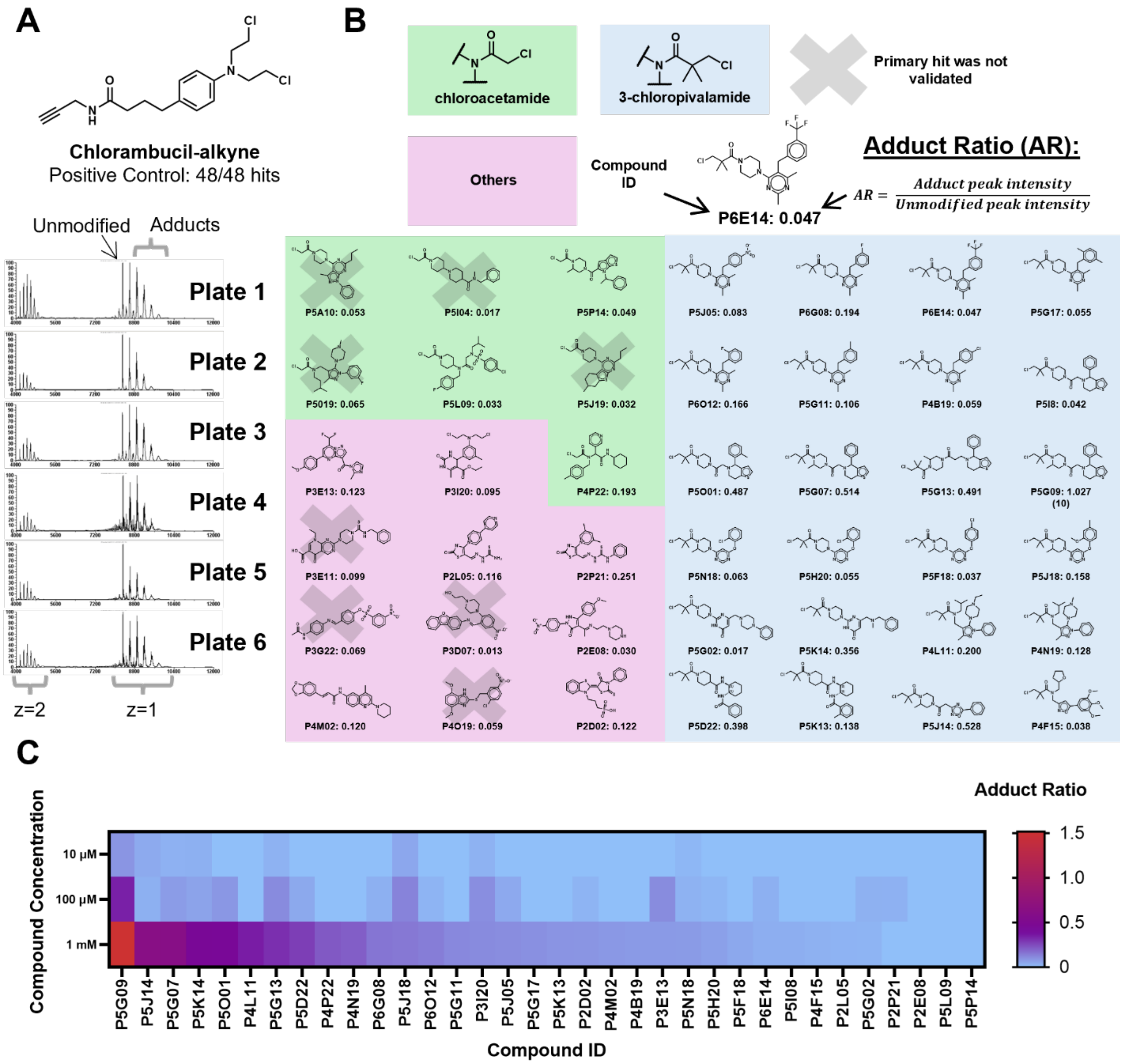


**Figure S5. Screening and hit validation of the 2,000-member electrophile library**. **A)** As a positive control for the assay, chlorambucil-alkyne (2 mM) was used. Eight replicates were included on each plate, and one example spectrum for each is shown. All 48 replicates of chlorambucil-alkyne were hits in the assay, demonstrating a low false-negative rate. **B)** Structures of the 42 primary hit compounds are shown, grouped by the electrophile that they contain. The adduct ratio (calculated as shown) is an indication of the extent of reaction with 20 μM **RNA1**. Structures with a shaded "X" were not able to be replicated during hit validation. Notably, all 24 hits containing the 3-chloropivalamide electrophile were replicable. **C)** The 34 validated, replicable hits were tested in dose response (n = 1) at 1, 0.1, and 0.01 mM, 18 of which showed detectable adduct formation at multiple doses.

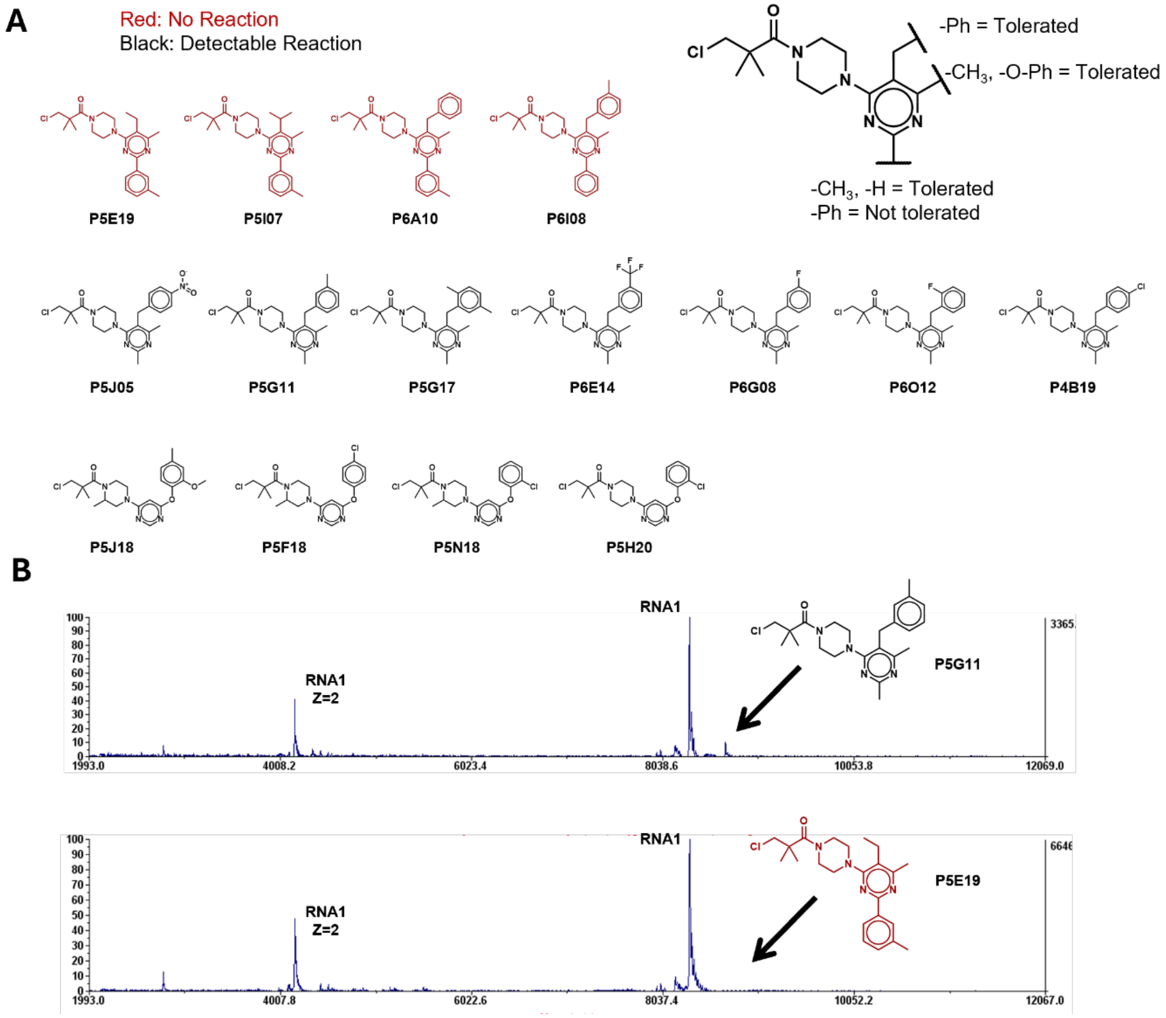


**Figure S6. Structure-activity relationships (SAR) for a common binding scaffold which contains the 3-chloropivalamide electrophile**. **A)** Structures of 14 compounds which contain a central 4-(piperazin-1-yl)pyrimidine motif and their corresponding reactivity at 2 mM compound concentration toward 20 μM **RNA1** in the primary screen. When hits (black) and non-hits (red) are compared, bulky aromatic substituents at the 2-position of the pyrimidine ring abolish reactivity toward **RNA1**. **B)** Two example spectra from the primary screen with **RNA1** show the reactivity (*top*) and lack of reactivity (*bottom*) for two example compounds from this set.

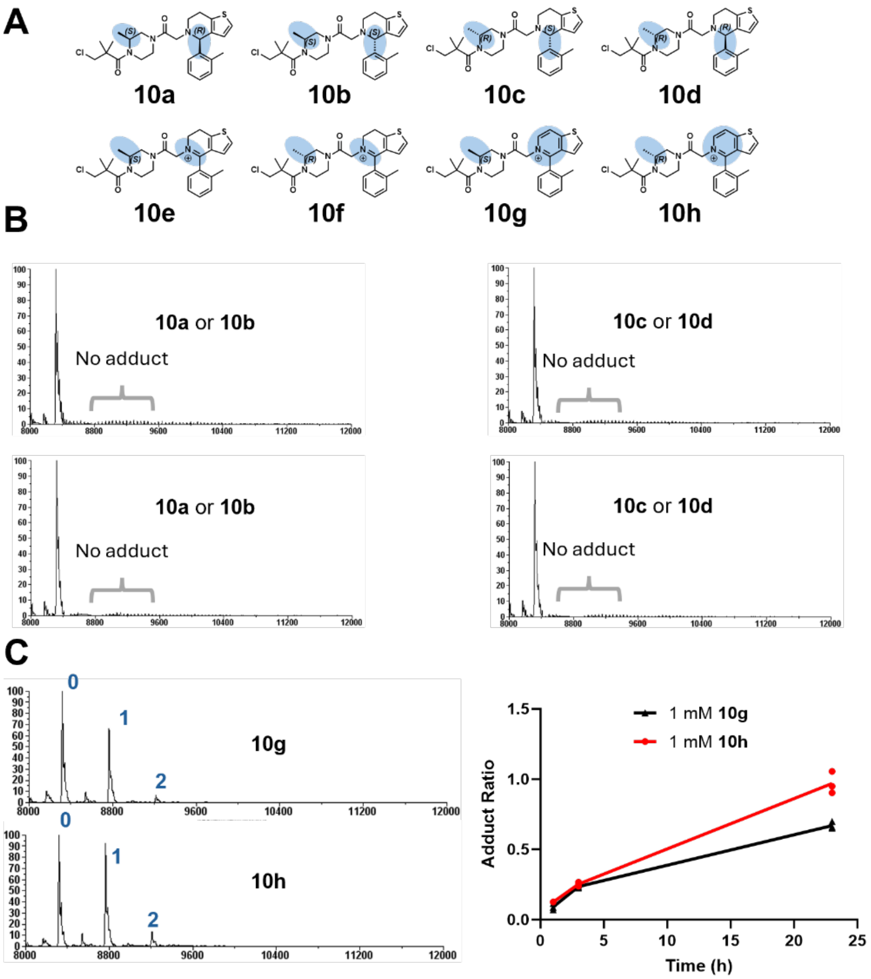


**Figure S7**. **The oxidized derivatives 10g and 10h contribute to the activity of the originally identified hit 10**. **A)** LC/MS analysis of the library stock of **10** revealed it was composed of a mixture of **10a-h**. To determine the active components, **10a-d** and **10g,h** were synthesized enantiomerically (See Synthetic Methods). Compounds **10e** and **10f** were not successfully synthesized due to oxidation to **10g** and **10h** during synthesis. **B)** All diastereomers of the intended hit compound **10** (**10a-d**) were inactive (showed no covalent adducts). **C)** Both oxidized forms **10g** and **10h** (1 mM) resulted in covalent adducts to **RNA1** (20 μM), with a slightly higher extent of reaction for the (*R*) enantiomer **10h**. The extent of adduct formation increased over an extended incubation (~24 h) at 37 °C (n = 3 technical replicates).

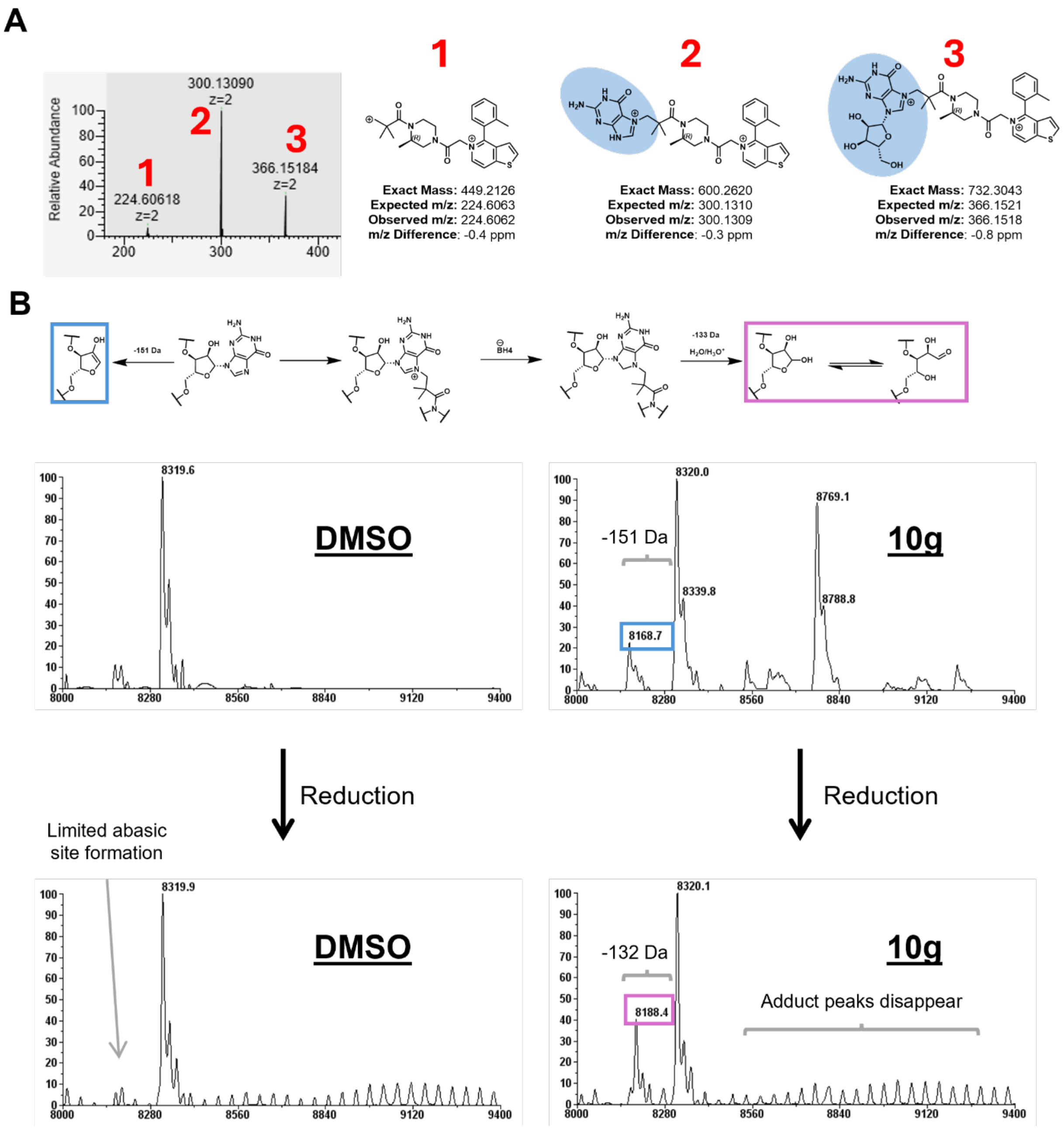


**Figure S8. Site of reactivity identified for the 3-chloropivalamide electrophile**. **A) RNA1** (20 μM) that was covalently modified with **10h** (1 mM) was enzymatically digested to nucleosides (see Methods). The resulting digests were analyzed by high-resolution LC-MS, which revealed that the adducts of **10h** with **RNA1** occur on a guanine base, as adducts to both guanine and guanosine were observed. **B)** To determine whether the site of reaction was at the N7 position of guanine, **RNA1** (20 μM) modified with **10g** (1 mM) was reduced with sodium borohydride (1 M), which will selectively reduce the C8-N7 double bond of N7-alkylated RNA. Following reduction, MALDI-TOF analysis revealed the loss of covalent adducts to **RNA1** and the appearance of a -132 Da peak, as would be expected for depurination and abasic site generation induced by reduction of N7-alkylated guanine.

| Electrophile | # Validated Hits | # in Library | Hit Rate | Example Molecule |
|---|---|---|---|---|
| 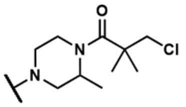 | 7 | 7 | 100% | 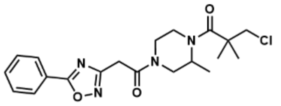 |
| 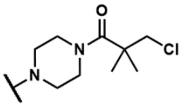 | 12 | 19 | 63% | 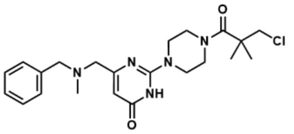 |
| 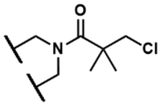 | 3 | 6 | 50% | 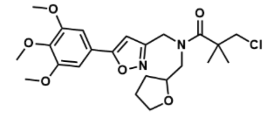 |
| 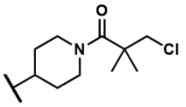 | 2 | 4 | 50% | 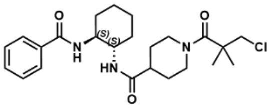 |
| 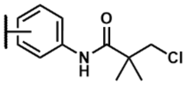 | 0 | 8 | 0% | 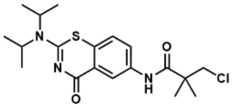 |
| 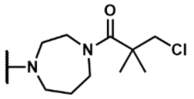 | 0 | 3 | 0% | 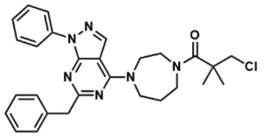 |
| 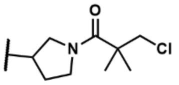 | 0 | 1 | 0% | 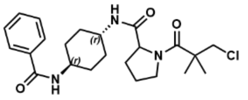 |
| 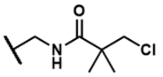 | 0 | 1 | 0% | 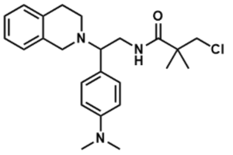 |

**Figure S9. Influence of amide bond properties on the reactivity of 3-chloropivalamide electrophiles**. The hit rate of compounds containing the 3-chloropivalamide electrophile in the screen against **RNA1** appeared to depend upon the amide substituent to which the electrophile was attached. Compounds containing piperazine or piperidine amides had greater than a 50% hit rate in the assay, whereas compounds with primary amines had a 0% hit rate. This suggested the amide bond may play a role in the reactivity of the 3-chloropivalamides, as has been observed for chloroacetamides.[4]

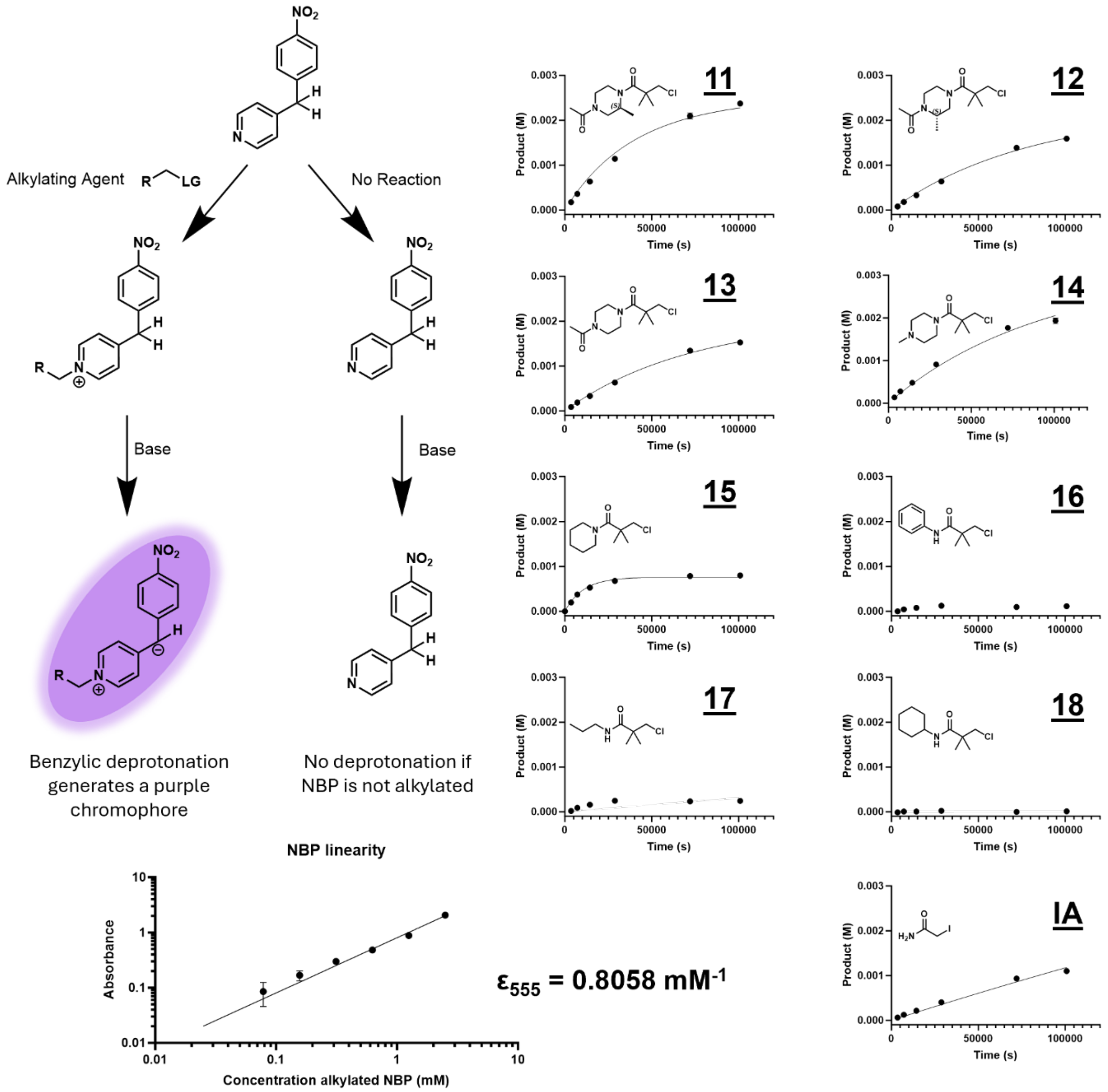


**Figure S10. Reactivity of electrophiles towards guanosine-like nucleophiles**. Alkylation of NBP results in base-mediated deprotonation of its benzylic protons, generating a purple chromophore.[5] The linearity of the assay (*Bottom Left*) was determined using alkylation of NBP with dimethyl sulfate, which completely alkylated the NBP as observed by LC/MS, from which an extinction coefficient was derived (n = 3 technical replicates). The assay was linear below 2 mM of alkylated NBP. The calculated extinction coefficient was used to determine molar yields of alkylated product upon reaction of 10 mM of NBP with 10 mM of **11-18** (and iodoacetamide) and fit to a curve (*Right*), which incorporates the rate of hydrolysis (Figure S11; see Methods).

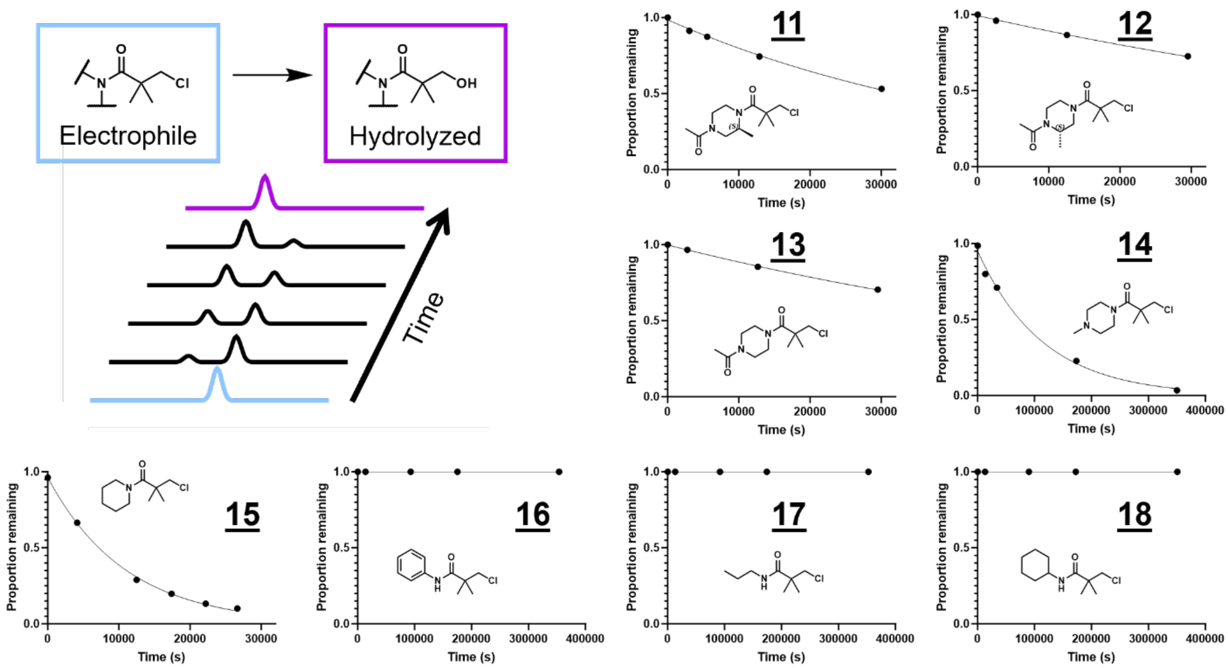


**Figure S11**. **Hydrolysis of 3-chloropivalamides**. Hydrolysis rates were determined by incubating **11-18** at 37 °C and monitoring hydrolysis by LC/MS (n = 1 replicate). Half-lives were determined by fitting to a first-order decay equation (see Methods). No hydrolysis was observed for **16-18**, thus hydrolysis rates could not be determined.

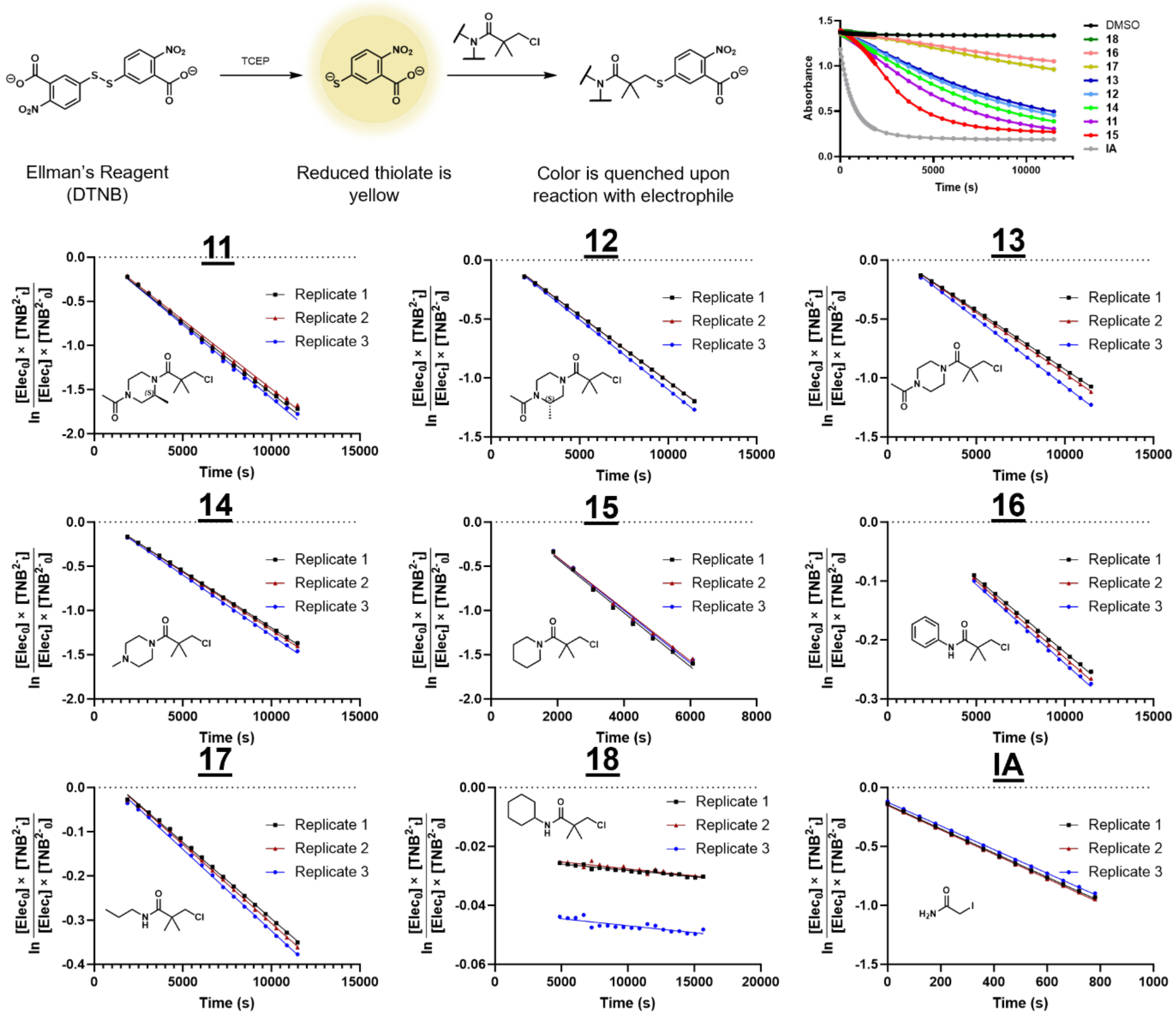


**Figure S12**. **Assessing thiol reactivity of the 3-chloropivalamides using Ellman's reagent (DTNB)**. Reduction of 5,5-dithiobis-(2-nitrobenzoic acid) (DTNB or Ellman's reagent) results in a yellow thiolate ($TNB^{2-}$).[4] Reaction of this thiolate with electrophiles quenches the absorbance, which was monitored over time. Reactions with 100 µM of $TNB^{2-}$ and 5 mM of **11-18,** or 0.5 mM iodoacetamide (IA), were carried out at 37 °C (n = 3 technical replicates). Absorbance was monitored over time (top right). Second-order rate constants for the reactions were determined from the slope of the lines plotted above (see Methods).

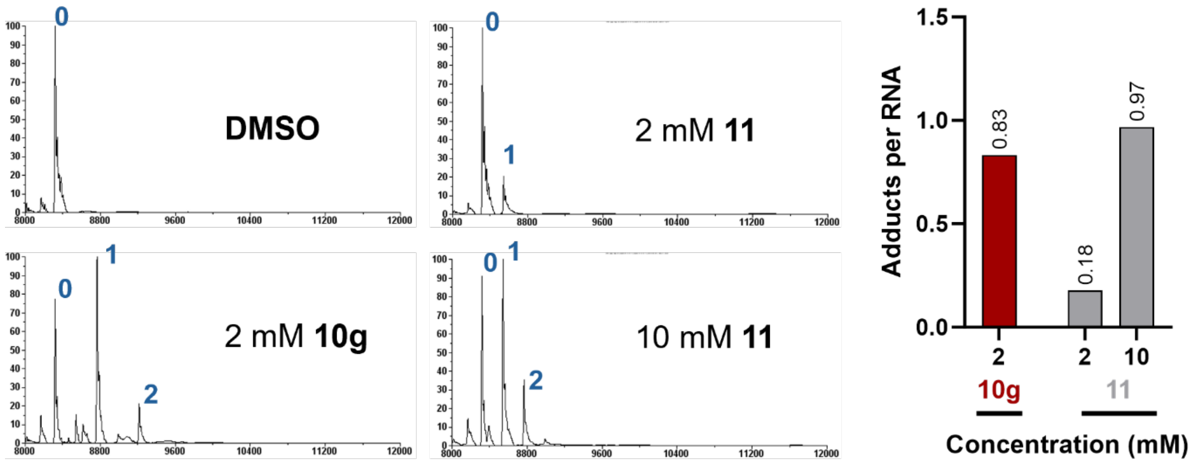


**Figure S13. Influence of the RNA-binding moiety on reactivity**. MALDI-TOF spectra for the reaction of **10g** or **11** with **RNA1** are shown with peaks labeled according to the number of covalent adducts (left). The compound with the binding element (**10g**) forms approximately five times more adducts per RNA (0.83 vs. 0.18) than the electrophile alone **11**, as quantified by peak height (right) (n=1).

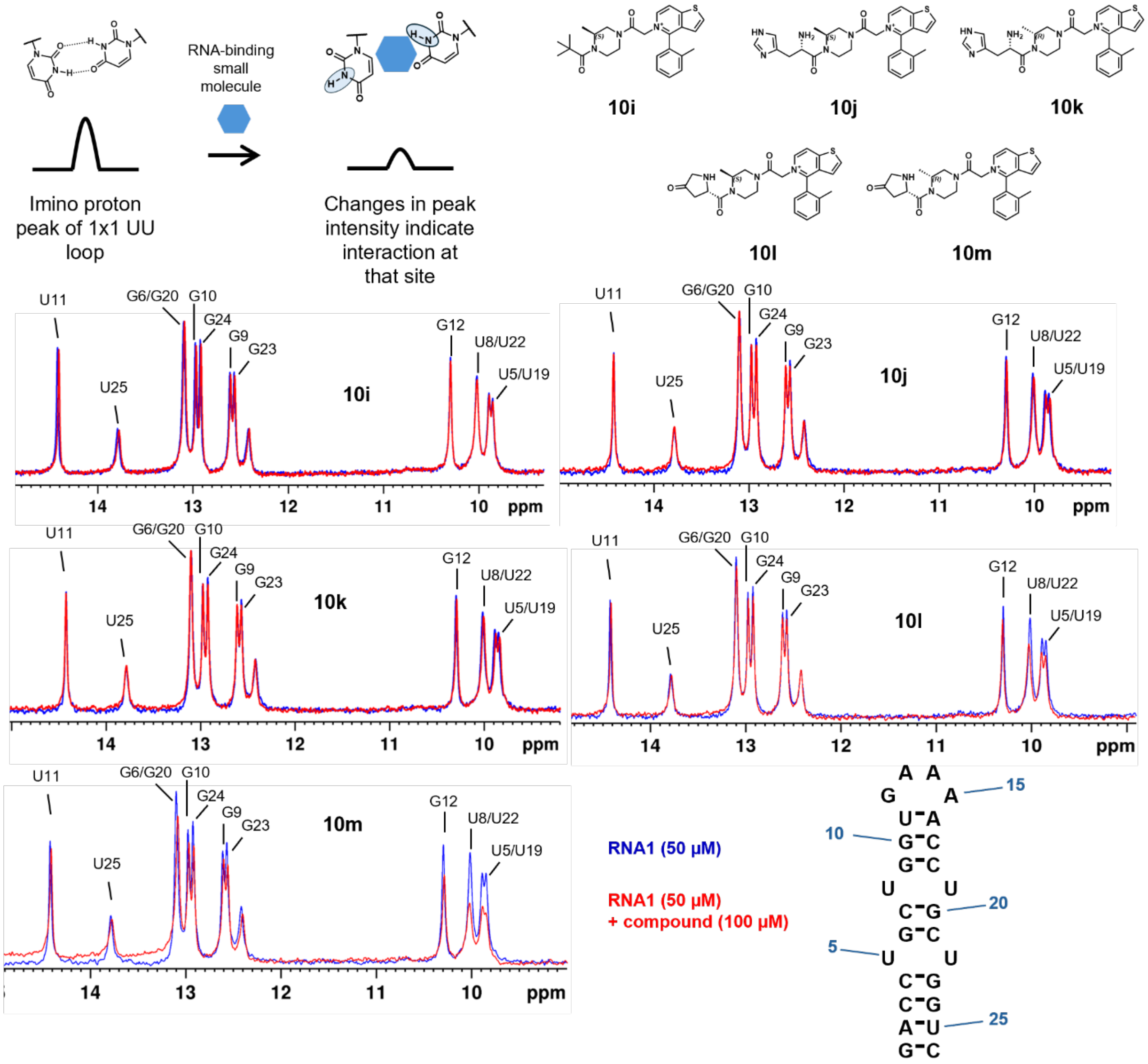


**Figure S14. The site of reactivity of 10g informs more potent r(CUG) repeat binders.** The binding of a non-covalent derivative of **10g**, **10i**, as well as four additional derivatives with the potential to form base-triple interactions with the Hoogsteen face of **RNA1** (**10j-m**) were assessed for binding to **RNA1** by monitoring changes in the RNA's imino proton spectrum by NMR spectrometry. Decreases in the intensity or chemical shift perturbation of imino protons indicate interaction between the compound and the corresponding nucleotides in the RNA. Compounds **10m** and **10l** show the most significant interaction, as assessed by the extent of decrease in intensity of the protons corresponding to the UU 1×1 nucleotide internal loops (U5, U8, U19, and U22).

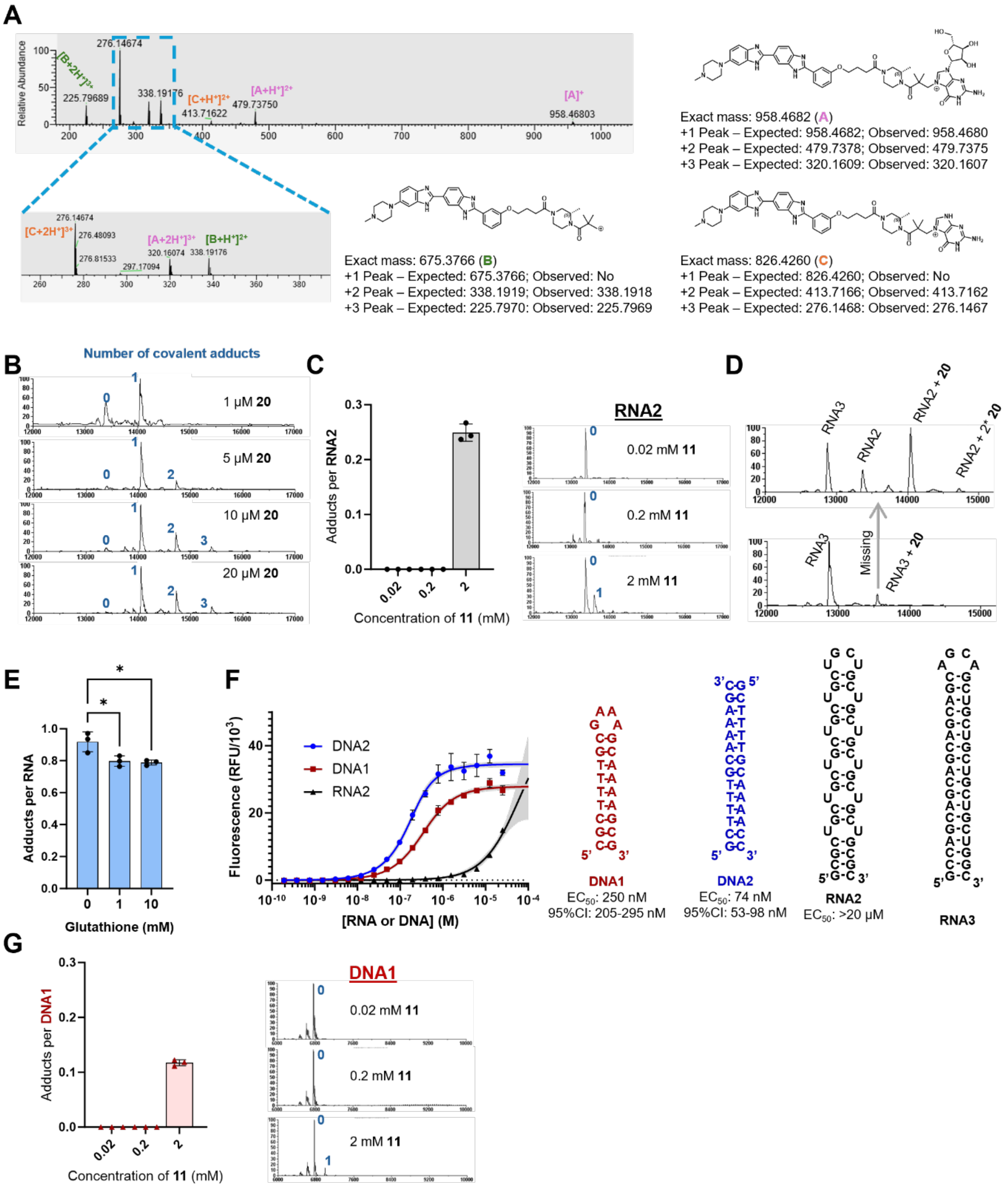


**Figure S15**. **Compound 20 selectively reacts with r(CUG) RNA *in vitro*. A) RNA2** modified with **20** was digested to nucleosides using an enzyme mix. Covalent adducts to both the guanine base and guanosine were observed. Structures were drawn with reaction at the N7 position, although this could not be identified unequivocally from the MS data. **B)** Representative MALDI-TOF spectra of **RNA2** (1 μM) incubated with **20** (1-20 μM) for 24 h at 37 °C. Peaks are labeled with the stoichiometry of covalent adducts that the peak represents. Quantification is found in Figure 6. **C)** *Left:* The electrophile lacking an RNA-binding component (**11**) shows no reaction with **RNA2** at 0.02 or 0.2 mM and low reactivity (< 25% modified) at 2 mM. *Right*: Representative MALDI-TOF spectra for each concentration. **D)** MALDI-TOF spectra show the selectivity of **20** for **RNA2** relative to **RNA3**, the base-paired control. When 20 μM of **20** was co-incubated with 20 μM of both **RNA2** and **RNA3**, no modification of **RNA3** was observed (n = 1) (*Top*). When, **20** was incubated with **RNA3** only, minimal adduct formation was observed (n = 1) (*Bottom*). Peaks are labeled with the RNA construct and/or adduct observed. **E)** Glutathione (1 or 10 mM) marginally reduces adduct formation between **20** (20 μM) and **RNA2** (20 μM). The extent of adduct formation (Adducts per RNA) was quantified using the peak height in the MALDI-TOF spectra (n = 3 technical replicates). **F)** The binding of **19** to model DNA constructs (**DNA1** and **DNA2**) which contain A/T stretches, preferred Hoechst recognition elements, as well as the r(CUG) repeat-containing RNA (**RNA2**) was monitored by the changes in the intrinsic fluorescence of **19** (n = 3 technical replicates). Gray shading shows the 95% confidence interval for the curve fitting. **G)** *Left:* The electrophile lacking an RNA-binding component (**11**) shows no reaction with **DNA1** at 0.02 or 0.2 mM and low reactivity (<15% modified) at 2 mM. *Right*: Representative MALDI-TOF spectra for each concentration.

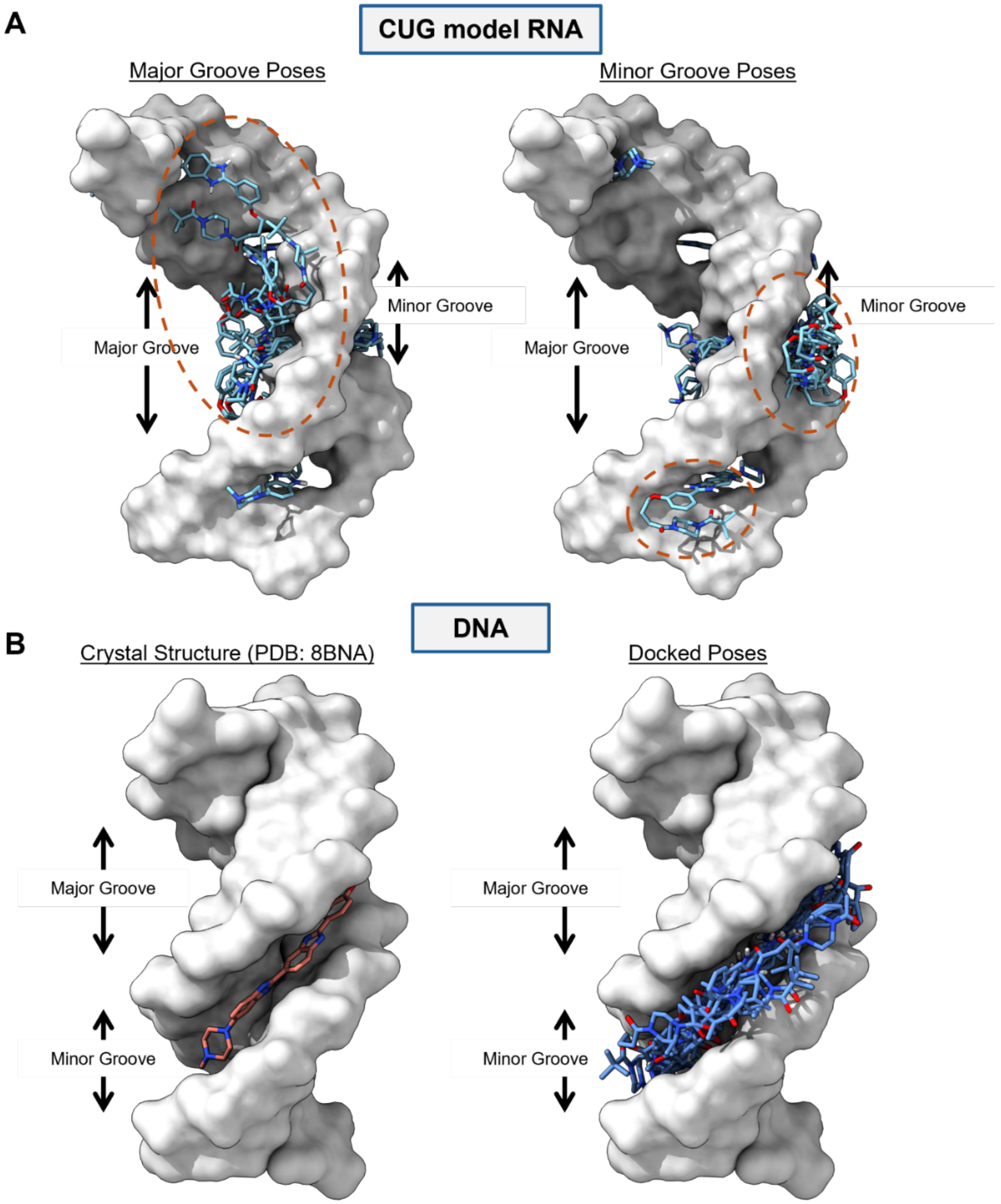


**Figure S16**. **Molecular docking supports that 19 binds differently to DNA than RNA, enabling reactivity in the RNA major groove**. **A)** Compound **19** was docked into a model structure of r(CUG)$^{exp}$ using AutoDock-GPU.[6] Ten of the docked poses, including 6 of the 8 lowest free energy structures, position the electrophile attachment site in the major groove (*Left*), which contains the proposed reactive site, while the other ten docked poses position the electrophile in the minor groove (*Right*). Locations of the electrophile attachment sites are circled in orange. **B)** *Left*: A previous crystal structure of Hoechst 33258 bound to a DNA (PDB: 8BNA)[7] showed binding within the minor groove. *Right*: Molecular docking of **19** into this DNA structure resulted in all 20 bound poses within the DNA minor groove, preventing presentation of the electrophilic site within the DNA major groove.

**Table S1.** Rate constants for reactions involving eight 3-chloropivalamide electrophiles (**11-18**) and iodoacetamide reacting with three nucleophiles were determined (see Methods). The ratio of NBP to DTNB rate constants is more than 200-fold higher for cyclic tertiary forms of the 3-chloro

| Structure | Compound | NBP Reactivity ($M^{-1}s^{-1}$) | Hydrolysis Rate ($s^{-1}$) | DTNB Reactivity ($M^{-1}s^{-1}$) | NBP/DTNB (Relative to IA) |
|---|---|---|---|---|---|
| 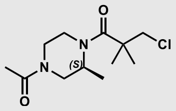 | **11** | $6.4 \pm 0.2 \times 10^{-4}$ | $2.1 \pm 0.3 \times 10^{-5}$ | $3.3 \pm 0.1 \times 10^{-2}$ | 393 |
| 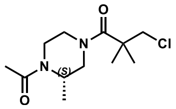 | **12** | $2.87 \pm 0.07 \times 10^{-4}$ | $1.1 \pm 0.2 \times 10^{-5}$ | $2.3 \pm 0.1 \times 10^{-2}$ | 252 |
| 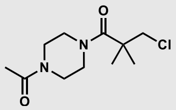 | **13** | $2.86 \pm 0.08 \times 10^{-4}$ | $1.2 \pm 0.1 \times 10^{-5}$ | $2.2 \pm 0.2 \times 10^{-2}$ | 265 |
| 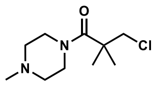 | **14** | $3.4 \pm 0.1 \times 10^{-4}$ | $8 \pm 4 \times 10^{-6}$ | $2.7 \pm 0.1 \times 10^{-2}$ | 260 |
| 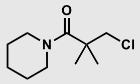 | **15** | $7.2 \pm 0.2 \times 10^{-4}$ | $9.1 \pm 0.6 \times 10^{-5}$ | $6.1 \pm 0.1 \times 10^{-2}$ | 241 |
| 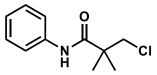 | **16** | $< 1 \times 10^{-4}$ | $< 1 \times 10^{-6}$ | $4.2 \pm 0.1 \times 10^{-3}$ | n.d. |
| 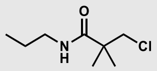 | **17** | $< 1 \times 10^{-4}$ | $< 1 \times 10^{-6}$ | $7.3 \pm 0.2 \times 10^{-3}$ | n.d. |
| 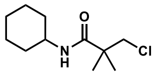 | **18** | $< 1 \times 10^{-4}$ | $< 1 \times 10^{-6}$ | $< 1 \times 10^{-3}$ | n.d. |
| 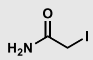 | **IA** | $1.25 \pm 0.06 \times 10^{-4}$ | $< 1 \times 10^{-6}$ | $2.55 \pm 0.04$ | 1 |

**Table S2.** Sequence of oligonucleotides used in this study. All RNA constructs were custom ordered from Horizon Discovery, while DNA constructs were ordered from Integrated DNA Technologies.

| Oligonucleotide Name | Sequence |
|---|---|
| RNA1 | 5’-GACCUGCUGGUGAAAACCUGCUGGUC-3’ |
| RNA2 | 5’-GCGCUGCUGCUGCUGCUGCUGCUGCUGCUGCUGCUGCUGCGC-3’ |
| RNA3 | 5’-GCCAGCAGCAGCAGCAGCAGCAGCUGCUGCUGCUGCUGGC-3’ |
| DNA1 | 5’-CGCTTTTGCGAAAGCAAAAGCG-3’ |
| DNA2 | 5’-GCTTTTGCAAAAGC-3’ |

### Methods

*Oligonucleotides.* RNAs were acquired as deprotected, desalted, and HPLC-purified oligonucleotides from Horizon Discovery. DNA oligonucleotides were ordered using standard desalting purification from Integrated DNA Technologies. Upon arrival, oligonucleotides were dissolved in Nanopure water, and their concentration determined by measuring the absorbance at 260 nm at room temperature in a Beckman Coulter DU800 spectrophotometer. Concentration was calculated using the extinction coefficient provided by Horizon Discovery or Integrated DNA Technologies.

*Pilot Screening of Nitrogen Mustards.* **RNA1** (11.1 µM) was folded in 1.11× Folding Buffer (1× is 12 mM $Na_2HPO_4$, 8 mM $NaH_2PO_4$, pH 7.0, and 0.2 mM EDTA,) by heating at 95 °C for 2 min and immediately snap cooling on ice for at least 10 min. Next, 27 µL of folded RNA was added to 3 µL of 20 mM compound prepared in DMSO (10% (v/v) final concentration). Solutions were incubated at 37 °C for 4 h. The RNA was cleaned up by ethanol precipitation using 5 µL of 6 M ammonium acetate (not pH adjusted) and 90 µL of pre-chilled 100% (v/v) ethanol. Solutions were precipitated for 1 h at -80 °C and then pelleted by centrifugation at 16,000 × *g* for 15 min. After removal of the supernatant, pellets were resuspended in 36 µL of 1 M ammonium acetate and re-precipitated to remove residual salt with 110 µL of 100% (v/v) ethanol. Precipitation and pelleting were performed the same as before, except pellets were dissolved in 20 µL of LC/MS-grade water. The concentration of RNA was determined using a NanoDrop 2000 Spectrophotometer (Thermo Scientific) using calculated extinction coefficients provided by Horizon Discovery.

*Capillary Electrophoresis for gel shift assay.* An Agilent Fragment Analyzer 5200 was used for capillary electrophoresis to assess adduct formation. RNA samples were diluted to 2 ng/µL in water, and samples were prepared following the DNF-470 Small RNA Kit protocol from the manufacturer. Samples were injected at 8.0 kV for 50 s and separated at 11.5 kV for 45 min using a 55 cm column. Results were analyzed using the Agilent ProSize data analysis software. Exported images were quantified using ImageJ to assess the abundance of covalent adducts. First, rectangles were drawn in each column, spanning from the bottom of the image to the top of the primary two bands observed in the DMSO (vehicle only) sample. All rectangles were aligned with the top of the DMSO rectangle. The signal was quantified using the "Plot Lanes" tool. These values represent the "Unmodified" intensity for each column. Then, a new rectangle was drawn from the top of the "Unmodified" rectangle to the top of each column. These were again quantified using the "Plot Lanes" tool, yielding the "Modified" intensity for each column. The ratio of the "Modified" to "Unmodified" intensity for each column (relative to DMSO) was calculated. Hits were identified as those with a ratio at least 4-fold higher than DMSO.

*Denaturing PAGE for gel shift assay.* A 20% (w/v) acrylamide denaturing gel (18.5 × 22 × 0.2 cm, 7.5 M urea, 18:1 acrylamide:bisacrylamide, 1× TBE (89 mM Tris-Borate, pH ~8.3, and 2 mM EDTA)) was pre-run at 250 V in 1× TBE buffer for 1.5 h. RNA samples were diluted to 10 ng/µL with loading dye (8 M Urea, 20 mM EDTA, 2 mM Tris, pH 7.4, and 0.02% (w/v) Orange-G dye), and 2 µL of each solution was loaded into the denaturing polyacrylamide gel. The gel was run for ~2 h at 250 V. Following electrophoretic separation, the gel was incubated with SYBR Gold dye

(1× final concentration; Invitrogen: S11494) in 1× TBE for 15 min, then washed in 1× TBE for 15 min. The gel was imaged using an Azure Sapphire Biomolecular Imager. Gels were imaged using an excitation wavelength of 488 nm and an emission wavelength of 518 nm with a 22 nm bandpass filter.

*MALDI-TOF analysis of RNA samples*.

A 1 µL aliquot of RNA was spotted onto a stainless steel MALDI plate, and the sample was allowed to dry in the air at room temperature. Then, 0.5 µL of THAP matrix (18 mg/mL 2,4,6-trihydroxyacetophenone and 50 mg/mL diammonium citrate in 55:45 water:acetonitrile) was added to each spot, and the matrix was allowed to crystallize at room temperature. Mass spectra were acquired by an Applied Biosystems 4800 plus MALDI-TOF mass spectrometer, using a 355 nm laser set to an intensity of 6800-7200, accumulating sub-spectra with a signal-to-noise greater than 15. Spectra were analyzed in Applied Biosystems/SciEX Data Explorer software.

*Chemical Space Mapping using UMAP*. To visualize the chemical space of the molecular dataset, Uniform Manifold Approximation and Projection (UMAP),[8] a non-linear dimensionality reduction technique, was employed. Molecular representations were first converted into numerical feature vectors using molecular descriptors calculated with Mordred.[9] UMAP was then applied to reduce the high-dimensional descriptor space to two dimensions while preserving local and global structure.

*Calculation of Physicochemical Properties using RDKit*. Physicochemical properties of the molecules were computed using RDKit,[3] an open-source cheminformatics toolkit. The molecular structures were first standardized by removing salts and neutralizing charges. Key properties, including molecular weight, LogP (octanol-water partition coefficient), topological polar surface area (TPSA), hydrogen bond donors and acceptors, rotatable bonds, and ring counts were calculated. These descriptors were used to assess drug-likeness and structural diversity across the dataset.

*MALDI-TOF screen to identify RNA-reactive electrophiles*. A 1900 µL aliquot of **RNA1** (Horizon Discovery, Table S2) at a concentration of 22.2 µM was folded in 1.11× Folding Buffer by heating at 95 °C for 2 min and then immediately snap cooling on ice for at least 10 min. Next, 4.5 µL of folded RNA was added into a 384-well PCR plate (Applied Biosystems REF: 4483285). The plate was centrifuged at 100 × *g* for 20 s, and then 0.5 µL of each compound (10 mM stock in DMSO; 10% (v/v) final concentration) was added. The plate was then covered with an aluminum plate cover (Axygen PCR-AS-600), centrifuged at 100 × *g* for 20 s, and incubated at 37 °C for 12-13 h.

Following the incubation period, 9 µL of RNAClean XP beads (Beckman Coulter: A63987) were added to each well. Then, 20 µL of 100% (v/v) isopropanol was added to facilitate precipitation of the small RNA construct onto the magnetic beads, carefully ensuring each well was thoroughly mixed. After incubating for 5 min at room temperature, the beads were pelleted using a magnetic rack (Permagen P384). The supernatant was removed, and the beads were washed four times with 85% (v/v) ethanol. Following the last wash, the plate was removed from the magnetic rack, and 5 µL of LC/MS grade water was added to each well to elute the RNA from the beads. After incubating at room temperature for 5 min, the plate was placed back onto the magnetic rack to pellet the beads. A 1 µL aliquot of this elution was spotted onto a stainless steel MALDI plate, and the sample was allowed to dry in the air at room temperature. THAP matrix was spotted on top of the samples, which were analyzed as above ("*MALDI-TOF analysis of RNA samples"*).

Hits were defined as compounds that showed a peak with a mass larger than the parent mass of the RNA (~8320 Da) with an intensity of at least 1% of the unmodified RNA. On each plate were eight negative controls (DMSO only) and eight positive controls (2 mM chlorambucil-alkyne; Figure S5). Identified hits were validated by replicating the assay using 45 µL of **RNA1** and 5 µL of each hit compound at final concentrations of 10 µM and 1 mM, respectively, in 1× Folding Buffer. After a 12 h incubation at 37 °C, samples were cleaned up for mass spectrometry using two rounds of ethanol precipitation (following procedure for "*Pilot Screening of Nitrogen Mustards"*) rather than with the RNAClean XP beads. Samples were analyzed by MALDI-TOF in "*MALDI-TOF analysis of RNA samples*".

*Reaction of 3-chloropivalamide electrophiles with oligonucleotides*. Unless otherwise stated, oligonucleotides (**RNA1-3** or **DNA1-2**) at a concentration of 22.2 µM in 1.11× Folding Buffer were folded by heating at 95 °C for 2 min, then immediately placed on ice for at least 10 min. Then, nine volume equivalents of RNA were added to 1 volume equivalent of compound in DMSO (e.g. 18 µL RNA to 2 µL compound). Reactions were incubated at 37 °C for 24 h. Then, reactions were cleaned up with RNAClean XP beads and analyzed by MALDI-TOF MS as described in *"MALDI-TOF screen to identify RNA-reactive electrophiles*".

*Nucleoside digestion to identify reaction site*. Covalently modified RNA (1 µg) was digested to nucleosides using the New England Biolabs Nucleoside Digestion Mix (Ref: M0649S) at 37 °C for 1 h. A 10 µL aliquot of this mix was diluted with 40 µL of 50:50 water:methanol with 0.1% (v/v) formic acid. Then, 10 µL of this diluted sample injected with a Vanquish HPLC system (Thermo Fisher Scientific), and mass spectra were acquired using positive ESI mode on an Orbitrap Exploris 120 (Thermo Fisher Scientific).

*Reduction of covalent RNA adducts*. **RNA1** (20 µM) was incubated with **10g** (2 mM) or DMSO (10% (v/v)) in 50 µL reactions prepared in 1× Folding Buffer for 13 h at 37 °C. Reactions were cleaned up with RNAClean XP beads (see above) and eluted with 25 µL of Nanopure water. Then, 40 µL of 1 M $NaBH_4$ was added to 10 µL of each eluted RNA solution, followed by incubation at room temperature for 4 h.[10] Samples were cleaned up by ethanol precipitation (twice) and then RNAClean XP beads (see above). Samples were analyzed by MALDI-TOF MS, as above.

*Glutathione competition experiments*. **RNA2** (20 µM) was folded using 1× Folding Buffer (as above), then supplemented with 0, 1, or 10 mM reduced glutathione. Then, **20** (20 µM final concentration; final concentration of 10% (v/v) DMSO) was added. The solutions were incubated for 24 h at 37 °C, then samples were cleaned up with RNAClean XP beads and assessed by MALDI-TOF MS as described above.

*Aqueous stability of electrophiles*. Compounds (1 mM) were incubated at 37 °C in 200 mM HEPES-KOH, pH 7.5, and 15% (v/v) DMSO. At the indicated time points, 20 μL was removed and diluted with 80 μL of 50:50 water:methanol with 0.1% (v/v) formic acid. Samples were immediately injected (10 μL) into an Agilent Infinity II liquid chromatography system coupled with an Agilent 6120 quadrupole LC/MS. Integration of the absorbance at 220 nm, 250 nm (for **16**), or the ion counts (for those compounds with insufficient absorbance) was used to quantify the percentage hydrolyzed at each time point. The resulting data were fit to a first-order decay equation as below:

$$P_{intact} = e^{-k_{hydrolysis}*t}$$

Where $P_{intact}$ is the proportion of the electrophile intact (*i.e.*, not hydrolyzed), $k_{hydrolysis}$ is the first-order rate constant for hydrolysis of the electrophile, and t is time (s). The data were fit using Graphpad Prism.

*NBP reactivity of electrophiles*. Compounds (10 mM) were incubated with NBP (10 mM) at 37 °C in 200 mM HEPES-KOH, pH 7.5, and 15% (v/v) DMSO. At the indicated time points, 10 μL of each solution was removed and placed into a clear 384-well plate (Fisher: 12565506). Then, 90 μL of 50:50 triethylamine:ethanol was added to each well, and the samples were thoroughly mixed by pipetting. The plate was covered with an optically clear plate cover (Applied Biosystems 4311971) and centrifuged at 100 × *g* for 20 s. The absorbance at 555 nm was measured in a TECAN infinite Pro M1000 plate reader after seal removal.

To test the linearity of the assay, NBP (10 mM) was completely alkylated with dimethyl sulfate (1 M) at 95 °C for 20 min in 200 mM HEPES-KOH, pH 7.5, and15% (v/v) DMSO. Next, two-fold dilutions into buffer (200 mM HEPES-KOH, pH 7.5, and 15% (v/v) DMSO) were performed. Samples were plated and measured as above. An extinction coefficient was determined by plotting the absorbance vs. concentration of alkylated NBP.

Because all samples fell in the linear range of the assay (below absorbance of 2), the concentration of alkylated NBP was determined using the calculated extinction coefficient. Due to the extended time points of the assay (over 24 h), some electrophiles had significant hydrolysis, impacting the extent of reaction with NBP. To account for this, an analytical solution for the combined first- and second-order rate equations was used to fit the data:[11]

$$[P] = [NBP]_0 \left[1 - e^{\frac{-k_2}{k_{hydrolysis}}[Elec]_0\left(1-e^{-k_{hydrolysis}t}\right)}\right]$$

Where P is the concentration (M) of alkylated NBP product, $[NBP]_0$ is the initial concentration of NBP, $k_2$ is the second order rate constant for reaction between the electrophile and NBP, $k_{hydrolysis}$ is the pseudo-first order rate constant determined previously (*Aqueous stability of electrophiles*), $[Elec]_0$ is the initial concentration (M) of the electrophile, and t is time (s).

*Thiol reactivity of electrophiles*. An assay using Ellman's reagent (DTNB, 5,5′-dithiobis- (2-nitrobenzoic acid)) was adapted from the London group.[4] Briefly, 55.5 μM of DTNB in 1.11×

Reaction Buffer (1× is 200 mM HEPES-KOH, pH 7.5, and 200 µM TCEP) was incubated at 37 °C for 30 min to equilibrate. During equilibration, 10 µL of 50 mM electrophile (or 5 mM iodoacetamide, since reaction kinetics are too rapid at higher concentrations) prepared in DMSO was added to a clear 384-well plate (Fisher: 12565506). Then, 90 µL of DTNB solution – pre-warmed to 37 °C – was added to each well (final concentration of 100 µM for the reduced $TNB^{2-}$ and 5 mM of each electrophile, or 0.5 mM iodoacetamide) and the plate was covered with an optically clear seal (Applied Biosystems: 4311971). The plate was immediately incubated in a TECAN infinite Pro M1000 plate reader at 37 °C, where absorbance at 412 nm was measured as a function of time.

To determine rate constants, absorbance was converted to concentration of $TNB^{2-}$ by subtracting the background absorbance (0 µM $TNB^{2-}$ - found from the last time point of fully alkylated $TNB^{2-}$ with iodoacetamide) and using the DMSO control as no alkylation (100 µM of $TNB^{2-}$). The concentration of the alkylated product at each time point was found by subtracting the calculated concentration of $TNB^{2-}$ at each time point from the initial concentration of $TNB^{2-}$ (100 µM). The concentration of electrophile at each time point was calculated by subtracting the calculated alkylated product concentration from the initial concentration of electrophile (5 mM for 3-chloropivalamides, 0.5 mM for iodoacetamide). The time in s was plotted on the x-axis, and the $ln\frac{[Elec]_0[TNB^{2-}]_t}{[Elec]_t[TNB^{2-}\}_0}$ on the y-axis, where $[Elec]_0$ and $[TNB]_0$ are the initial concentrations of the electrophile and $TNB^{2-}$, respectively, and $[Elec]_t$ and $[TNB^{2-}]_t$ are the concentrations of electrophile and $TNB^{2-}$ at each time point, respectively. The second order rate constant for reaction with DTNB ($k_{DTNB}$) was determined as below:

$$slope = k_{DTNB} * ([TNB^{2-}]_0 - [Elec]_0)$$

The slope of the linear portion of this graph corresponds to the second order rate constant ($k_{DTNB}$) multiplied by the difference in initial concentrations of $TNB^{2-}$ and the electrophile.

*Imino proton NMR spectra to assess compound binding*. NMR spectra were recorded on a 700MHz Bruker Avance III spectrometer equipped with a cryogenic probe. **RNA1** was dissolved in NMR Buffer (5 mM $KH_2PO_4/K_2HPO_4$, pH 6.0, and 0.25 mM EDTA) or NMR Buffer supplemented with 50 mM NaCl and reannealed by heating to 95 °C for 3 min, and then slowly cooling the sample to room temperature before adding to Shigemi tubes (Shigemi, Inc.). Then, $^1$H 1D NMR of RNA exchangeable (imino) protons were acquired in 5% (v/v) $D_2O$ and 95% $H_2O$ at 9˚C in the absence of compound. Compounds dissolved in $D_6$-DMSO were then added to the RNA sample (0.5% (v/v) DMSO) to achieve final concentrations of 1:2 molar equivalents of **RNA1**: compound (50 µM of RNA and 100 µM of compound). A 1:2 ratio was used since there are two UU internal loops and hence two potential compound binding sites. NMR spectra were processed in Topspin 4.0.6.

*Assessing compound binding by monitoring changes in fluorescence intensity*. The core binding component of **19** is a derivative of the Hoechst 33258 and 33342 dyes, whose fluorescence increases upon binding to oligonucleotides.[12] Oligonucleotides were folded in 1× Binding Buffer (8 mM $Na_2HPO_4/NaH_2PO_4$ pH 7.0, 185 mM NaCl, 1 mM EDTA) by heating to 95 °C for 2 min then snap cooling on ice for at least 10 min. Serial 1:1 dilutions of the oligonucleotides into 1× Binding

Buffer were performed, then 10 μL of these solutions were added to a non-binding 384-well black plate (Corning 4514). Then, 10 μL of a 1 μM solution of **19** in 1× Binding Buffer (0.1% (v/v) DMSO) supplemented with 80 μg/mL bovine serum albumin (BSA) was added to each well and thoroughly mixed. The plate was covered with an opaque seal (Axygen PCR-AS-600) and incubated with shaking (800 rpm) for 30 min at room temperature. The fluorescence (360/480 nm excitation/emission with 20 nm bandwidth) was measured using a TECAN infinite pro M1000 plate reader after seal removal. After subtracting background fluorescence of the compound (no RNA added), Plots were made using GraphPad Prism, fit to the equation below.

$$\frac{Max}{2}(EC_{50} + [\mathbf{19}]_0 + [Oligo]_x - \sqrt{{EC_{50}}^2 + 2EC_{50}([\mathbf{19}]_0 + [Oligo]_x) + ([\mathbf{19}]_0 - [Oligo]_x)^2} = F_x$$

Where *Max* is the maximum fluorescence at plateau, $EC_{50}$ is the concentration at half-maximal fluorescence, $[\mathbf{19}]_0$ is the concentration of compound **19**, $[Oligo]_x$ is the concentration of DNA or RNA at each concentration *x*, and $F_x$ is the fluorescence at each concentration *x*. For the fitting, $[\mathbf{19}]_0$ was constrained so that all curves share the same value. Because binding was non-saturating for **RNA2**, a constraint was placed to limit *Max* to less than 50,000, enabling approximate determination of $EC_{50}$, assuming a similar degree of fluorescence enhancement as observed for binding to **DNA1** and **DNA2**.

*Molecular docking of compounds to RNA*. Chimera[13] was used for preparation of the RNA structure for docking using the Amberff14SB force field parameters. The 3D structures of small molecules were created with OBabel[14] from the corresponding SMILES input file and geometry-optimized with general AMBER force field (GAFF)[15] in 5000 cycles prior to further processing for docking. Polar hydrogen atoms and Gasteiger[16] charges (partial charges of each atom) were then added to the small molecules using Chimera. The protonation state of the compounds was calculated at the physiological pH (7.4) prior to docking with OBabel. This protonation state was further validated with pKa prediction by Graph-Convolutional Neural Network.[17] The Grid file was generated from ligand and receptor pdbqt files, applying prepare_gpf4.py, autogrid4 and prepare_dpf4.py script to prepare the docking parameter file. AutoDock-GPU[18] was then used to dock the ligands against the receptor. Twenty different structures were computed with AutoDock-GPU using the Solis Wets[19] method of search. The location (major or minor groove) of the pivalamide moiety for each of the twenty docked positions was determined manually.

### Synthetic Methods:

**Abbreviations:**

DCM: Dichloromethane

DIPEA: Diisopropyl ethyl amine

DMSO: Dimethyl sulfoxide

DMF: N,N-Dimethylformamide

EtOAc: Ethyl acetate

HATU: Hexafluorophosphate azabenzotriazole tetramethyl uronium

HPLC: High-performance liquid chromatography

Hx: hexanes

LC/MS: Liquid chromatography–mass spectrometry

MeOH: Methanol

$R_f$: Retention Factor

RP HPLC: Reversed-phase high-performance liquid chromatography

rt: room temperature, ~22 °C

TEA: triethylamine

TFA: Trifluoroacetic acid

THF: Tetrahydrofuran

TLC: Thin layer chromatography

**General Synthetic Methods**

$^{1}H$ NMR and $^{13}C$ NMR spectra were collected on a 400 MHz UltraShieldTM or a 600 MHz UltraShieldTM NMR spectrometer (Bruker). Silica gel column chromatography was performed using a Biotage Isolera One flash purification system. Preparative RP HPLC was performed using a Waters 1525 Binary HPLC pump equipped with a Waters 2487 dual absorbance detector system and a Waters Sunfire C18 OBD 5 µm, 19 ×150 mm S-14 column. Unless otherwise stated, a linear gradient with a flow rate of 5 mL/min from 0-100% MeOH in water with 0.1% (v/v) TFA was applied over 60 min. Purity was assessed by analytical HPLC using a Waters Symmetry C18 5 µm, 4.6 × 150 mm column with a flow rate of 1 mL/min and a linear gradient from 0-100% MeOH in water with 0.1% (v/v) TFA applied over 60 min; absorbance was monitored at 220, 254, and/or 345 nm. High-resolution mass spectra were acquired using positive ESI mode on an Orbitrap Exploris 120 (Thermo Fisher Scientific) coupled to a Vanquish HPLC system (Thermo Fisher Scientific).

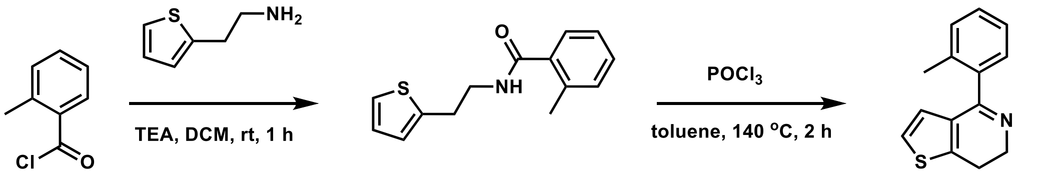


**10-p1** **10-p2**

**Scheme 1.** Synthesis of intermediate **10-p2**.

*Synthesis of 2-Methyl-N-(2-(thiophen-2-yl)ethyl)benzamide (**10-p1**)*

To a stirred solution of 2-methylbenzoyl chloride (CAS: 933-88-0, 5 mL, 41.66 mmol, 1 eq) in DCM (200 mL) were added 2-(thiophen-2-yl)ethan-1-amine (CAS: 30433-91-1, 8.8 mL, 62.49 mmol, 1.5 eq) and TEA (11.5 mL, 83.32 mmol, 2 eq). The reaction mixture was stirred for 1 h at rt. After completeness of reaction was observed by TLC ($R_f$ = 0.3, EtOAc/Hx, 3:7), the solvent was removed under reduced pressure, and the residue was precipitated with water (5 mL) 3 times. The solid was collected and dried under reduced pressure to afford **10-p1** as a white solid (8.17 g, 80% yield).

$^{1}H$ NMR ($CDCl_3$, 400 MHz): δ 7.40-7.22 (m, 5H), 7.06-6.94 (m, 2H), 5.99 (bs, 1H), 3.81-3.70 (m, 2H), 3.24 (t, $J$ = 4.0 Hz, 2H), 2.49 (s, 3H); $^{13}C$ NMR ($CDCl_3$, 100 MHz): δ 170.4, 141.6, 136.7, 136.4, 131.3, 130.1, 127.;4, 127.0, 126.0, 125.8, 124.4, 41.4, 30.3, 20.1; HRMS (ESI) m/z: [M + H]+ for $C_{14}H_{16}NOS^{+}$ calculated 246.0947, found 246.0945.

*Synthesis of 4-(o-Tolyl)-6,7-dihydrothieno[3,2-c]pyridine (**10-p2**)*

To a stirred solution of compound **10-p1** (1.24 g, 5.06 mmol) in toluene (100 mL) was added phosphorus oxychloride (CAS: 10025-87-3, 4.7 mL, 50 mmol, 10 eq). The reaction mixture was stirred for 2 h at 140 °C. After completeness of reaction was observed by TLC ($R_f$ = 0.4, MeOH/DCM, 1:15), the reaction mixture was quenched with $NaHCO_3$ (20 mL) and extracted with DCM (30 mL) 3 times. The organic layer was dried over $MgSO_4$, filtered, and concentrated under reduced pressure. The residue was purified by silica gel column chromatography (EtOAc:Hx, 1:19) to afford **10-p2** as a white solid (990 mg, 86% yield).

$^1$H NMR ($CDCl_3$, 400MHz): δ 7.29-7.25 (m, 2H), 7.24-7.18 (m, 2H), 6.96 (d ,*J* = 5.1 Hz, 1H), 6.61 (d, *J* = 5.2 Hz, 1H), 3.97 (t, *J* = 8.0 Hz, 2H), 2.94 (t, *J* = 8.1 Hz, 2H), 2.20 (s, 3H); $^{13}$C NMR ($CDCl_3$, 100 MHz): δ 163.9, 143.0, 139.2, 135.5, 132.1, 130.4, 128.5, 128.1, 125.68, 125.66, 121.8, 48.3, 22.2, 19.7; HRMS (ESI) m/z: [M + H]+ for $C_{14}H_{14}NS$ calculated 228.0842 found 228.0840.

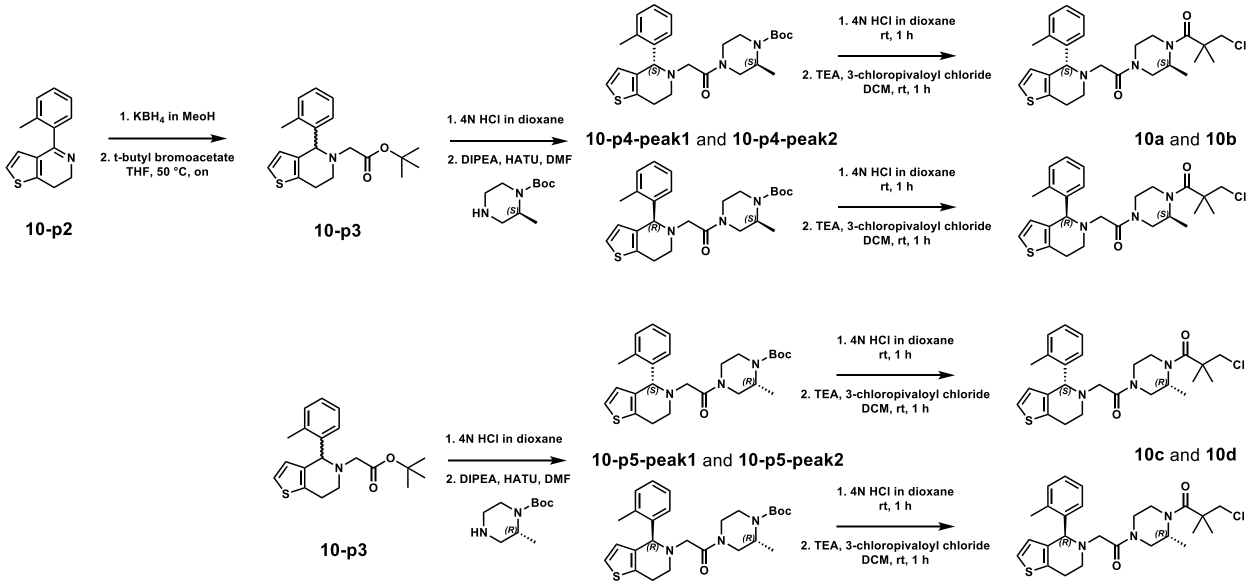


**Scheme 2.** Synthesis of diastereomers **10a-d**.

*Synthesis of tert-butyl 2-(4-(o-tolyl)-6,7-dihydrothieno[3,2-c]pyridin-5(4H)-yl)acetate (**10-p3**)*

To a stirred solution of **10-p2** (700 mg, 3.1 mmol, 1 eq) in anhydrous MeOH was added potassium borohydride (660 mg, 6.2 mmol, 2 eq). The reaction was stirred at rt for 8 h. The solvent was removed under reduced pressure, and the reaction was resuspended in water (40 mL). The product was extracted with DCM (10 mL) 3 times, and the organic layer was dried with sodium sulfate and filtered; solvent was removed under reduced pressure. The crude product was then dissolved in THF (3 mL) followed by addition of potassium carbonate (2.1 g, 15.2 mmol, ~5 eq) and *tert*-butyl 2-bromoacetate (CAS: 5292-43-3, 330 mg, 6.1 mmol, ~2 eq). The reaction was stirred at 50 °C overnight. The solvent was then removed under reduced pressure, and the product was purified using silica gel column chromatography (0-100% DCM in hexanes) to afford **10-p3** (386 mg, 37% yield).

$^{1}H$ NMR ($CDCl_3$, 400MHz): δ 7.27-7.23 (m, 1H), 7.18-7.09 (m, 3H), 6.90 (d, *J* = 5.2 Hz, 1H), 6.20 (d, *J* = 5.2 Hz, 1H), 5.06 (s, 1H), 3.27-3.04 (m, 5H), 2.88-2.78 (m, 1H), 2.29 (s, 3H), 1.43 (s, 9 H); HRMS (ESI) m/z: [M + H]+ for $C_{20}H_{26}NO_2S^+$ calculated 344.1679 found 344.1688.

*Synthesis of tert-Butyl-(S)-2-methyl-4-(2-((R)-4-(o-tolyl)-6,7-dihydrothieno[3,2-c]pyridin-5(4H)-yl)acetyl)-piperazine-1-carboxylate and tert-butyl-(S)-2-methyl-4-(2-((S)-4-(o-tolyl)-6,7-dihydrothieno[3,2-c]pyridin-5(4H)-yl)acetyl)piperazine-1-carboxylate (**10-p4-peak1** and **10-p4-peak2**)*

To a stirred solution of **10-p3** (150 mg, 0.44 mmol, 1 eq) in DCM (1 mL) was added 4 N HCl in dioxane (4 mL). The reaction mixture was stirred at rt overnight, and then the solvent was removed under reduced pressure. The crude solid was resuspended in dry DMF (2 mL) with DIPEA (270 mg, 2.09 mmol, ~5 eq) and HATU (218 mg, 0.57 mmol, 1.3 eq). After stirring at room temperature for 2 min, *tert*-butyl (*S*)-2-methylpiperazine-1-carboxylate (CAS: 169447-70-5, 209 mg, 1.04 mmol, 2.4 eq) was added. The reaction was stirred overnight at room temperature. Solvent was removed under reduced pressure. The reaction mixture was resuspended in DCM (20 mL) and washed with water (20 mL). The organic layer was dried with sodium sulfate, filtered, and solvent was removed under reduced pressure. The products were purified via RP-HPLC (60 min, 40-100% methanol:water + 0.1% (v/v) TFA). The two diastereomer products were purified separately, yielding **10-p4-peak1** (21.6 mg, 21% yield) and **10-p4-peak2** (15.5 mg, 15% yield). Though the two diastereomers were separated successfully, the stereochemistry of the benzylic stereocenter was not assigned.

**10-p4-peak1**: $^{1}H$ NMR ($CDCl_3$, 400MHz): δ 7.35-7.24 (m, 4H, overlap with solvent residual peak), 7.10 (dd, *J* = 5.2, 1.7 Hz, 1H), 6.48 (bs, 0.5H), 6,40 (bs, 0.5H), 6.18 (t, *J* = 5.6 Hz, 1H), 4.46-4.20 (m, 4H), 4.01-3.77 (m, 3H), 3.63-3.53 (m, 1H), 3.45-3.28 (m, 1H), 3.17-2.76 (m, 4H), 2.42 (s, 1.5H), 2.39 (s, 1.5H), 1.45 (s, 9H), 1.06 (d, *J* = 6.8 Hz, 1.5H), 1.04 (d, *J* = 6.8 Hz, 1.5H); HRMS (ESI) m/z: [M + H]+ for $C_{26}H_{36}N_3O_3S^+$ calculated 470.2472 found 470.2467. Analytical HPLC: retention time – 37.4 min.

**10-p4-peak2:** $^{1}H$ NMR ($CDCl_3$, 400MHz): δ 7.36-7.24 (m, 4H, overlap with solvent residual peak), 7.12 (t, *J* = 5.9 Hz, 1H), 6.53 (bs, 0.5H), 6.37 (bs, 0.5H), 6.21 (d, *J* = 5.3 Hz, 0.5H), 6.17 (d, *J* = 5.3 Hz, 0.5H), 4.52-4.26 (m, 3H), 4.18 (d, *J* = 13 Hz, 1H), 4.07-3.78 (m, 3H), 3.67-3.51 (m, 1H), 3.50-2.68 (m, 5H), 2.43 (s, 1.5H), 2.39 (s, 1.5H), 1.45 (s, 9H), 1.02 (d, *J* = 6.7 Hz, 1.5H), 0.75 (d, *J* = 6.8 Hz, 1.5H); HRMS (ESI) m/z: [M + H]+ for $C_{26}H_{36}N_3O_3S^+$ calculated 470.2472 found 470.2467. Analytical HPLC: retention time – 39.5 min.

*Synthesis of tert-Butyl-(R)-2-methyl-4-(2-((S)-4-(o-tolyl)-6,7-dihydrothieno[3,2-c]pyridin-5(4H)-yl)acetyl)-piperazine-1-carboxylate and tert-butyl-(R)-2-methyl-4-(2-((R)-4-(o-tolyl)-6,7-dihydrothieno[3,2-c]pyridin-5(4H)-yl)acetyl)piperazine-1-carboxylate (**10-p5-peak1** and (**10-p5-peak2**)*

To a stirred solution of **10-p3** (150 mg, 0.44 mmol 1 eq) in DCM (1 mL) was added 4 N HCl in dioxane (4 mL). The reaction mixture was stirred at rt overnight, and then the solvent was removed under reduced pressure. The crude solid was resuspended in dry DMF (2 mL) with DIPEA (270 mg, 2.09 mmol, 5 eq) and HATU (218 mg, 0.57 mmol, 1.3 eq). After stirring at room temperature for 2 min, *tert*-butyl (*R*)-2-methylpiperazine-1-carboxylate (CAS: 170033-47-3, 209 mg, 1.04 mmol, 2.4 eq) was added. The reaction was stirred overnight at room temperature, after which the solvent was removed under reduced pressure. The reaction mixture was resuspended in DCM (20 mL) and washed with water (20 mL). The organic layer was dried with sodium sulfate, filtered, and solvent was removed under reduced pressure. The products were purified via RP-HPLC (60 min, 40-100% methanol:water + 0.1% (v/v) TFA). The two diastereomer products were purified separately, yielding **10-p5-peak1** (12.8 mg, 12% yield) and **10-p5-peak2** (14.1 mg, 14% yield). Though the two diastereomers were separated successfully, the stereochemistry of the benzylic stereocenter was not assigned.

**10-p5-peak1**: $^1$H NMR ($CDCl_3$, 400MHz): δ 7.37-7.24 (m, 4H, overlap with solvent residual peak), 7.13 (d, *J* = 5.2 Hz, 1H), 6.48 (bs, 0.5H), 6.41 (bs, 0.5H), 6.20 (t, *J* = 5.4 Hz, 1H), 4.44-3.96 (m, 5H), 3.94-3.72 (m, 2H), 3.70-3.51 (m, 1H), 3.51-2.70 (m, 5H), 2.42 (s, 1.5H), 2.39 (s, 1.5H), 1.45 (s, 9H), 1.07 (d, *J* = 6.8 Hz, 1.5H), 0.97 (d, *J* = 6.7 Hz, 1.5H); HRMS (ESI) m/z: [M + H]+ for $C_{26}H_{36}N_3O_3S^+$ calculated 470.2472 found 470.2469. Analytical HPLC: retention time – 37.6 min.

**10-p5-peak2**: $^1$H NMR ($CDCl_3$, 400MHz): δ 7.37-7.23 (m, 4H, overlap with solvent residual peak), 7.12 (t, *J* = 5.6 Hz, 1H), 6.53 (bs, 0.5H), 6.37 (bs, 0.5H), 6.20 (d, *J* = 5.3 Hz, 0.5H), 6.17 (d, *J* = 5.3 Hz, 0.5H), 4.53-4.26 (m, 3H), 4.18 (d, *J* = 12 Hz, 1H), 4.02-3.74 (m, 3H), 3.68-3.53 (m, 1H), 3.50-2.68 (m, 5H), 2.43 (s, 1.5H), 2.39 (s, 1.5H), 1.45 (s, 9H), 1.03 (d, *J* = 6.7 Hz, 1.5H), 0.74 (d, *J* = 6.7 Hz, 1.5H); HRMS (ESI) m/z: [M + H]+ for $C_{26}H_{36}N_3O_3S^+$ calculated 470.2472 found 470.2468. Analytical HPLC: retention time – 39.6 min.

*Synthesis of 3-Chloro-2,2-dimethyl-1-((S)-2-methyl-4-(2-((R)-4-(o-tolyl)-6,7-dihydrothieno[3,2-c]pyridin-5(4H)-yl)acetyl)piperazin-1-yl)propan-1-one* ***and*** *3-chloro-2,2-dimethyl-1-((S)-2-methyl-4-(2-((S)-4-(o-tolyl)-6,7-dihydrothieno[3,2-c]pyridin-5(4H)-yl)acetyl)piperazin-1-yl)propan-1-one (**10a** and **10b**)*

To a stirred solution of **10-p4-peak1** (22 mg, 47 µmol, 1 eq) in DCM (1 mL), 4 N HCl in dioxane (1 mL) was added. The reaction was stirred for 1 h at rt, and then the solvent was removed under reduced pressure. The crude product was then dissolved in DCM (0.5 mL) followed by addition of triethylamine (22 mg, 220 µmol, ~5 eq) and 3-chloropivaloyl chloride (CAS: 4300-97-4, 8.4 mg, 54 µmol, 1.1 eq). The reaction was stirred for 1 h at rt, and solvent was removed under reduced pressure. The product was purified by RP-HPLC (60 min, 20-100% methanol:water + 0.1% (v/v) TFA), affording a white solid (**10a** or **10b**, 4.6 mg, 20% yield).

$^1$H NMR ($CDCl_3$, 400MHz): δ 7.36-7.23 (m, 4H, overlap with solvent residual peak), 7.12 (d, *J* = 5.2 Hz, 0.5H), 7.09 (d, *J* = 5.2 Hz, 0.5H), 6.42 (bs, 0.5H), 6.30 (bs, 0.5H), 6.20 (d, *J* = 5.3 Hz, 0.5 H), 6.17 (d, *J* = 5.3 Hz, 0.5H), 4.72 (m, 0.5H), 4.64-4.14 (m, 4H), 4.14-3.95 (m, 1H), 3.83-3.50 (m, 5H), 3.46-3.34 (m, 0.5H), 3.33-2.98 (m, 3H), 2.93-2.70 (m, 1H), 2.42 (s, 1.5H), 2.38 (s, 1.5H), 1.38-1.33 (m, 6H), 1.17 (d, *J* = 6.7 Hz, 1.5H), 1.13 (d, *J* = 6.8 Hz, 1.5H); HRMS (ESI) m/z: [M + H]+ for $C_{26}H_{35}ClN_3O_2S^+$ calculated 488.2133 found 488.2132. Analytical HPLC: retention time – 34.0 min.

To a stirred solution of **10-p4-peak2** (15 mg, 32 µmol, 1 eq) in DCM (1 mL), 4 N HCl in dioxane (1 mL) was added. The reaction was stirred for 1 h at rt, and then the solvent was removed under reduced pressure. The crude product was then dissolved in DCM (0.5 mL) followed by addition of triethylamine (22 mg, 220 µmol, ~7 eq) and 3-chloropivaloyl chloride (CAS: 4300-97-4, 6 mg, 38 µmol, 1.2 eq). The reaction was stirred for 1 h at rt, solvent was removed under reduced pressure, and the product was purified by RP-HPLC (60 min, 40-100% methanol:water + 0.1% (v/v) TFA), affording a white solid (**10a** or **10b**, 9.3 mg, 60% yield).

$^1$H NMR ($CDCl_3$, 400MHz): δ 7.36-7.25 (m, 4H, overlap with solvent residual peak), 7.12 (t, *J* = 4.5 Hz, 1H), 6.46 (bs, 0.5H), 6.39 (bs, 0.5H), 6.19 (t, *J* = 5.6 Hz, 1H), 4.79-4.66 (m, 0.5H), 4.66-4.15 (m, 4H), 4.15-3.98 (m, 0.5H), 3.97-3.75 (m, 2H), 3.75-3.52 (m, 4H), 3.39-2.99 (m, 3H), 2.99-2.66 (m, 1H), 2.43 (s, 1.5H), 2.38 (s, 1.5H), 1.38-1.33 (m, 6H), 1.11 (d, *J* = 6.8 Hz, 1.5H), 0.86 (d, *J* = 6.7 Hz, 1.5H); HRMS (ESI) m/z: [M + H]+ for $C_{26}H_{35}ClN_3O_2S^+$ calculated 488.2133 found 488.2131. Analytical HPLC: retention time – 35.6 min.

*Synthesis of 3-Chloro-2,2-dimethyl-1-((R)-2-methyl-4-(2-((S)-4-(o-tolyl)-6,7-dihydrothieno[3,2-c]pyridin-5(4H)-yl)acetyl)piperazin-1-yl)propan-1-one* ***and*** *3-chloro-2,2-dimethyl-1-((R)-2-methyl-4-(2-((R)-4-(o-tolyl)-6,7-dihydrothieno[3,2-c]pyridin-5(4H)-yl)acetyl)piperazin-1-yl)propan-1-one (**10c** and **10d**)*

To a stirred solution of **10-p5-peak1** (12 mg, 26 µmol, 1 eq) in DCM (1 mL), was added 4 N HCl in dioxane (1 mL). The reaction was stirred for 1 h at rt, then the solvent was removed under reduced pressure. The crude product was then dissolved in DCM (0.5 mL) followed by addition of triethylamine (22 mg, 220 µmol, ~8.5 eq) and 3-chloropivaloyl chloride (CAS: 4300-97-4, 6 mg, 38 µmol, 1.5 eq). The reaction was stirred for 1 h at rt, then the product was purified by RP-HPLC (60 min, 20-100% methanol:water + 0.1% (v/v) TFA), affording a white solid (**10c** or **10d**, 3.7 mg, 30% yield).

$^1$H NMR ($CDCl_3$, 400MHz): δ 7.36-7.23 (m, 4H, overlap with solvent residual peak), 7.12 (d, *J* = 5.3 Hz, 0.5H), 7.09 (d, *J* = 5.2 Hz, 0.5H), 6.42 (bs, 0.5H), 6.31 (bs, 0.5H), 6.20 (d, *J* = 5.3 Hz, 0.5 H), 6.18 (d *J* = 5.2 Hz, 0.5H), 4.79-4.66 (m, 0.5H), 4.62-4.14 (m, 4H), 4.14-3.93 (m, 1H), 3.83-3.50 (m, 5H), 3.44-3.35 (m, 0.5H), 3.22-3.02 (m, 3H), 2.94-2.69 (m, 1H), 2.42 (s, 1.5H), 2.38 (s, 1.5H), 1.40-1.32 (m, 6H), 1.17 (d, *J* = 6.6 Hz, 1.5H, 1.13 (d, *J* = 6.8 Hz, 1.5H); HRMS (ESI) m/z: [M + H]+ for $C_{26}H_{35}ClN_3O_2S^+$ calculated 488.2133 found 488.2130. Analytical HPLC: retention time – 34.0 min.

To a stirred solution of **10-p5-peak2** (15 mg, 32 µmol, 1 eq) in DCM (1 mL), 1 mL of 4 N HCl in dioxane was added. The reaction was stirred for 1 h at rt, then the solvent was removed under reduced pressure. The crude product was then dissolved in DCM (0.5 mL) followed by addition of triethylamine (22 mg, 220 µmol, ~7 eq) and 3-chloropivaloyl chloride (CAS: 4300-97-4, 7.2 mg, 47 µmol, 1.5 eq). The reaction was stirred for 1 h at rt, solvent was removed under reduced pressure, and the product was purified by RP-HPLC (60 min, 20-100% methanol:water + 0.1% (v/v) TFA), affording a white solid (**10c** or **10d**, 1.6 mg, 10% yield).

$^1$H NMR ($CDCl_3$, 400MHz): δ 7.36-7.23 (m, 4H, overlap with solvent residual peak), 7.09 (d, *J* = 5.2 Hz, 0.5H), 7.08 (d, *J* = 5.2 Hz, 0.5H), 6.30 (bs, 0.5H), 6.26 (bs, 0.5H), 6.18 (d, *J* = 5.2 Hz, 0.5H), 6.17 (d, *J* = 5.2 Hz, 0.5 H), 4.77-4.42 (m, 2H), 4.42-3.98 (m, 3H), 3.87-3.61 (m, 4H), 3.61-3.29 (m, 2H), 3.29-2.62 (m, 4H), 2.42 (s, 1.5H), 2.36 (s, 1.5H), 1.37-1.33 (m, 6H), 1.14 (d, *J* = 6.7

Hz, 1.5H), 0.81 (d, *J* = 6.7 Hz, 1.5H); HRMS (ESI) m/z: [M + H]+ for $C_{26}H_{35}ClN_3O_2S^+$ calculated 488.2133 found 488.2133. Analytical HPLC: retention time – 35.6 min.

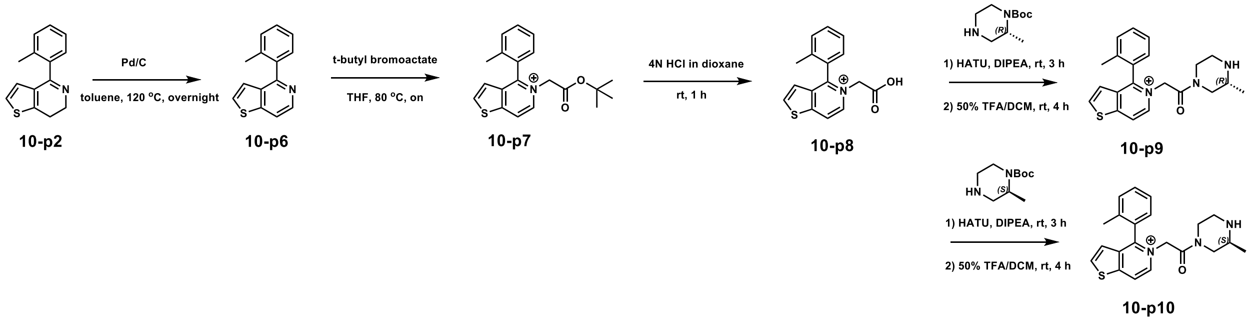

**Scheme 3.** Synthesis of oxidized intermediates **10-p9** and **10-p10**.

*Synthesis of 4-(o-Tolyl)thieno[3,2-c]pyridine (**10-p6**)*

To a stirred solution of compound **10-p2** (500 mg, 2.20 mmol) in toluene (100 mL), was added Pd/C (10 wt%, 500 mg). The reaction mixture was stirred overnight at 120 °C. After completeness of reaction was observed by TLC ($R_f$ = 0.6, MeOH/DCM, 1:15), the reaction mixture was washed with brine (20 mL) and extracted with DCM (20 mL) 3 times. The organic layer was dried over $MgSO_4$, filtered, and concentrated under reduced pressure. The residue was purified by silica gel column chromatography (EtOAc:Hx, 1:12) to afford **10-p6** as a white solid (425 mg, 86% yield).

$^1$H NMR ($CDCl_3$, 400MHz): δ 8.55 (d, *J* = 8.0 Hz, 1H), 7.80 (dd, *J* = 4.0, 0.9 Hz, 1H), 7.43 (d, *J* = 8.0 Hz, 1H), 7.40-7.27 (m, 4H), 7.14 (dd, *J* = 4.0, 0.9 Hz, 1H), 2.19 (s, 3H); $^{13}$C NMR ($CDCl_3$, 100 MHz): δ 156.7, 147.9, 142.4, 139.5, 136.5, 135.2, 130.9, 129.8, 128.9, 127.1, 126.0, 123.9, 116.5, 20.2; HRMS (ESI) m/z: [M + H]+ for $C_{14}H_{12}NS$ calculated 226.0685, found 226.0684.

*Synthesis* of 5-(2-(tert-butoxy)-2-oxoethyl)-4-(o-tolyl)thieno[3,2-c]pyridin-5-ium (**10-p7**)

To a stirred solution of compound **10-p6** (500 mg, 2.22 mmol, 1 eq) in THF (50 mL) was added *tert*-butyl-2-bromoacetate (0.6 mL, 4.44 mmol, 2 eq). The reaction mixture was stirred overnight at 80 °C. After completeness of reaction was observed by TLC ($R_f$ = 0.2, MeOH/DCM, 1:10), the solvent was removed under reduced pressure, and the residue was purified by silica gel column chromatography (EtOAc:Hx, 1:5) to afford compound **10-p7** as a white solid (543 mg, 72% yield).

$^1$H NMR ($CDCl_3$, 400MHz): δ 9.64 (d, *J* = 8.0 Hz, 1H), 8.53 (d, *J* = 8.0 Hz, 1H), 7.87 (d, *J* = 4.0 Hz, 1H), 7.61 (td, *J* = 8.0, 1.5 Hz, 1H), 7.48 (d, *J* = 8.0 Hz, 1H), 7.43 (t, *J* = 8.0 Hz, 1H), 7.32 (dd, *J* = 8.0, 4.0 Hz, 1H), 7.05 (d, *J* = 8.0 Hz, 1H), 6.19 (d, *J* = 24 Hz, 1H), 6.31 (d, *J* = 20 Hz, 1H), 2.07 (s, 3H), 1.38 (s, 9H); $^{13}$C NMR ($CDCl_3$, 150 MHz): δ 155.4, 154.4, 151.9, 140.2, 137.4, 136.8, 134.4, 132.3, 131.8, 128.9, 128.8, 127.1, 124.4, 120.4, 85.1, 59.7, 54.2, 28.0, 20.0, 18.9, 17.5; HRMS (ESI) m/z: [M]+ for $C_{20}H_{22}NO_2S$ calculated 340.1366 found 340.1364.

*Synthesis of 5-(carboxymethyl)-4-(o-tolyl)thieno[3,2-c]pyridin-5-ium(**10-p8**)*

The compound **10-p7** (300 mg, 0.88 mmol) was stirred in 4N HCl in dioxane (10 mL) for 1 h at rt. After completeness of reaction was observed by TLC ($R_f$ = 0.1, MeOH/DCM, 2:10), the solvent was removed under reduced pressure, and the residue was purified by silica gel column chromatography (DCM:MeOH, 49:1) to afford compound **10-p8** as a white solid (177 mg, 71% yield).

$^1$H NMR ($CD_3OD$, 400MHz): δ 8.75 (s, 2H), 8.20 (dd, *J* = 4.0, 1.7 Hz, 1H). 7.65 (t, *J* = 8.0 Hz, 1H), 7.67 (d, *J* = 8.0 Hz, 1H), 7.50 (t, *J* = 8.0 Hz, 1H), 7.13 (dd, *J* = 8.0, 2.0 Hz, 1H), 6.24 (d, *J* = 16 Hz, 1H), 4.94 (d, *J* = 16 Hz, 1H), 2.05 (s, 3H); $^{13}$C NMR ($CD_3OD$, 150 MHz): δ 155.5, 153.0, 139.0, 138.9, 138.0, 137.0, 133.1, 132.6, 130.6, 129.9, 128.1, 126.2, 121.7, 19.5; HRMS (ESI) m/z: [M]+ for $C_{16}H_{14}NO_2S$ calculated 284.0740 found 284.0738.

*Synthesis of 5-(2-((R)-3-Methylpiperazin-1-yl)-2-oxoethyl)-4-(o-tolyl)thieno[3,2-c]pyridin-5-ium (**10-p9**)*

To a stirred solution of compound **10-p8** (480 mg, 1.69 mmol, 1 eq) in DMF (10 mL) were added HATU (770 mg, 2.03 mmol, 1.2 eq), DIPEA (0.9 mL, 5.07mmol, 3 eq), and *tert*-butyl (*R*)-2-methylpiperazine-1-carboxylate (410 mg, 2.03 mmol, 1.2 eq). The reaction mixture was stirred for 3 h at rt. After completeness of reaction was observed by TLC ($R_f$ = 0.5, MeOH/DCM, 1:8), the reaction mixture was washed with brine (5 mL) and extracted with DCM (10 mL) 3 times. The organic layer was dried over $MgSO_4$, filtered and concentrated under reduced pressure. The residue was purified by flash silica gel column chromatography (MeOH:DCM, 3:97).

The purified product was stirred in 50% (v/v) TFA/DCM (3 mL) for 4 h at rt. After completeness of reaction was observed by TLC ($R_f$ = 0.3, MeOH/DCM, 1:8), the solvent was removed under reduced pressure, and the residue was purified by silica gel column chromatography (MeOH:DCM, 3:97) to afford compound **10-p9** as a white solid (320 mg, 52% yield).

$^1$H NMR ($CDCl_3$, 400MHz): δ 9.09-8.96 (m, 1H), 8.58-8.46 (m, 1H), 7.89 (dt, *J* = 8.0, 4.0 Hz, 1H), 7.68-7.61 (m, 1H), 7.54-7.42 (m, 2H), 7.31-7.27 (m, 1H), 7.11-7.05 (m, 1H), 6.39 (dd, *J* = 72, 16 Hz, 0.5H), 5.93-5.51 (m, 1H), 5.08 (dd, *J* = 36, 16 Hz, 0.5H), 4.32 (t, *J* = 12 Hz, 1H), 3.76 (dd, *J* = 28, 16 Hz, 1H), 3.60-3.42 (m, 1H), 3.35-2.87 (m, 5H), 2.03 (s, 3H), 1.37-1.22 (m, 3H); $^{13}$C NMR ($CDCl_3$, 150 MHz): δ 163.3, 163.2, 163.1, 162.3, 162.0, 161.9, 161.6, 154.3, 154.2, 152.8, 152.7, 152.6, 138.9, 138.8, 138.7, 137.6, 136.8, 136.3, 136.2, 134.6, 134.5, 132.8, 131.7, 128.9, 128.8, 128.7, 128.6, 128.5, 127.4, 127.3, 124.7, 124.6, 120.7, 120.6, 120.5, 118.0, 116.1, 58.6, 58.5, 51.0, 50.9, 50.8, 47.7, 46.3, 43.0, 42.8, 41.5, 41.4, 38.9, 29.8, 19.6, 19.5, 15.9, 15.8, 15.4, 15.3; HRMS (ESI) m/z: [M]+ for $C_{21}H_{24}N_3OS$ calculated 366.1635 found 366.1632.

*Synthesis of 5-(2-((S)-3-Methylpiperazin-1-yl)-2-oxoethyl)-4-(o-tolyl)thieno[3,2-c]pyridin-5-ium (**10-p10**)*

To a stirred solution of compound **10-p8** (480 mg, 1.69 mmol, 1 eq) in DMF (10 mL) were added HATU (700 mg, 2.03 mmol, 1.2 eq), DIPEA (0.9 mL, 5.07 mmol, 3 eq), and *tert*-butyl (*S)*-2-methylpiperazine-1-carboxylate (410 mg, 2.03 mmol, 1.2 eq). The reaction mixture was stirred for 3 h at rt. After completeness of reaction was observed by TLC ($R_f$ = 0.5, MeOH/DCM, 1:8), the reaction mixture was washed with brine (5 mL) and extracted with DCM (10 mL) 3 times. The organic layer was dried over $MgSO_4$, filtered, and concentrated under reduced pressure. The residue was purified by flash silica gel column chromatography (MeOH:DCM, 3:97).

The purified product was stirred in 50% (v/v) TFA/DCM (3 mL) for 4 h at rt. After completeness of reaction was observed by TLC ($R_f$ = 0.63, MeOH/DCM, 1:8), the solvent was removed under reduced pressure, and the residue was purified by silica gel column chromatography (MeOH:DCM, 3:97) to afford compound **10-p10** as a white solid (303 mg, 49% yield).

$^1$H NMR ($CDCl_3$, 400MHz): δ 9.09-8.98 (m, 1H), 8.58-8.47 (m, 1H), 7.89 (td, *J* = 4.0, 1.6 Hz, 1H), 7.68-7.59 (m, 1H), 7.54-7.40 (m, 2H), 7.31-7.27 (m, 1H), 7.12-7.04 (m, 1H), 6.36 (dd, *J* = 64, 20 Hz, 0.5H), 5.95-5.51 (m, 1H), 5.09 (dd, *J* = 36, 24 Hz, 0.5H), 4.31 (t, *J* = 12 Hz, 1H), 3.76 (dd, *J* = 20, 12 Hz, 1H). 3.60-3.40 (m, 1H), 3.59-2.68 (m, 6H), 2.03 (s. 3H). 1.39-1.16 (m, 3H); $^{13}$C NMR ($CDCl_3$, 150 MHz): δ 163.2, 163.1, 162.1, 161.9, 138.9, 138.8, 138.7, 137.6, 137.5, 134.5, 134.4, 132.8, 132.1, 131.8, 131.7, 128.9, 128.8, 127.5, 127.4, 127.3, 124.7, 124.6, 124.5, 120.5, 120.4, 118.0, 116.1, 68.4, 21.1, 50.8, 45.4, 43.0, 42.9, 42.8, 38.9, 19.6, 15.9, 15.8, 15.4, 15.3; HRMS (ESI) m/z: [M]+ for $C_{21}H_{24}N_3OS$ calculated 366.1635 found 366.1633.

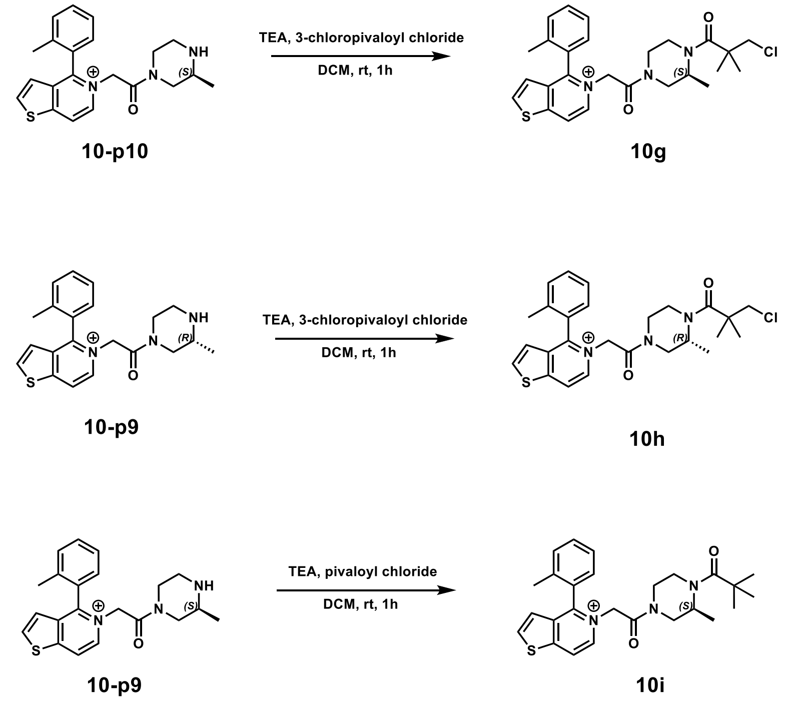


**Scheme 4.** Synthesis of oxidized electrophiles and binder **10g, 10h**, and **10i**.

*Synthesis of 5-(2-((S)-4-(3-Chloro-2,2-dimethylpropanoyl)-3-methylpiperazin-1-yl)-2-oxoethyl)-4-(o-tolyl)thieno[3,2-c]pyridin-5-ium (**10g**)*

To a stirred solution of **10-p10** (13.5 mg, 37 μmol, 1 eq) in DCM (0.75 mL) were added TEA (8.2 mg, 81 μmol, 2.2 eq) and 3-chloropivaloyl chloride (CAS: 4300-97-4, 6.3 mg, 41 μmol, 1.1 eq). The reaction mixture was stirred for 1 h at rt. After completeness of reaction was observed by

LC/MS, the reaction mixture was purified by RP-HPLC (60 min, 20-100% methanol in water + 0.1% (v/v) TFA), affording **10g** (4.2 mg, 23% yield).

$^1$H NMR (d6-DMSO, 400MHz): δ 9.01-8.95 (m, 1H), 8.90-8.83 (m, 1H), 8.40-8.33 (m, 1H), 7.68-7.25 (m, 4H), 7.15-7.06 (m, 1H), 5.76-5.00 (m, 2H), 4.57-4.34 (m, 1H), 4.17-3.91 (m, 2H), 3.81-3.68 (m, 3H), 3.16-2.64 (m, 3H), 2.04-1.96 (m, 3H), 1.29-1.22 (m, 6H), 1.01-0.50 (m, 3H); HRMS (ESI) m/z: [M]+ for $C_{26}H_{31}ClN_3O_2S^+$ calculated 484.1820 found 484.1817.

*Synthesis of 5-(2-((R)-4-(3-Chloro-2,2-dimethylpropanoyl)-3-methylpiperazin-1-yl)-2-oxoethyl)-4-(o-tolyl)thieno[3,2-c]pyridin-5-ium (**10h**)*

To a stirred solution of **10-p9** (3.0 mg, 8.2 μmol, 1 eq) in DCM (0.5 mL) were added TEA (2.0 mg, 20 μmol, 2.4 eq) and 3-chloropivaloyl chloride (CAS: 4300-97-4, 1.5 mg, 10 μmol, 1.2 eq). The reaction mixture was stirred for 1 h at rt. After completeness of reaction was observed by LC/MS, the reaction mixture was purified by RP-HPLC (60 min, 20-100% methanol in water + 0.1% (v/v) TFA), affording **10h** (3.4 mg, 86% yield).

$^1$H NMR (d6-DMSO, 400MHz): δ 8.99-8.95 (m, 1H), 8.90-8.83 (m, 1H), 8.37-8.32 (m, 1H), 7.68-7.25 (m, 4H), 7.15-7.06 (m, 1H), 5.75-5.02 (m, 2H), 4.56-4.32 (m, 1H), 4.17-3.89 (m, 2H), 3.82-3.66 (m, 2H), 3.65-3.56 (m, 1H), 2,97-2.64 (m, 3H), 2.04-1.94 (m, 3H), 1.28-1.24 (m, 6H), 0.98-0.51 (m, 3H); HRMS (ESI) m/z: [M]+ for $C_{26}H_{31}ClN_3O_2S^+$ calculated 484.1820 found 484.1818.

*Synthesis of 5-(2-((S)-3-Methyl-4-pivaloylpiperazin-1-yl)-2-oxoethyl)-4-(o-tolyl)thieno[3,2-c]pyridin-5-ium (**10i**)*

To a stirred solution of **10-p10** (13.5 mg, 37 μmol, 1 eq) in DCM (0.75 mL) were added TEA (8.2 mg, 81 μmol, 2.2 eq) and pivaloyl chloride (CAS: 3282-30-2, 4.9 mg, 41 μmol, 1.1 eq). The reaction mixture was stirred for 1 h at rt. After completeness of reaction was observed by LC/MS, the reaction mixture was purified by RP-HPLC (60 min, 20-100% methanol in water + 0.1% (v/v) TFA), affording **10i** (5.0 mg, 30 % yield).

$^1$H NMR (d6-DMSO, 400MHz): δ 8.99-8.95 (m, 1H), 8.88-8.83 (m, 1H), 8.37-8.34 (m, 1H), 7.68-7.25 (m, 4H), 7.15-7.06 (m, 1H), 5.82-5.02 (m, 2H), 4.56-4.34 (m, 1H), 4.17-3.89 (m, 2H), 3.64-3.33 (m, 1H, overlap with $H_2O$), 3.22-2.61 (m, 3H), 2.04-1.96 (m, 3H), 1.21-1.10 (m, 9H), 0.96-0.50 (m, 3H); HRMS (ESI) m/z: [M]+ for $C_{26}H_{32}N_3O_2S^+$ calculated 450.2210 found 450.2207.

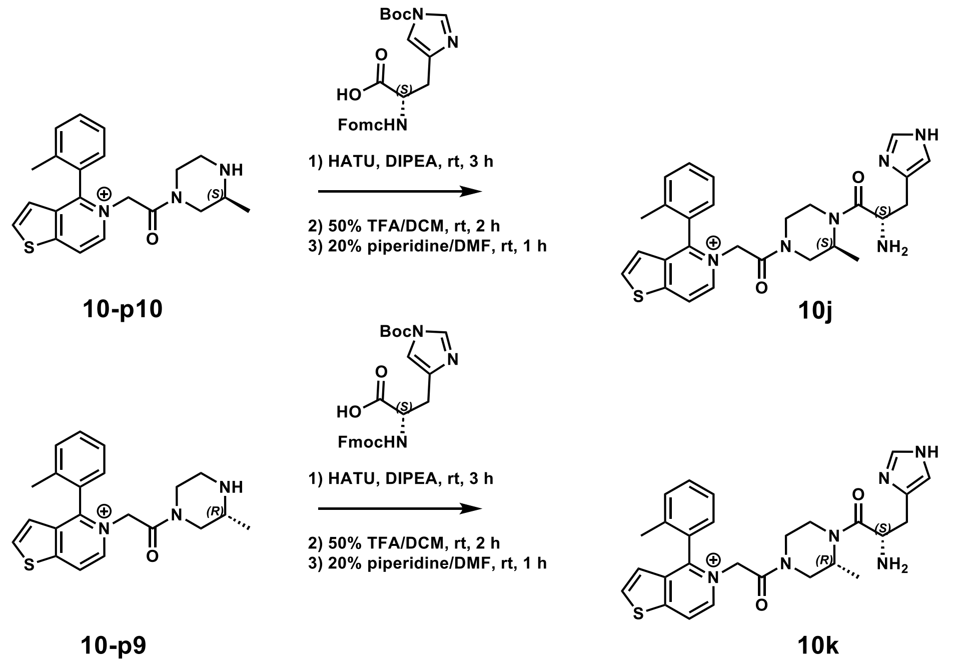


**Scheme 5.** Synthesis of histidine-derived base triple compounds **10j** and **10k**.

*Synthesis of 5-(2-((S)-4-(L-Histidyl)-3-methylpiperazin-1-yl)-2-oxoethyl)-4-(o-tolyl)thieno[3,2-c]pyridin-5-ium (**10j**)*

To a stirred solution of Fmoc-His(Boc)-OH (CAS: 81379-52-4, 100 mg, 0.21 mmol, 2 eq) in DMF (5 mL) were added HATU (80 mg, 0.21 mmol, 2 eq) and DIPEA (80 µL, 0.44 mmol, 4 eq). The reaction mixture was stirred for 20 min at rt, followed by addition of **10-p10** (40 mg, 0.11 mmol, 1 eq). The reaction mixture was stirred for 3 h at rt. After completeness of reaction was observed by TLC ($R_f$ = 0.8, MeOH/DCM, 1:7), the solvent was removed under reduced pressure, and the residue was purified by flash silica gel column chromatography (0-20% MeOH in DCM).

The purified mixture was stirred in 50% (v/v) TFA/DCM (1 mL) for 2 h at rt. After completeness of reaction was observed by TLC ($R_f$ = 0.3, MeOH/DCM, 1:8), the solvent was removed under reduced pressure. Next, 20% (v/v) piperidine/DMF (1 mL) was added to the reaction mixture and stirred for 1 h at rt. After, the solvent was removed under reduced pressure, and the residue was purified by RP-HPLC to afford **10j** as a white solid (4.7 mg, 8% yield).

$^1$H NMR ($CD_3OD$, 400MHz): δ 8.94-8.88 (m, 1H), 8.79 (dd, *J* = 4, 2 Hz, 1H), 8.76-8.69 (m, 1H), 8.24 (d, *J* = 4 Hz, 1H), 7.69-7.31 (m, 5H), 7.16 (q, *J* = 4 Hz, 1H), 5.89-5.58 (m, 1H), 5.54-5.30 (m, 1H), 4.72-4.11 (m, 3.5 H), 3.87-3.33 (m, 2.5 H), 3.28-3.14 (m, 2H), 3.13-2.62 (m, 2H), 2.11-2.03 (m, 3H), 1.07 (td, *J* = 24, 8 Hz, 1.5H), 0.09-0.62 (m, 1.5H); $^{13}$C NMR ($CD_3OD$, 150 MHz): δ 168.0, 167.9, 167.3, 165.5, 165.4, 165.3, 165.1, 165.0, 162.9, 162.7, 156.5, 156.4, 156.3, 156.2, 1553.4, 153.3, 139.3, 139.2, 139.1, 139.0, 138.9, 138.7, 138.6, 138.1, 137.9, 137.5, 137.4, 136.2, 133.4, 133.3, 133.1, 132.9, 132.8, 130.4, 130.3, 130.2, 130.1, 130.0, 129.9, 128.2, 128.1, 128.0, 127.9, 125.3, 125.2, 125.1, 121.9, 121.8, 121.7, 120.1, 120.0, 119.0, 117.0, 60.0, 59.7, 59.6, 59.5, 50.6, 50.5, 50.4, 50.3, 50.1, 50.0, 47.8, 47.6, 47.5, 47.4, 47.3, 46.9, 45.6, 45.4, 43.3, 41.6, 41.5, 41.3,

41.2, 37.9, 37.6, 27.4, 27.2, 19.7, 19.6, 19.5, 19.4, 16.5, 16.3, 15.8, 15.6, 15.2, 15.0, 14.3, 14.1; HRMS (ESI) m/z: [M+]+ for $C_{27}H_{32}N_6O_2S$ calculated 503.2224 found 503.2227.

*Synthesis of 5-(2-((R)-4-(L-Histidyl)-3-methylpiperazin-1-yl)-2-oxoethyl)-4-(o-tolyl)thieno[3,2-c]pyridin-5-ium (**10k**)*

To a stirred solution of Fmoc-His(Boc)-OH (100 mg, 0.21 mmol, 2 eq) in DMF (5 mL) were added HATU (80 mg, 0.21 mmol, 2 eq) and DIPEA (80 µL, 0.44 mmol, 4 eq). The reaction mixture was stirred for 20 min at rt, and then **10-p9** (40 mg, 0.11 mol, 1 eq) was added. The reaction mixture was stirred for 3 h at rt. After completeness of reaction was observed by TLC ($R_f$ = 0.8, MeOH/DCM, 1:7), the solvent was removed under reduced pressure and the residue was purified by flash silica gel column chromatography (0-20% MeOH in DCM).

The purified mixture was stirred in 50% (v/v) TFA/DCM (1 mL) for 2 h at rt. A fter completeness of reaction was observed by TLC ($R_f$ = 0.3, MeOH/DCM, 1:8), the solvent was removed under reduced pressure. After the solvent was removed, 20% (v/v) piperidine/DMF (1 mL) was added to the reaction mixture, which was stirred for 1 h at rt. After, the solvent was removed under reduced pressure and residue was purified by RP-HPLC to afford compound **10k** as a white solid (6.0 mg, 11% yield).

$^1$H NMR ($CD_3OD$, 400MHz): δ 8.84-8.66 (m, 2H), 8.23 (t, *J* = 4 Hz, 1H), 7.86-7.29 (m, 5H), 7.21-7.13 (m, 1H), 7.11-6.98 (m, 1H), 5.92-5.19 (m, 2H), 4.65-4.42 (m, 2H), 4.36-3.98 (m, 2H), 3.97-3.55 (m. 2H), 3.51-3.38 (m, 1H), 3.09-2.76 (m, 3H), 2.06 (s, 3H), 1.15-0.50 (m, 3H); $^{13}$C NMR ($CD_3OD$, 150 MHz): δ 169.3, 169.0, 165.7, 165.4, 165.3, 165.2, 165.1, 166.4, 163.1, 163.0, 165.4, 156.3, 156.2, 156.1, 153.4, 153.3, 153.2, 139.6, 139.4, 139.3, 139.2, 139.1, 139.0, 138.9, 138.8, 138.7, 138.6, 138.2, 138.1, 138.0, 137.9, 137.5, 137.4, 137.3, 137.2, 133.5, 133.4, 133.3, 133.2, 133.1, 133.0, 132.9, 132.8, 132.7, 130.4, 130.3, 130.2, 130.1, 130.0, 129.9, 128.2, 128.1, 128.0, 125.3, 125.2, 125.1, 122.0, 121.9, 121.8, 121.7, 119.3, 118.4, 118.0, 117.3, 59.2, 51.6, 47.4, 47.2, 47.1, 46.6, 46.5, 46.4, 46.2, 45.4, 45.3, 45.2, 43.5, 43.4, 41.0, 40.8, 40.7, 37.9, 37.8, 37.5, 37.4, 31.0, 30.0, 19.7, 19.6, 19.5, 19.4, 16.55, 16.4, 15.7, 15.5, 15.4, 15.3, 14.5, 14.3; HRMS (ESI) m/z: [M]+ for $C_{27}H_{32}N_6O_2S$ calculated 503.2224 found 503.2219.

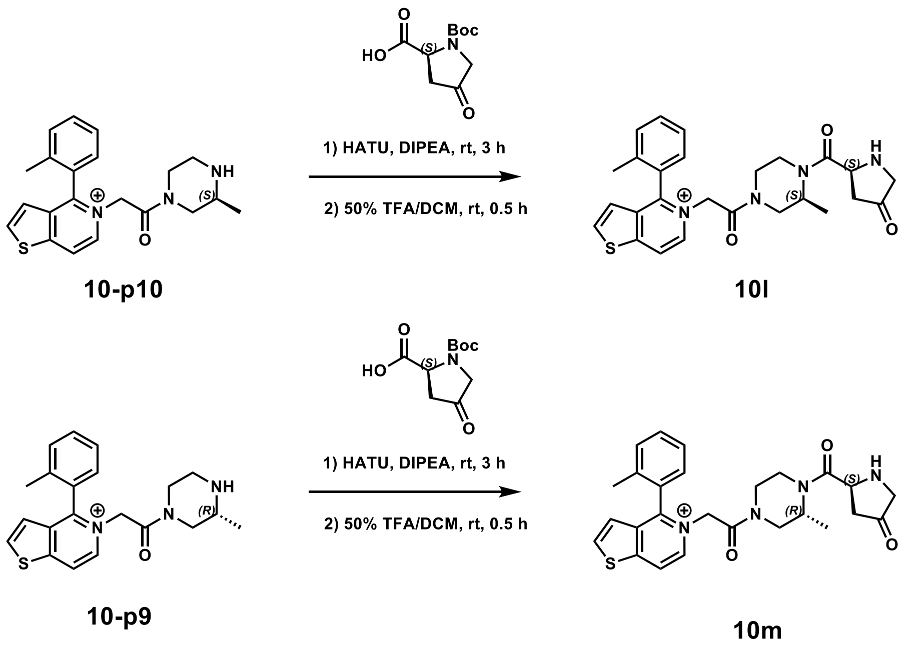


**Scheme 6.** Synthesis of 4-oxoproline-derived base triple compounds **10l** and **10m**.

*Synthesis of 5-(2-((S)-3-Methyl-4-((S)-4-oxopyrrolidine-2-carbonyl)piperazin-1-yl)-2-oxoethyl)-4-(o-tolyl)thieno[3,2-c]pyridin-5-ium (**10l**)*

To a stirred solution of (*S*)-1-(*tert*-butoxycarbonyl)-4-oxopyrrolidine-2-carboxylic acid (CAS: 84348-37-8, 36 mg, 160 μmol, 2 eq) in DMF (5 mL) were added HATU (60 mg, 160 μmol, 2 eq) and DIPEA (60 μL, 33 μmol, 4 eq). The reaction mixture was stirred for 20 min at rt, then **10-p10** (30 mg, 80 μmol, 1 eq) was added. The reaction mixture was stirred for 3 h at rt. After completeness of reaction was observed by TLC ($R_f$ = 0.7, MeOH/DCM, 1:5), the solvent was removed under reduced pressure and the residue was purified by flash silica gel column chromatography (0-20% MeOH in DCM).

The purified mixture was stirred in 50% (v/v) TFA/DCM (2 mL) for 30 min at rt. After completeness of reaction was observed by TLC ($R_f$ = 0.1, MeOH/DCM, 1:5), the solvent was removed under reduced pressure, and the residue was purified by RP-HPLC to afford compound **10l** as a white solid (6.1 mg, 32% yield).

$^1$H NMR ($CD_3OD$, 400MHz): δ 8.81-8.68 (m, 2H), 8.26-8.22 (m, 1H), 7.71-7.33 (m. 4H), 7.20-7.13 (m, 1H), 5.87-5.59 (m, 1H), 5.48-5.17 (m, 1H), 4.74-4.50 (m, 1H), 4.46-3.93 (m, 2H), 3.85-3.36 (m, 3H), 3.26-2.48 (m, 4H), 2.07 (s, 3H), 1.21-0.65 (m, 3H); $^{13}$C NMR ($CD_3OD$, 150 MHz): δ 168.1, 168.0, 167.8, 167.7, 165.5, 165.3, 156.5, 156.3, 156.2, 153.4, 153.3, 139.3, 139.1, 137.5, 133.4, 133.3, 133.1, 132.9, 132.8, 130.4, 130.3, 130.1, 130.0, 128.2, 128.1, 125.3, 125.2, 125.1, 121.9, 121.8, 121.7, 60.0, 59.7, 59.6, 58.6, 58.4, 55.8, 47.8, 47.7, 47.5, 45.5, 45.2, 43.3, 38.1, 37.7, 19.7, 19.6, 19.5, 18.7, 17.3, 15.3, 14.4, 14.2; HRMS (ESI) m/z: [M]+ for $C_{26}H_{29}N_4O_3S$ calculated 477.1955 found 477.1951.

*Synthesis of 5-(2-((R)-3-Methyl-4-((S)-4-oxopyrrolidine-2-carbonyl)piperazin-1-yl)-2-oxoethyl)-4-(o-tolyl)thieno[3,2-c]pyridin-5-ium (**10m**)*

To a stirred solution of (*S*)-1-(*tert*-butoxycarbonyl)-4-oxopyrrolidine-2-carboxylic acid (CAS: 84348-37-8, 36 mg, 160 µmol, 2 eq) in DMF (5 mL) were added HATU (60 mg, 160 µmol, 2 eq) and DIPEA (60 µL, 330 µmol, 4 eq). The reaction mixture was stirred for 20 min at rt, and then **10-p9** (30 mg, 80 µmol, 1 eq) was added. The reaction mixture was stirred for 3 h at rt. After completeness of reaction was observed by TLC ($R_f$ = 0.7, MeOH/DCM, 1:5), the solvent was removed under reduced pressure and the residue was purified by flash silica gel column chromatography (0-20% MeOH in DCM).

The purified mixture was stirred in 50% (v/v) TFA/DCM (2 mL) for 30 min at rt. After completeness of reaction was observed by TLC ($R_f$ = 0.2, MeOH/DCM, 1:5), the solvent was removed under reduced pressure, and the residue was purified by RP-HPLC to afford compound **10m** as a white solid (5.7 mg, 30% yield).

$^{1}$H NMR ($CD_3OD$, 400MHz): δ 8.81-8.68 (m, 2H), 8.24 (d. *J* = 4.0 Hz, 1H), 7.69-7.34 (m, 4H), 7.19-7.13 (m, 1H), 5.89-5.59 (m, 1H), 5.50-5.16 (m, 1H). 4.82-4.58 (m, 1H), 4.33-4.11 (m, 1H), 4.06-3.38 (m, 3H), 3.27-2.51 (m, 4H), 2.047 (s, 3H), 2.01-1.84 (m, 1H), 1.21-0.65 (m, 3H); $^{13}$C NMR (CD3OD, 150 MHz): δ 168.1, 168.0, 167.8, 167.7, 165.5, 165.4, 165.3, 165.2, 165.1, 165.0, 162.7, 162.4, 156.5, 156.4, 156.3, 156.2, 153.4, 153.3, 139.3, 139.2, 139.1, 139.0, 138.8, 138.7, 138.6, 138.5, 138.4, 138.2, 138.0, 137.5, 137.4, 133.4, 133.3, 133.2, 133.1, 133.0, 132.9, 132.8, 130.4, 130.3, 130.2, 130.1, 129.9, 128.3, 128.2, 128.1, 125.3, 125.2, 125.1, 121.9, 121.8, 121.7, 119.0, 117.0, 104.4, 60.0, 59.9, 59.8, 59.7, 59.6, 59.5, 58.4, 58.2, 53.9, 53.7, 53.5, 47.4, 47.1, 47.0, 46.8, 45.3, 45.2, 43.3, 43.2, 42.9, 41.2, 40.9, 40.8, 38.2, 38.1, 37.9, 37.8, 19.7, 19.6, 19.5, 16.2, 16.0, 15.4, 15.3, 15.2, 15.1, 15.0, 14.5, 14.3, 14.2; HRMS (ESI) m/z: [M]+ for $C_{26}H_{29}N_4O3S$ calculated 477.1955 found 477.1951.

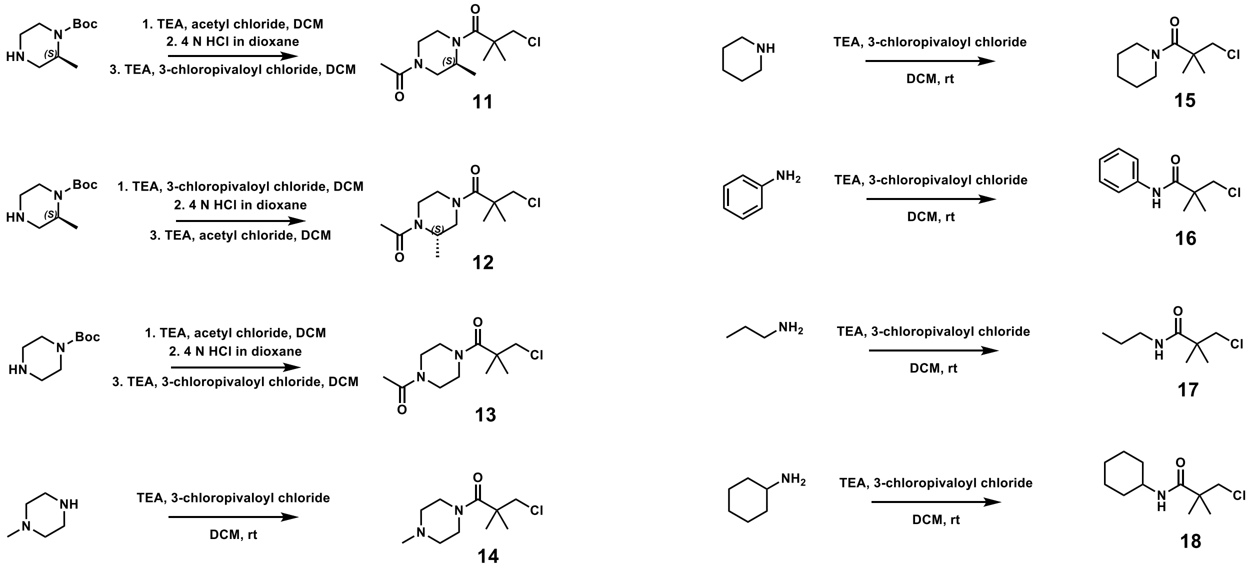


**Scheme 7.** Synthesis of electrophile library (**11-18**) for structure-reactivity relationships.

*Synthesis of (S)-1-(4-Acetyl-2-methylpiperazin-1-yl)-3-chloro-2,2-dimethylpropan-1-one (**11**)*

To a stirred solution of *tert*-butyl (*S*)-2-methylpiperazine-1-carboxylate (CAS: 169447-70-5, 22.8 mg, 114 µmol, 1 eq) and TEA (38.1 µL, 273 µmol, 2.4 eq) in DCM (0.5 mL) was added acetyl chloride (CAS: 75-36-5, 9.7 µL, 137 µmol, 1.2 eq). The reaction was stirred for 1 h at rt, and reaction completion was confirmed by LC/MS. The reaction was quenched by addition of water (0.5 mL), and the crude product was extracted 3 times with 0.5 mL of DCM. The solvent was dried with sodium sulfate, filtered, and the solvent removed under reduced pressure. The crude product was resuspended with 0.5 mL of DCM followed by addition of 0.5 mL 4 N HCl in dioxane. The reaction was stirred at rt for 1 h, then the solvent was removed under reduced pressure. The crude product was resuspended with 0.5 mL of DCM, to which TEA (84 µL, 600 µmol, ~5 eq) and 3-chloropivaloyl chloride (CAS: 4300-97-4, 17.7 µL, 137 µmol, ~1.2 eq) were added. The reaction was stirred for 1 h at rt, and reaction progression was monitored by LC/MS. Solvent was removed under reduced pressure, and the product was purified by RP-HPLC (60 min, 20-100% methanol:water + 0.1% (v/v) TFA), yielding **11** (12.4 mg, 48 µmol, 42% yield).

$^1$H NMR (DMSO-d6, 400MHz): δ 4.52 (m, 1H), 4.30-3.99 (m, 2H), 3.86-3.56 (m, 3H), 3.29-2.97 (m, 2H), 2.82-2.58 (m, 1H), 2.05 (s, 1.5H), 1.99 (s, 1.5H), 1.32-1.26 (m, 6H), 1.11 (d, *J* = 6.7 Hz, 1.5H), 1.04 (d, *J* = 6.6 Hz, 1.5H); HRMS (ESI) m/z: [M + H]+ for $C_{12}H_{22}ClN_2O_2^+$ calculated 261.1364 found 261.1364.

*Synthesis of (S)-1-(4-Acetyl-3-methylpiperazin-1-yl)-3-chloro-2,2-dimethylpropan-1-one (**12**)*

To a stirred solution of *tert*-butyl (*S*)-2-methylpiperazine-1-carboxylate (CAS: 169447-70-5, 24.0 mg, 120 µmol, 1 eq) and TEA (40.1 µL, 288 µmol, 2.4 eq) in DCM (1 mL) was added 3-chloropivaloyl chloride (CAS: 4300-97-4, 18.6 µL, 144 µmol, 1.2 eq). The reaction was stirred for 1 h at rt, and reaction completion was confirmed by LC/MS. The reaction was quenched by addition of water (0.5 mL), and the crude product was extracted 3 times with 0.5 mL of DCM. The solvent was dried with sodium sulfate, filtered, and the solvent removed under reduced pressure. The crude product was resuspended with 0.5 mL of DCM followed by addition of 0.5 mL 4 N HCl in dioxane. The reaction was stirred at rt for 1 h, then the solvent was removed under reduced pressure. The crude product was resuspended with 0.5 mL of DCM, to which TEA (80 µL, 574 µmol, ~5 eq) and acetyl chloride (10.2 µL, 144 µmol, ~1.2 eq) were added. The reaction was stirred for 1 h at rt and reaction progression was monitored by LC/MS. Solvent was removed under reduced pressure and the product was purified by RP-HPLC (60 min, 20-100% methanol:water + 0.1% (v/v) TFA), yielding **12** (15.3 mg, 59 µmol, 49% yield).

$^1$H NMR (DMSO-d6, 400MHz): δ 4.57-4.46 (m, 0.5H), 4.23-4.03 (m, 3H), 3.88-3.80 (m, 1H), 3.80-3.74 (m, 1H), 3.70-3.60 (m, 0.5H), 3.35-3.20 (m, 0.5H), 3.11-2.85 (m, 2H), 2.85-2.72 (m, 0.5H), 2.03 (s, 1.5H), 2.00 (s, 1.5H), 1.30 (s, 3H), 1.29 (s, 3H), 1.12 (d, *J* = 6.3 Hz, 1.5H), 1.01 (d, *J* = 6.5 Hz, 1.5H); HRMS (ESI) m/z: [M + H]+ for $C_{12}H_{22}ClN_2O_2^+$ calculated 261.1364 found 261.1363.

*Synthesis of 1-(4-Acetylpiperazin-1-yl)-3-chloro-2,2-dimethylpropan-1-one (**13**)*

To a stirred solution of *tert*-butyl piperazine-1-carboxylate (CAS: 57260-71-6, 22.9 mg, 123 µmol, 1 eq) and TEA (41.1 µL, 295 µmol, 2.4 eq) in DCM (0.5 mL) was added acetyl chloride (CAS: 75-36-5, 10.5 µL, 148 µmol, 1.2 eq). The reaction was stirred for 1 h at rt, and reaction completion was confirmed by LC/MS. The reaction was quenched by addition of water (0.5 mL), and the crude product was extracted 3 times with 0.5 mL of DCM. The solvent was dried with sodium sulfate, filtered, and the solvent removed under reduced pressure. The crude product was resuspended with 0.5 mL of DCM and then 0.5 mL 4 N HCl in dioxanes was added. The reaction was stirred at rt for 1 h, followed by removing solvent under reduced pressure. The crude product was resuspended in 0.5 mL of DCM, to which TEA (84 µL, 600 µmol, ~5 eq) and 3-chloropivaloyl chloride (CAS: 4300-97-4, 17.7 µL, 137 µmol, 1.1 eq) were added. The reaction was stirred for 1 h at rt, and reaction progress was monitored by LC/MS. Solvent was removed under reduced pressure, and the product was purified by RP-HPLC (60 min, 20-100% methanol:water + 0.1% (v/v) TFA), yielding **13** (18.1 mg, 73 µmol, 60% yield).

$^1$H NMR (DMSO-d6, 400MHz): δ 3.78 (s, 2H), 3.60-3.49 (m, 4H), 3.46-3.40 (m, 4H), 2.02 (s, 3H), 1.29 (s, 6H); HRMS (ESI) m/z: [M + H]+ for $C_{11}H_{20}ClN_2O_2^+$ calculated 247.1208 found 247.1207.

*Synthesis of 3-Chloro-2,2-dimethyl-1-(4-methylpiperazin-1-yl)propan-1-one (**14**)*

A stirred solution of 1-methylpiperazine (CAS: 109-01-3, 20 mg, 200 µmol, 1 eq) and TEA (49 mg, 480 µmol, 2.4 eq) in dry DCM (1 mL) was brought to 0 °C in an ice bath. To this solution, 3-chloropivaloyl chloride (CAS: 4300-97-4, 37 mg, 240 µmol, 1.2 eq) was slowly added. After addition, the solution was brought to room temperature and stirred for 2 h. The solvent was removed under reduced pressure, and and the product was purified by silica gel column chromatography (0-100% Hx:DCM), affording **14** (29.4 mg, 67% yield).

$^1$H NMR ($CDCl_3$, 400MHz): δ 3.72 (s, 2H), 3.68 (t, *J* = 4.3 Hz, 4H), 2.46 (t, *J* = 4.7 Hz, 4H), 2.34 (s, 3H), 1.39 (s, 6H); HRMS (ESI) m/z: [M + H]+ for $C_{10}H_{20}ClN_2O^+$ calculated 219.1259 found 219.1259.

*Synthesis of 3-Chloro-2,2-dimethyl-1-(piperidin-1-yl)propan-1-one (**15**)*

A stirred solution of piperidine (CAS: 110-89-4, 18 mg, 210 µmol, 1 eq) and TEA (51 mg, 510 µmol, 2.4 eq) in dry DCM (1 mL) was brought to 0 °C in an ice bath. To this solution, 3-chloropivaloyl chloride (CAS: 4300-97-4, 39 mg, 250 µmol, 1.2 eq) was slowly added. After addition, the solution was brought to room temperature and stirred for 2 h. The solvent was removed under reduced pressure, and the product was purified by silica gel column chromatography (0-100% Hx:DCM), affording **15** (17.2 mg, 40% yield).

$^1$H NMR ($CDCl_3$, 400MHz): δ 3.73 (s, 2H), 3.56 (t, *J* = 5.4 Hz, 4H), 1.70-1.62 (m, 2H), 1.62-1.53 (m, 4H), 1.38 (s, 6H); HRMS (ESI) m/z: [M + H]+ for $C_{10}H_{19}ClNO^+$ calculated 204.1150 found 204.1149.

*Synthesis of 3-Chloro-2,2-dimethyl-N-phenylpropanamide (**16**)*

A stirred solution of aniline (CAS: 62-53-3, 50 mg, 540 µmol,1 eq) and TEA (130 mg, 1.3 mmol, 2.4 eq) in dry DCM (2 mL) was brought to 0 °C in an ice bath. To this solution, 3-chloropivaloyl chloride (CAS: 4300-97-4, 100 mg, 640 µmol, 1.2 eq) was slowly added. After addition, the solution was stirred for 5 h at rt. The solvent was removed under reduced pressure, and the product was purified using silica gel column chromatography (1:9 EtOAc:Hx) to afford a white solid, **16** (94.2 mg, 83% yield).

$^1$H NMR ($CDCl_3$, 400MHz): δ 7.57 (bs, 1H), 7.52 (d, *J* = 7.6 Hz, 2H), 7.31 (t, *J* = 7.7 Hz, 2H), 7.12 (t, *J* = 7.4 Hz, 1H), 3.69 (s, 2H), 1.40 (s, 6H); HRMS (ESI) m/z: [M + H]+ for $C_{11}H_{15}ClNO^+$ calculated 212.0837 found 212.0835.

*Synthesis of 3-Chloro-2,2-dimethyl-N-propylpropanamide (**17**)*

A stirred solution of propylamine (CAS: 107-10-8, 20 mg, 340 µmol, 1 eq) and TEA (82 mg, 810 µmol, 2.4 eq) in dry DCM (2 mL) was brought to 0 °C in an ice bath. To this solution, 3-chloropivaloyl chloride (CAS: 4300-97-4, 63 mg, 410 µmol) was slowly added. After addition, the solution was brought to room temperature and stirred for 2 h. The reaction was purified by silica gel column chromatography (0-100% DCM in hexanes). The solvent was removed under reduced pressure, yielding a clear liquid, **17** (27.8 mg, 46% yield).

$^1$H NMR ($CDCl_3$, 400MHz): δ 5.81 (bs, 1H), 3.62 (s, 2H), 3.24 (q, *J* = 6.6 Hz, 2H), 1.54 (sext, *J* = 7.3 Hz, 2H), 1.29 (s, 6H), 0.93 (t, *J* = 7.4 Hz, 3H); HRMS (ESI) m/z: [M + H]+ for $C_8H_{17}ClNO^+$ calculated 178.0993 found 178.0993.

*Synthesis of 3-Chloro-N-cyclohexyl-2,2-dimethylpropanamide (**18**)*

A stirred solution of cyclohexamine (CAS: 108-91-8, 20 mg, 200 µmol, 1 eq) and TEA (49 mg, 480 µmol, 2.4 eq) in dry DCM (1 mL) was brought to 0 °C in an ice bath. To this solution, 3-chloropivaloyl chloride (CAS: 4300-97-4, 38 mg, 240 µmol) was slowly added. After addition, the solution was brought to room temperature and stirred for 2 h. Solvent was removed under reduced pressure, and the product was purified by RP-HPLC, affording **18** (7.8 mg, 18% yield).

$^1$H NMR (DMSO-d6, 400MHz): δ 7.25 (d, *J* = 7.9 Hz, 1H), 3.69 (s, 2H), 3.60-3.49 (m, 1H), 1.74-1.62 (m, 4H), 1.62-1.52 (m, 1H), 1.31-1.17 (m, 4H), 1.14 (s, 6H), 1.12-1.01 (m, 1H); HRMS (ESI) m/z: [M + H]+ for $C_{11}H_{21}ClNO^+$ calculated 218.1306 found 218.1305.

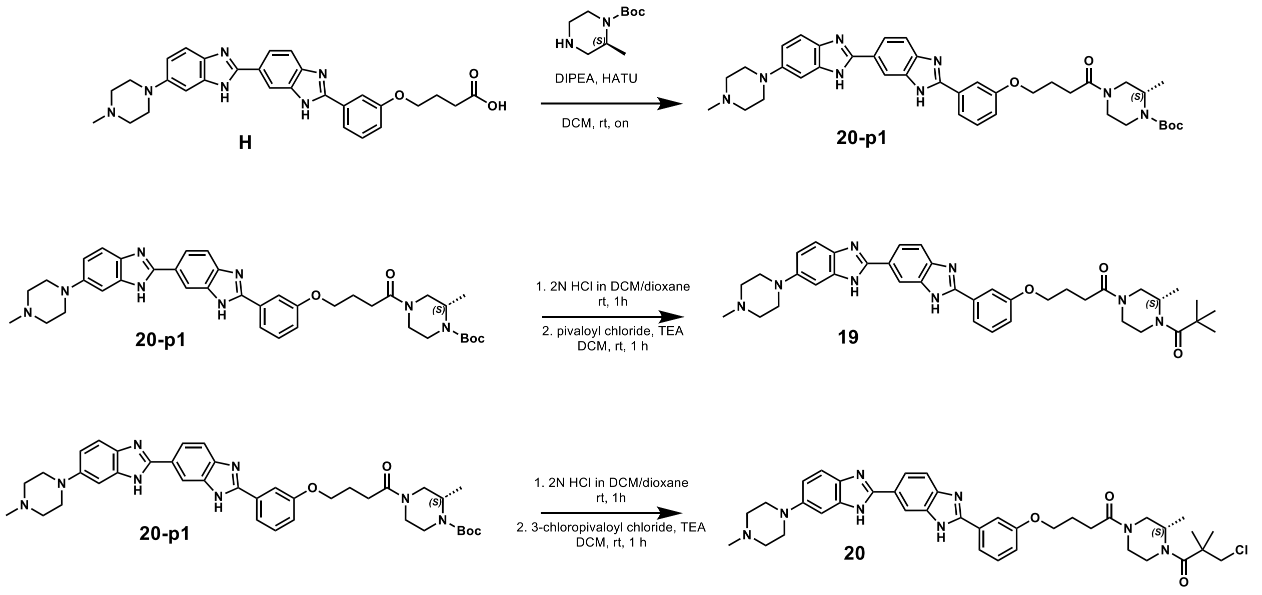

**Scheme 8.** Synthesis of non-covalent (**19**) and covalent (**20**) derivatives of **H**.

*Synthesis of 4-(3-(6-(4-Methylpiperazin-1-yl)-1H,3'H-[2,5'-bibenzo[d]imidazol]-2'-yl)phenoxy)butanoic acid (**H**).* **H** was synthesized as previously described.[20]

*Synthesis of tert-butyl-(S)-2-Methyl-4-(4-(3-(6-(4-methylpiperazin-1-yl)-1H,3'H-[2,5'-bibenzo[d]imidazol]-2'-yl)phenoxy)butanoyl)piperazine-1-carboxylate (**20-p1**)*

HATU (164 mg, 430 µmol, 1.1 eq) was added to a stirred solution of **H** (200 mg, 390 µmol, 1 eq) and DIPEA (152 mg, 1.18 mmol, 3.0 eq) in DCM (2 mL). After 10 min at rt, (*S*)-*tert*-butyl 2-methylpiperazine-1-carboxylate (CAS: 169447-70-5, 157 mg, 780 µmol, 2 eq) was added. The reaction was stirred at rt overnight. The reaction was then washed with water (2 mL, twice), and the organic layer was purified by RP-HPLC (40-100% MeOH in water + 0.1% (v/v) TFA). Fractions were lyophilized to afford a yellow solid, **20-p1** (136 mg, 50% yield).

$^1$H NMR (DMSO-d6, 400MHz): δ 10.12 (bs, 1H), 8.49 (s, 1H), 8.07 (dd, *J* = 8.6, 1.6 Hz, 1H), 7.92 (d, *J* = 8.6 Hz, 1H), 7.84 (d, *J* = 8.3 Hz, 1H), 7.82 (s, 1H), 7.73 (d, *J* = 9.0 Hz, 1H), 7.52 (t, *J* = 7.9 Hz, 1H), 7.33 (dd, $J_1$ = 9.1, 2.0 Hz, 1H), 7.24 (d, *J* = 1.9 Hz, 1H), 7.18-7.13 (m, 1 H), 4.30-3.79 (m, 7H), 3.78-3.46 (M, 4H), 3.35-3.01 (m, 5H), 3.01-2.78 (m, 4H), 2.76-2.52 (m, 2H), 2.11-1.97 (m, 2H), 1.40 (d, *J* = 1.0 Hz, 9H), 1.07 (d, *J* = 6.7 Hz, 1.5H), 0.99 (d, *J* = 6.6 Hz, 1.5H); HRMS (ESI) m/z: [M + H]+ for $C_{39}H_{49}N_8O_8^+$ calculated 693.3871 found 693.3867.

*Synthesis of* (S)-1-(3-Methyl-4-pivaloylpiperazin-1-yl)-4-(3-(6-(4-methylpiperazin-1-yl)-1H,3'H-[2,5'-bibenzo[d]imidazol]-2'-yl)phenoxy)butan-1-one (**19**)

**20-p1** (21 mg, 30 µmol, 1 eq) was stirred for 1 h at rt with 2 N HCl in 50:50 dioxane:DCM (2 mL). Once the reaction was confirmed complete by LC/MS, solvent was removed under reduced pressure. The crude product was dissolved in DCM (0.6 mL) and stirred at 0 °C in an ice bath.

To this solution, triethylamine (17 mg, 170 μmol, ~6 eq) and pivaloyl chloride (CAS: 3282-30-2, 4.1 mg, 34 μmol, 1.1 eq) were added, and then the solution was brought to rt and stirred for 1 h. The product was purified by RP-HPLC (60 min, 40-100% methanol:water + 0.1% (v/v) TFA). Fractions were lyophilized to afford a yellow solid, **19** (17.5 mg, 86% yield).

$^1$H NMR (DMSO-d6, 400MHz): δ 10.20 (b, 1H), 8.50 (d, *J* = 0.9 Hz, 1H), 8.07 (dd, *J* = 8.5, 1.6 Hz, 1H), 7.92 (d, *J* = 8.5 Hz, 1H), 7.84 (d, *J* = 7.7 Hz, 1H), 7.82 (s, 1H), 7.73 (d, *J* = 9.0 Hz, 1H), 7.52 (t, *J* = 7.9 Hz, 1H), 7.34 (dd, *J* = 9.1, 2.1 Hz, 1H), 7.26 (d, *J* = 2.0 Hz, 1H), 7.20-7.13 (m, 1 H), 4.53 (m, 1H), 4.35-4.01 (m, 4H), 4.01-3.68 (m, 3H), 3.68-3.46 (m, 2H), 3.45-3.15 (m, 3 H), 3.15-2.97 (m, 3H), 2.92 (s, 3H), 2.87-2.56 (m, 3H), 2.12-1.96 (m, 2H), 1.18 (d, *J* = 1.9 Hz, 9H), 1.10 (d, *J* = 6.5 Hz, 1.5H), 1.02 (d, *J* = 6.3 Hz, 1.5H); HRMS (ESI) m/z: [M + H]+ for $C_{39}H_{49}N_8O_3^+$ calculated 677.3922 found 677.3918.

*Synthesis of (S)-1-(4-(3-Chloro-2,2-dimethylpropanoyl)-3-methylpiperazin-1-yl)-4-(3-(6-(4-methylpiperazin-1-yl)-1H,3'H-[2,5'-bibenzo[d]imidazol]-2'-yl)phenoxy)butan-1-one (**20**)*

Compound **20-p1** (21 mg, 30 μmol, 1 eq) was stirred for 1 h at rt with 2 N HCl in 50:50 dioxane:DCM (2 mL). Once reaction was confirmed complete by LC/MS, solvent was removed under reduced pressure. The crude product was dissolved in DCM (0.6 mL) and stirred at 0 °C in an ice bath. To this solution, triethylamine (17 mg, 170 μmol, ~6 eq) and 3-chloropivaloyl chloride (CAS: 4300-97-4, 5.3 mg, 34 μmol, 1.1 eq) were added, the solution was brought to rt, and then the reaction was stirred for 1 h. The product was purified by RP-HPLC (60 min, 40-100% methanol:water + 0.1% (v/v) TFA). Fractions were lyophilized to afford a yellow solid, **20** (14.0 mg, 66% yield).

$^1$H NMR (DMSO-d6, 400MHz): δ 10.18 (b, 1H), 8.51 (s, 1H), 8.08 (dd, *J* = 8.5, 1.5 Hz, 1H), 7.92 (d, *J* = 8.5 Hz, 1H), 7.85 (d, *J* = 7.7 Hz, 1H), 7.83 (s, 1H), 7.73 (d, *J* = 9.0 Hz, 1H), 7.52 (t, *J* = 7.8 Hz, 1H), 7.34 (dd, *J* = 9.1, 2.0 Hz, 1H), 7.25 (d, *J* = 1.9 Hz, 1H), 7.19-7.13 (m, 1 H), 4.59-4.48 (m, 1H), 4.33-4.02 (m, 4H), 4.02-3.67 (m, 5H), 3.67-3.46 (m, 2H), 3.46-3.17 (m, 3H), 3.17-2.98 (m, 3H), 2.92 (s, 3H), 2.87-2.58 (m, 3H), 2.11-1.97 (m, 2H), 1.28 (d, *J* = 5.0 Hz, 6H), 1.11 (d, *J* = 6.6 Hz, 1.5H), 1.04 (d, *J* = 6.2 Hz, 1.5H); HRMS (ESI) m/z: [M + H]+ for $C_{39}H_{48}ClN_8O_3^+$ calculated 711.3532 found 711.3530.

### COMPOUND CHARACTERIZATION

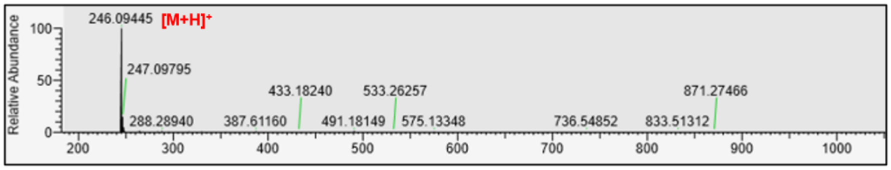


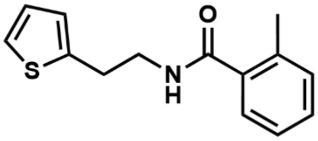


**10-p1**

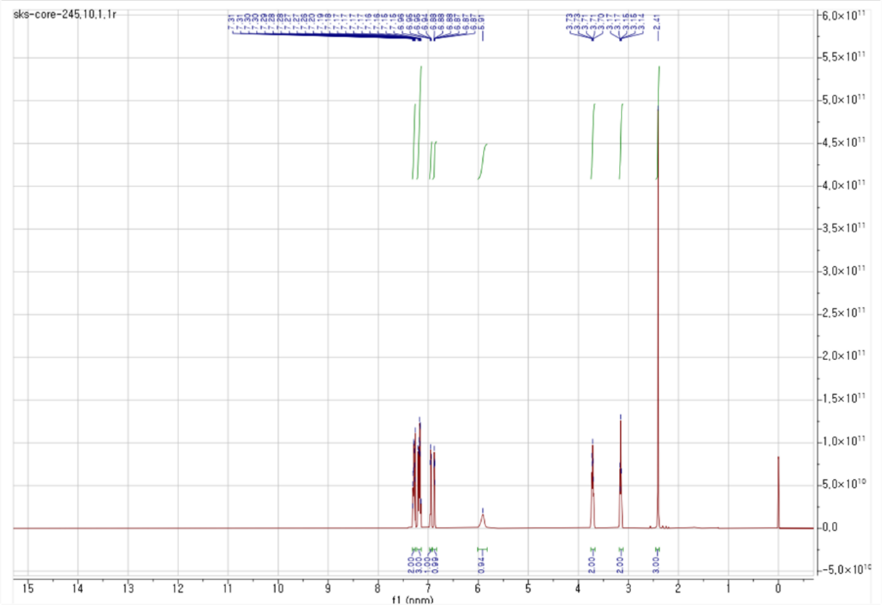


$^{1}H$ NMR, 400 MHz, $CDCl_3$

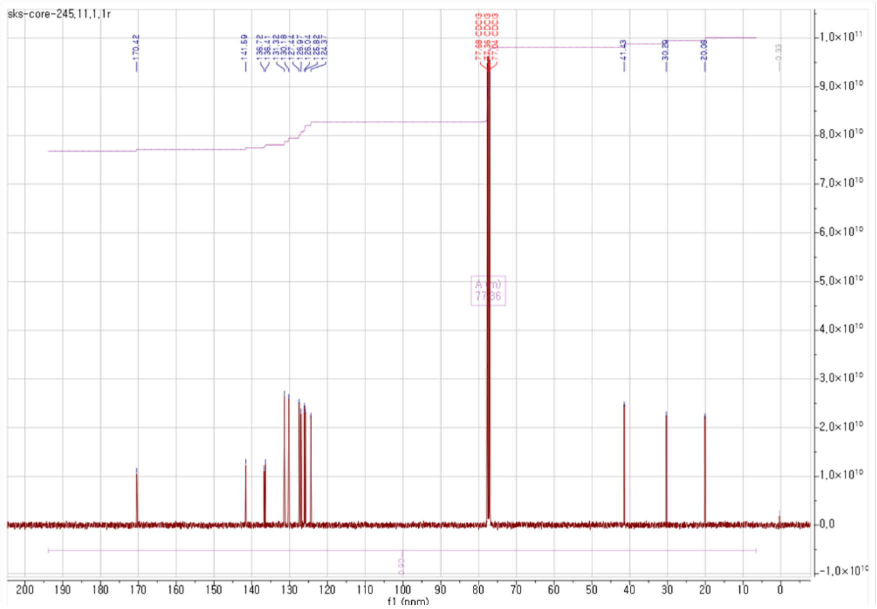


$^{13}C$ NMR, 100 MHz, $CDCl_3$

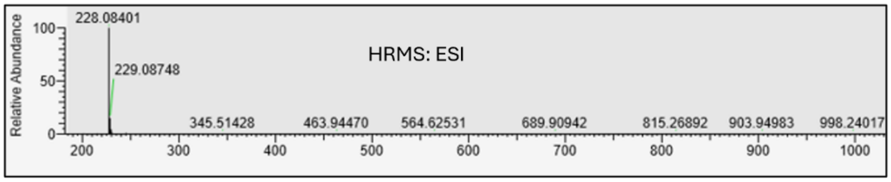


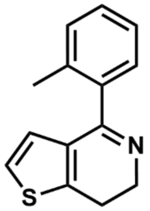


**10-p2**

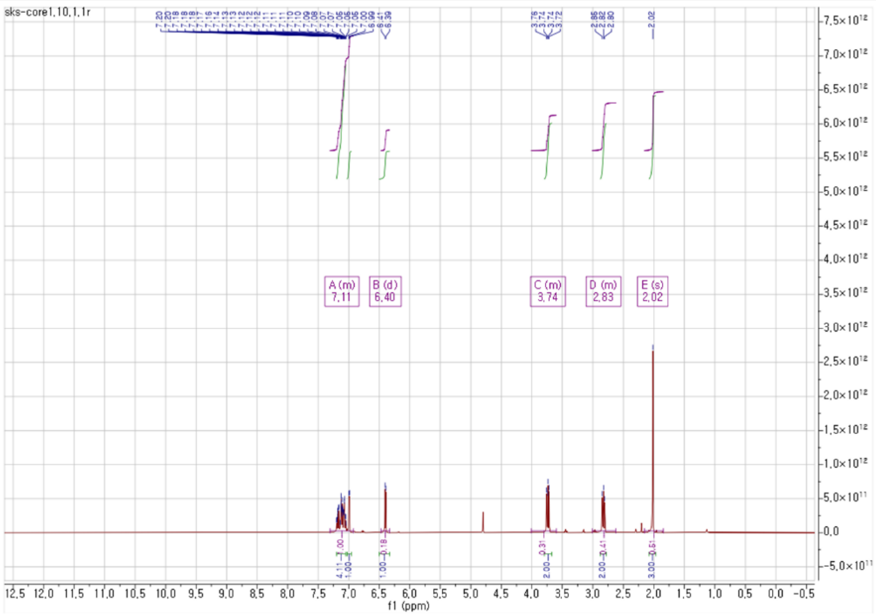


$^{1}H$ NMR, 400 MHz, $CDCl_3$

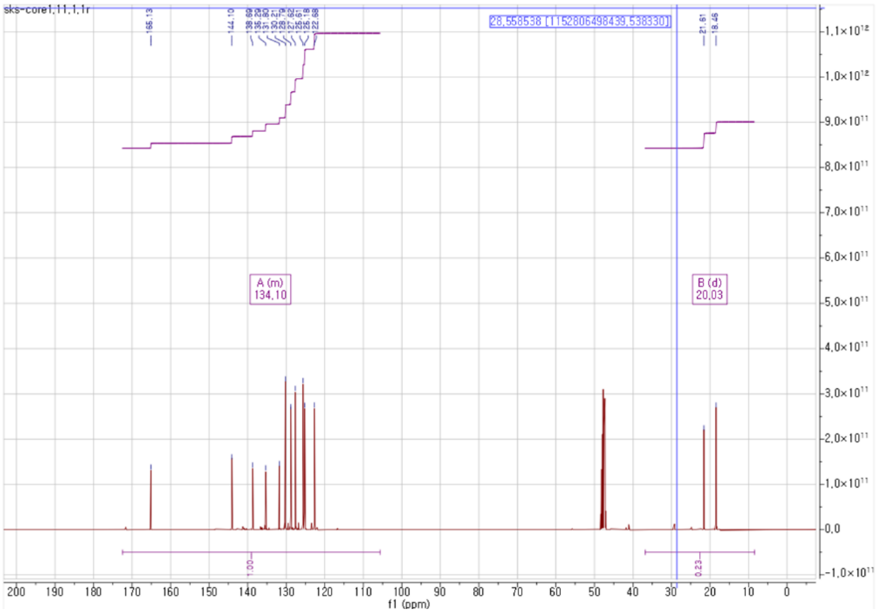


$^{13}C$ NMR, 100 MHz, $CDCl_3$

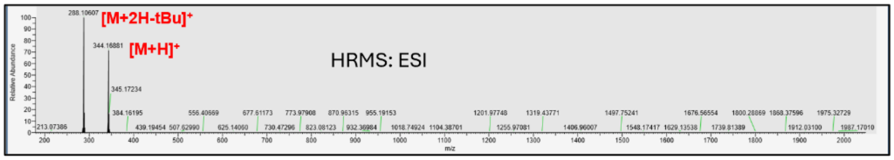


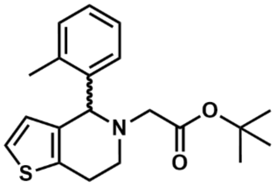

**10-p3**

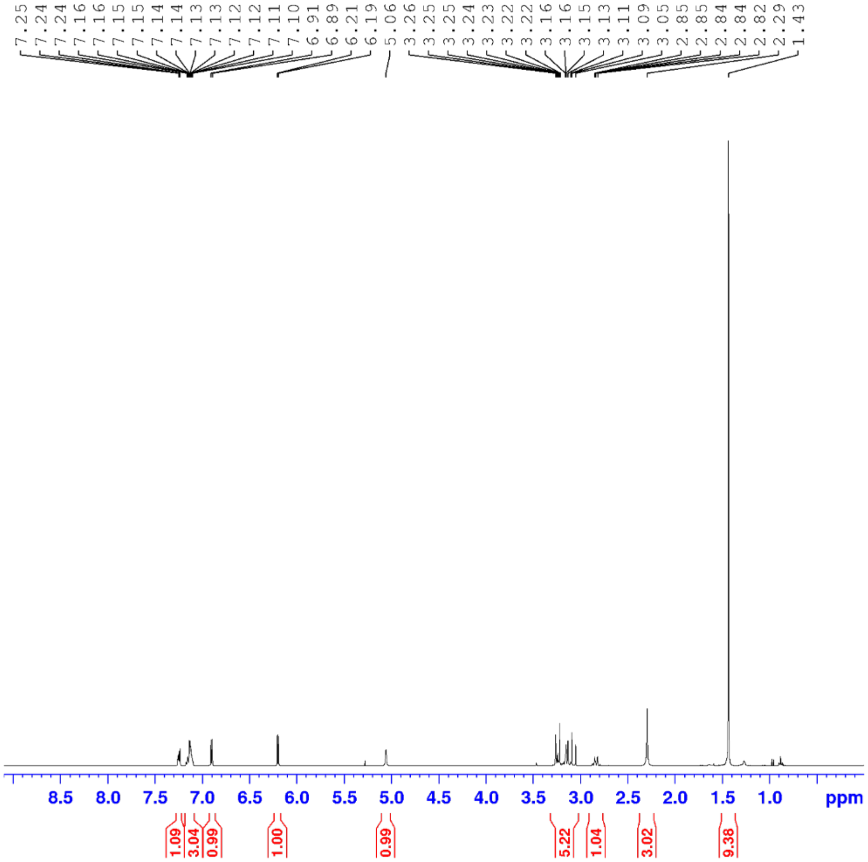


$^1$H NMR, 400 MHz, $CDCl_3$

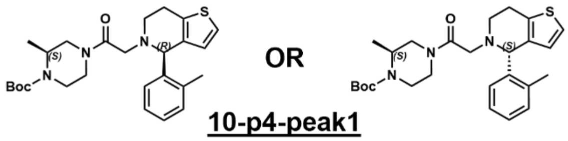


**10-p4-peak1**

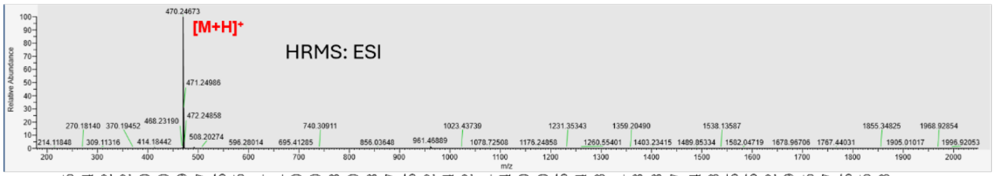


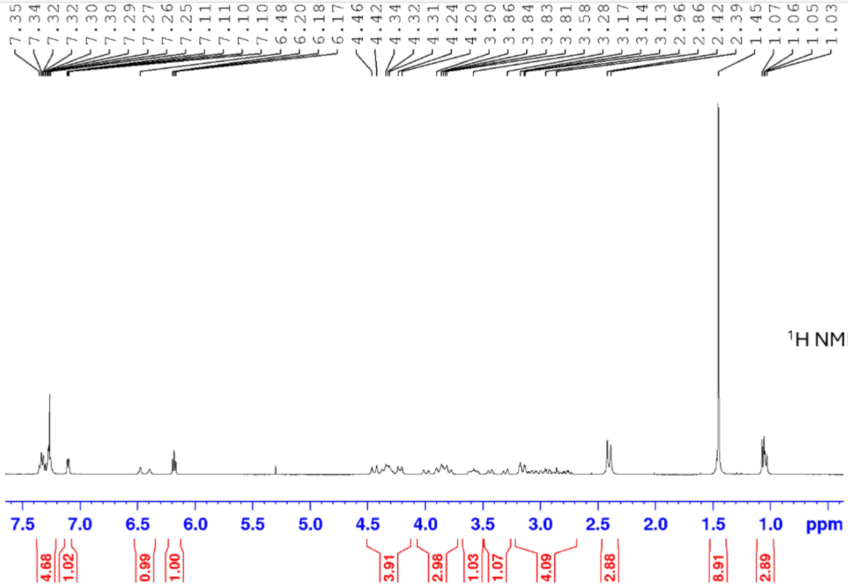

$^1$H NMR, 400 MHz, $CDCl_3$

**10-p4-peak2**

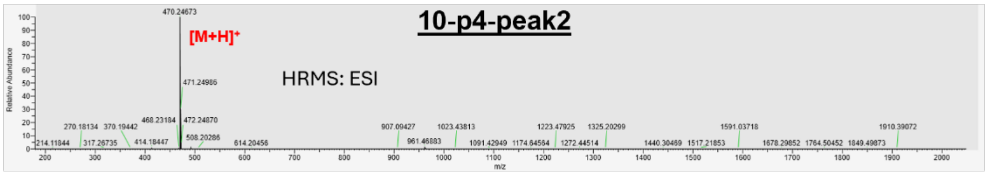


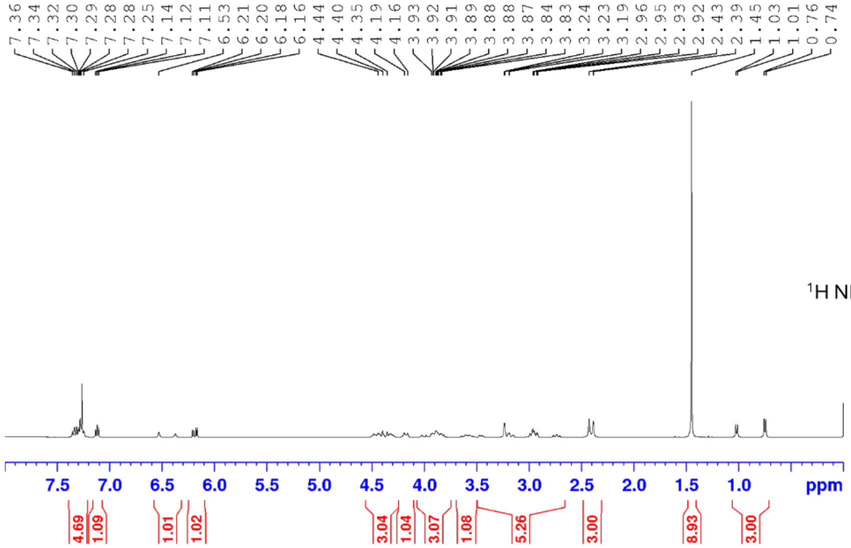

$^1$H NMR, 400 MHz, $CDCl_3$

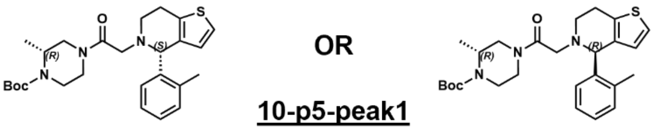


### 10-p5-peak1

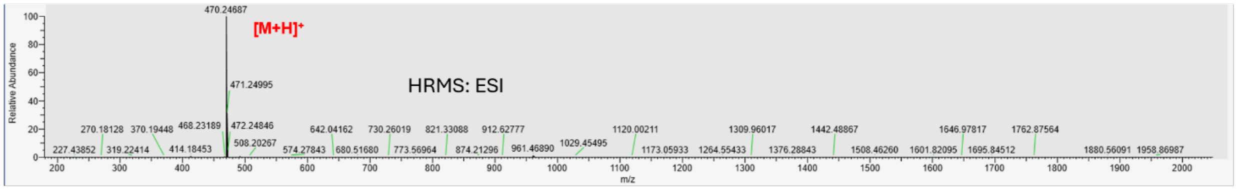


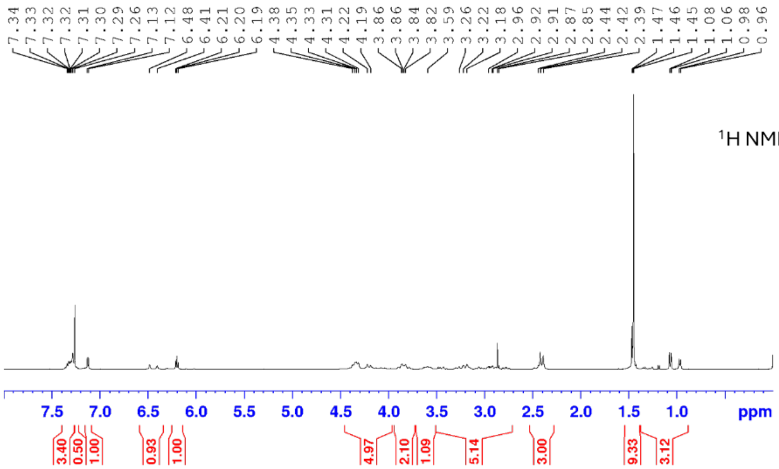


### 10-p5-peak2

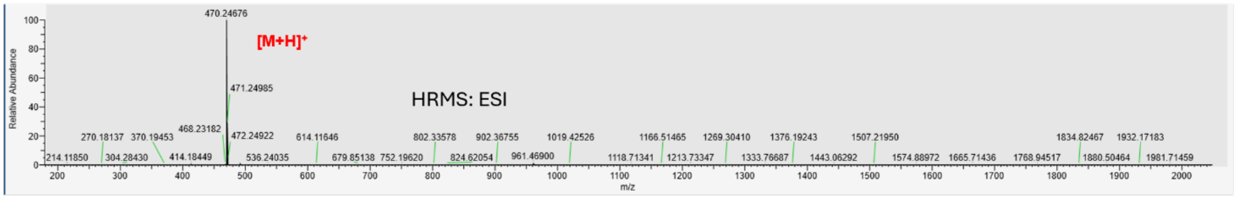


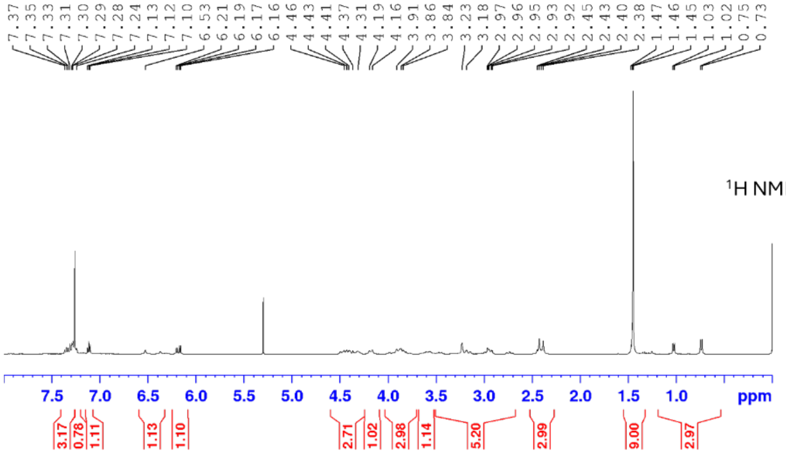

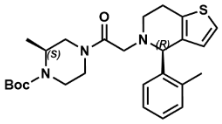

**AND**

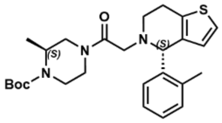

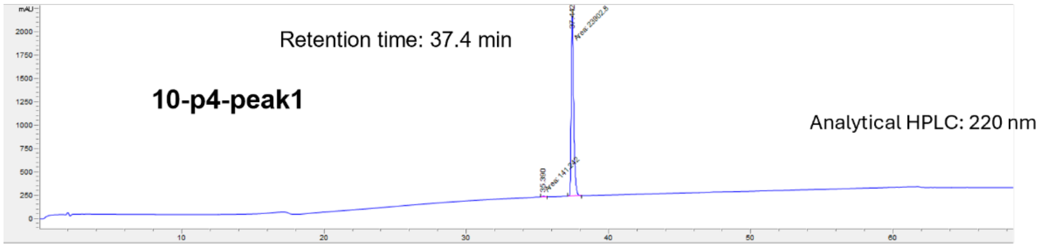

Retention time: 37.4 min
10-p4-peak1
Analytical HPLC: 220 nm


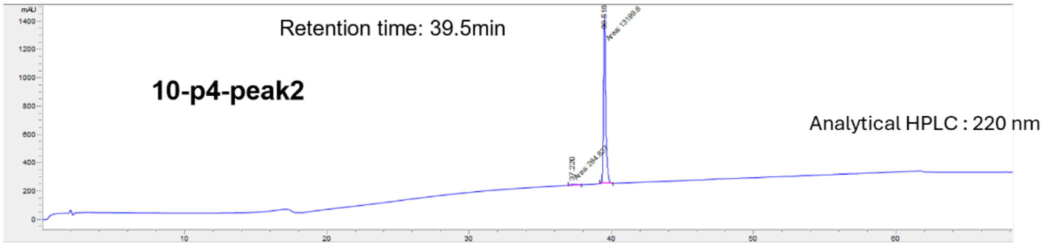

Retention time: 39.5min
10-p4-peak2
Analytical HPLC : 220 nm


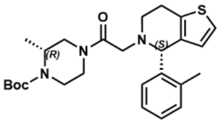

**AND**

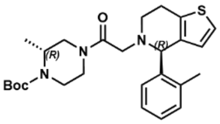

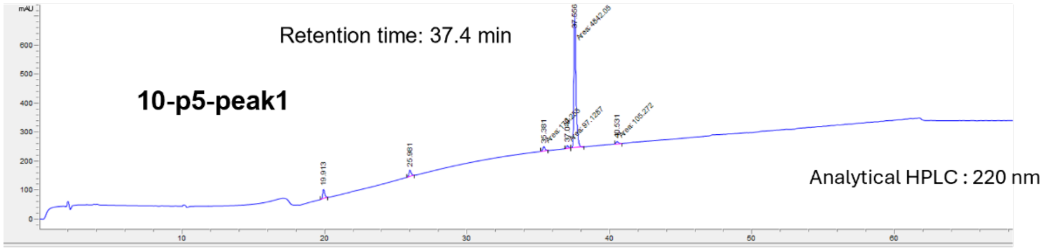

Retention time: 37.4 min
10-p5-peak1
Analytical HPLC : 220 nm


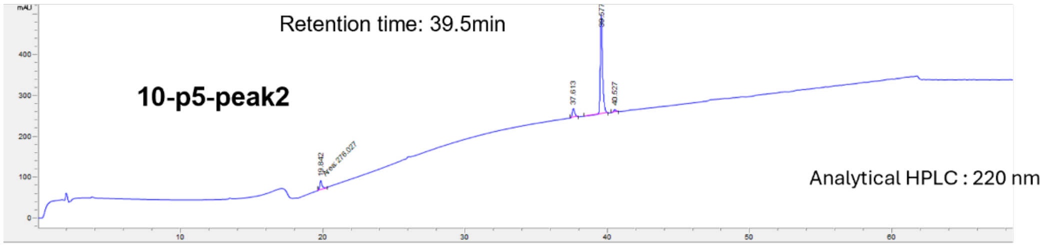

Retention time: 39.5min
10-p5-peak2
Analytical HPLC : 220 nm

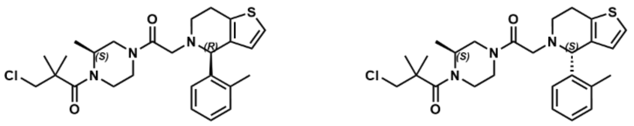

**10a and 10b**

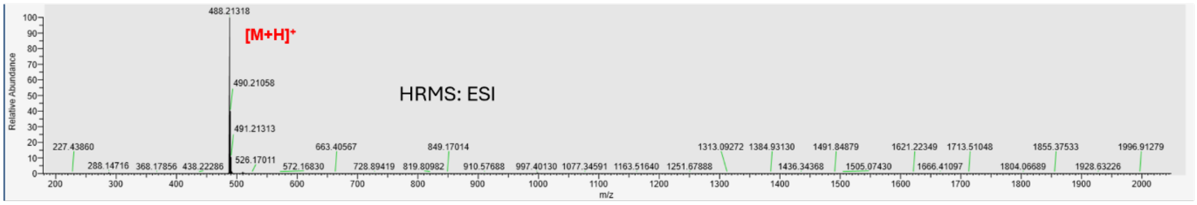
488.21318
[M+H]+
490.21058
491.21313
HRMS: ESI
Relative Abundance
m/z

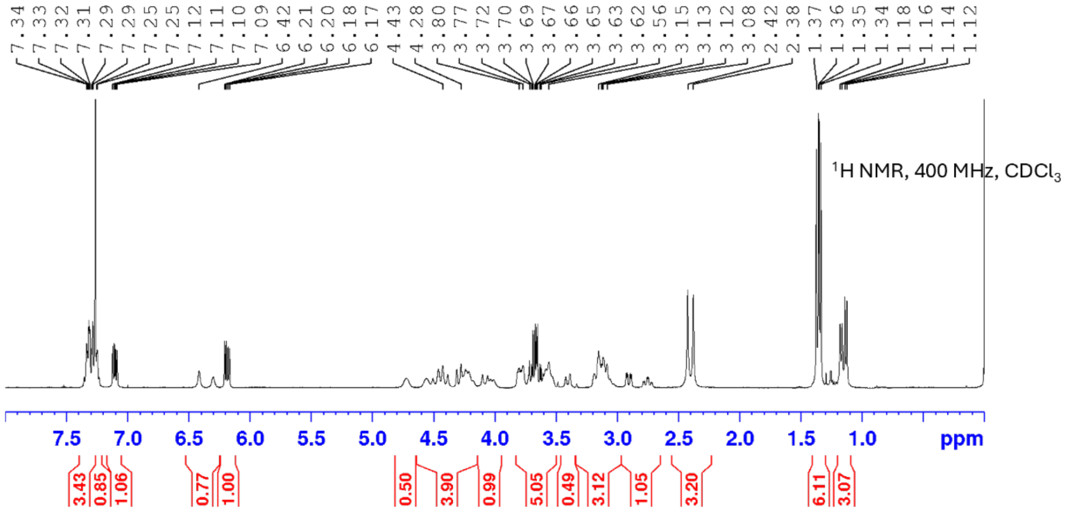
1H NMR, 400 MHz, CDCl3
ppm

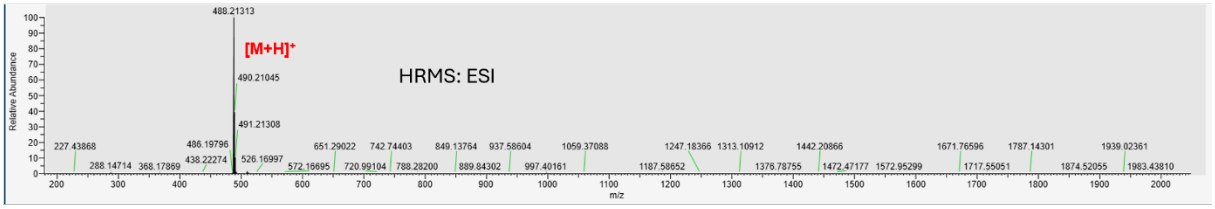
488.21313
[M+H]+
490.21045
491.21308
HRMS: ESI
Relative Abundance
m/z

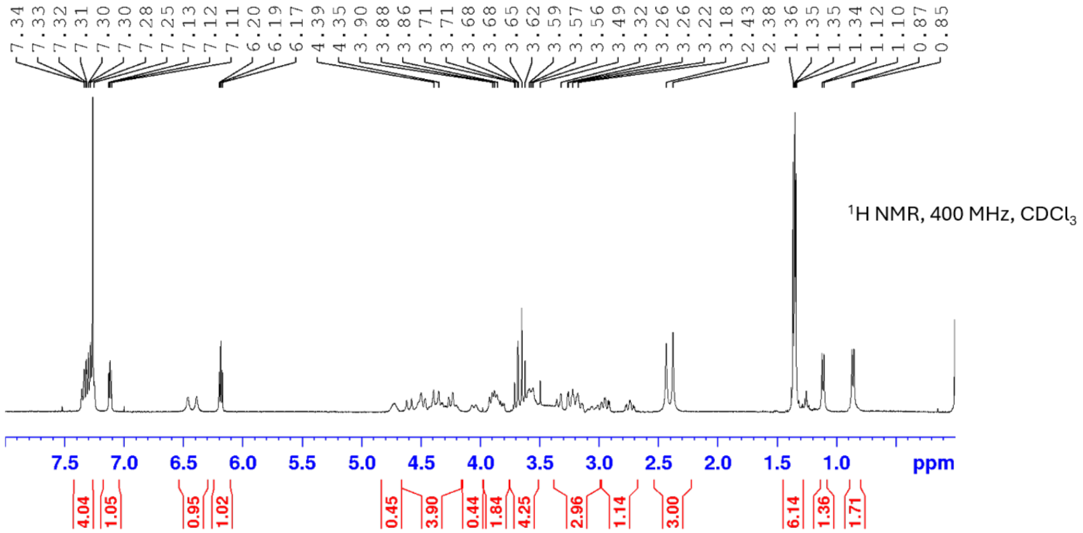
1H NMR, 400 MHz, CDCl3
ppm

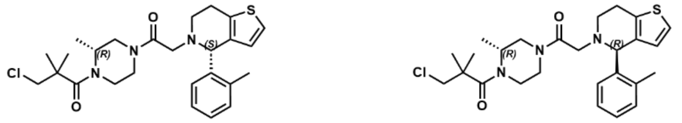
Cl
N
O
S
(R)
(S)
(R)
(R)

**10c and 10d**

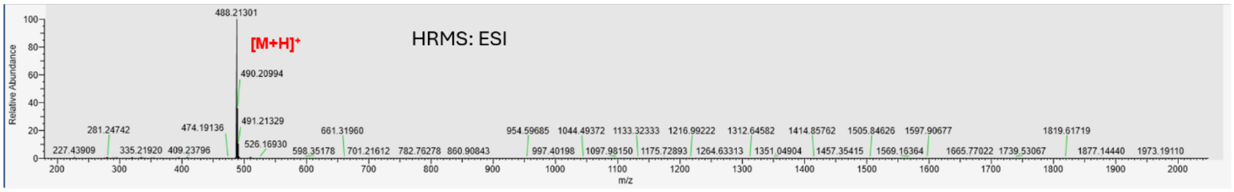
HRMS: ESI
[M+H]+
488.21301
490.20994
491.21329
526.16930
474.19136
281.24742
227.43909
335.21920
409.23796
598.35178
661.31960
701.21612
782.76278
860.90843
954.59685
997.40198
1044.49372
1097.98150
1133.32333
1175.72893
1216.99222
1264.63313
1312.64582
1351.04904
1414.85762
1457.35415
1505.84626
1569.16364
1597.90677
1665.77022
1739.53067
1819.61719
1877.14440
1973.19110
Relative Abundance
m/z

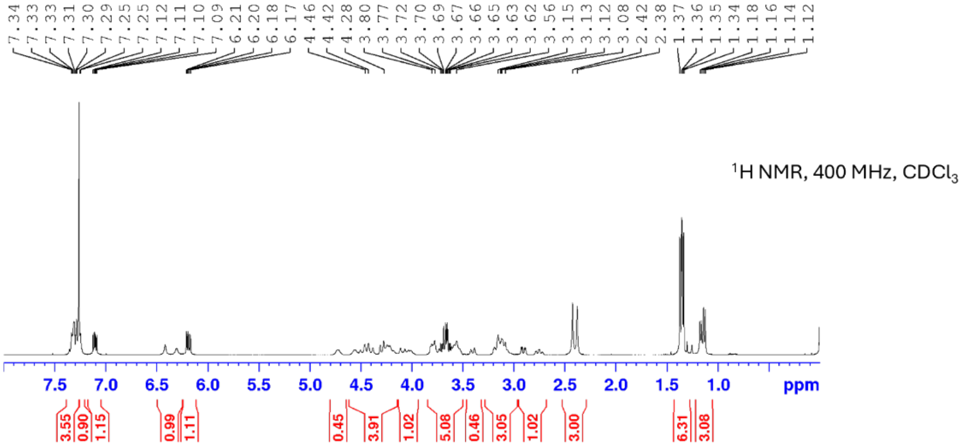
7.34 7.33 7.33 7.31 7.30 7.29 7.25 7.25 7.12 7.11 7.10 7.09 6.21 6.20 6.18 6.17 4.46 4.42 4.28 3.80 3.77 3.72 3.70 3.69 3.67 3.66 3.65 3.63 3.62 3.56 3.15 3.13 3.12 3.08 2.42 2.38 1.37 1.36 1.35 1.34 1.18 1.16 1.14 1.12
1H NMR, 400 MHz, CDCl3
ppm
3.55 0.90 1.15 0.99 1.11 0.45 3.91 1.02 5.08 0.46 3.05 1.02 3.00 6.31 3.08

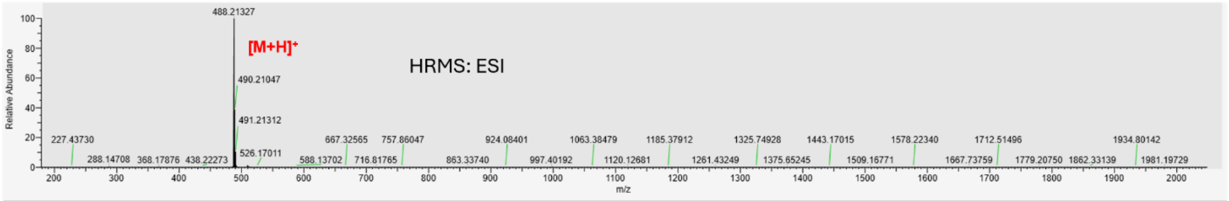
HRMS: ESI
[M+H]+
488.21327
490.21047
491.21312
526.17011
227.43730
288.14708
368.17876
438.22273
588.13702
667.32565
716.81765
757.86047
863.33740
924.08401
997.40192
1063.38479
1120.12681
1185.37912
1261.43249
1325.74928
1375.65245
1443.17015
1509.16771
1578.22340
1667.73759
1712.51496
1779.20750
1862.33139
1934.80142
1981.19729
Relative Abundance
m/z

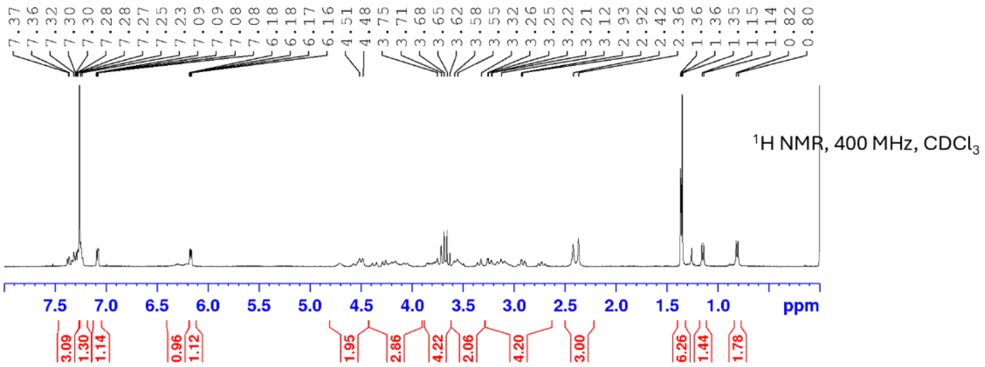
7.37 7.36 7.32 7.30 7.30 7.28 7.27 7.25 7.23 7.09 7.09 7.08 7.08 6.18 6.17 6.16 4.51 4.48 3.75 3.71 3.68 3.65 3.62 3.58 3.55 3.32 3.26 3.25 3.22 3.21 3.12 2.93 2.92 2.42 2.36 1.36 1.36 1.35 1.15 1.14 0.82 0.80
1H NMR, 400 MHz, CDCl3
ppm
3.09 1.30 1.14 0.96 1.12 1.95 2.86 4.22 2.06 4.20 3.00 6.26 1.44 1.78

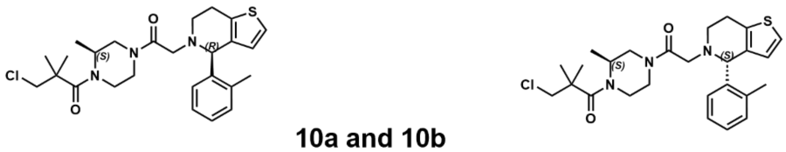


**10a and 10b**

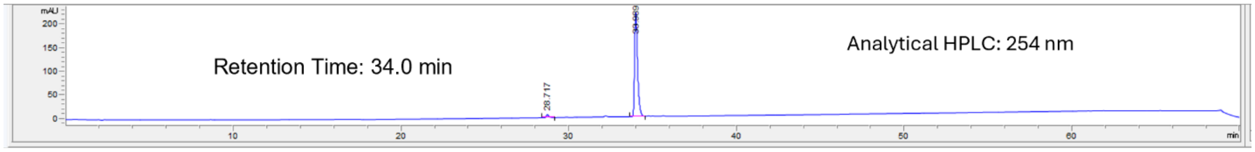


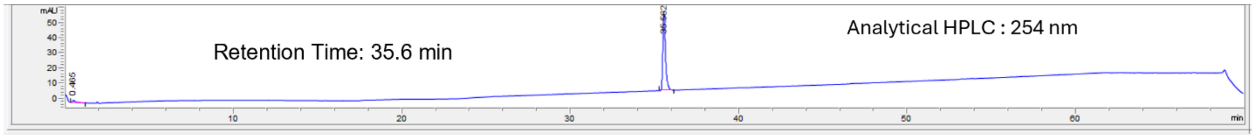


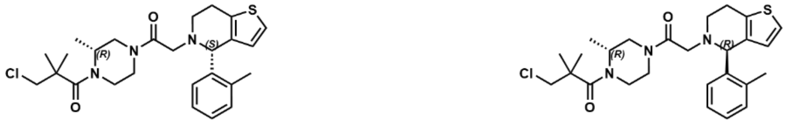


**10c and 10d**

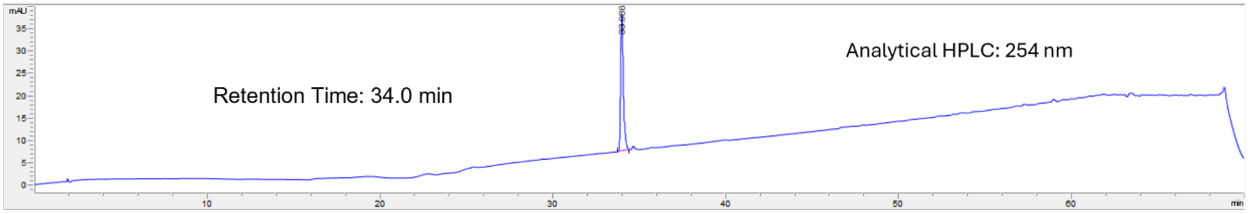


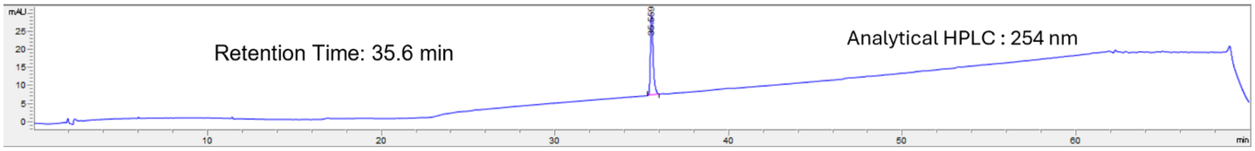

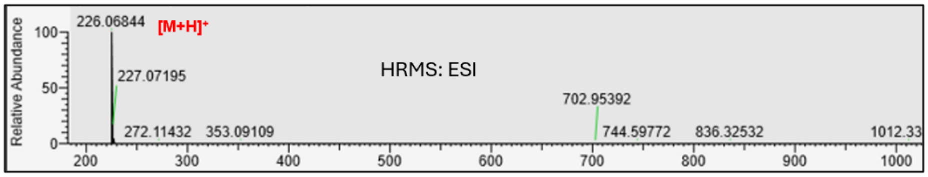


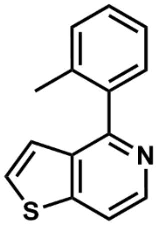

**10-p6**

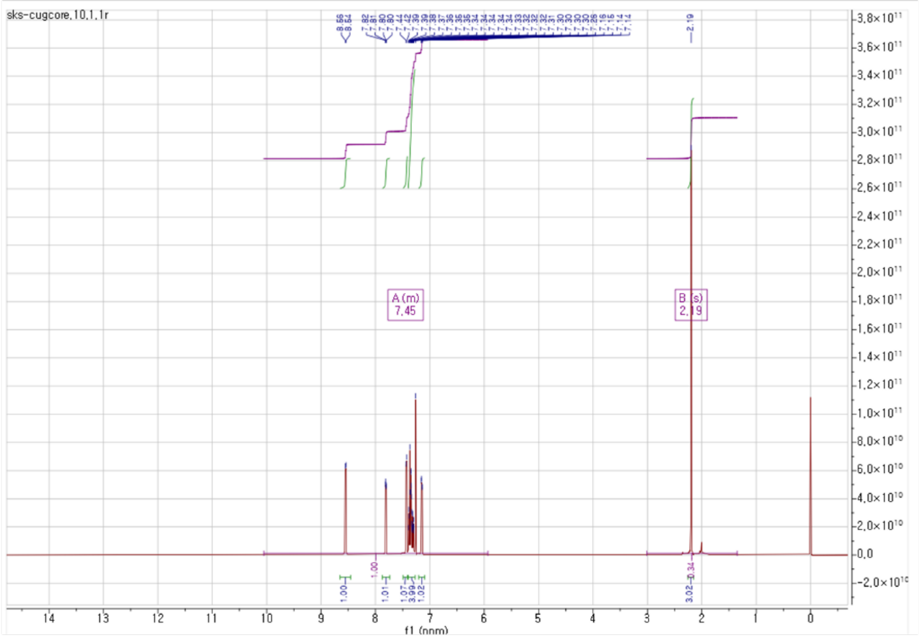


$^{1}H$ NMR, 400 MHz, $CDCl_3$

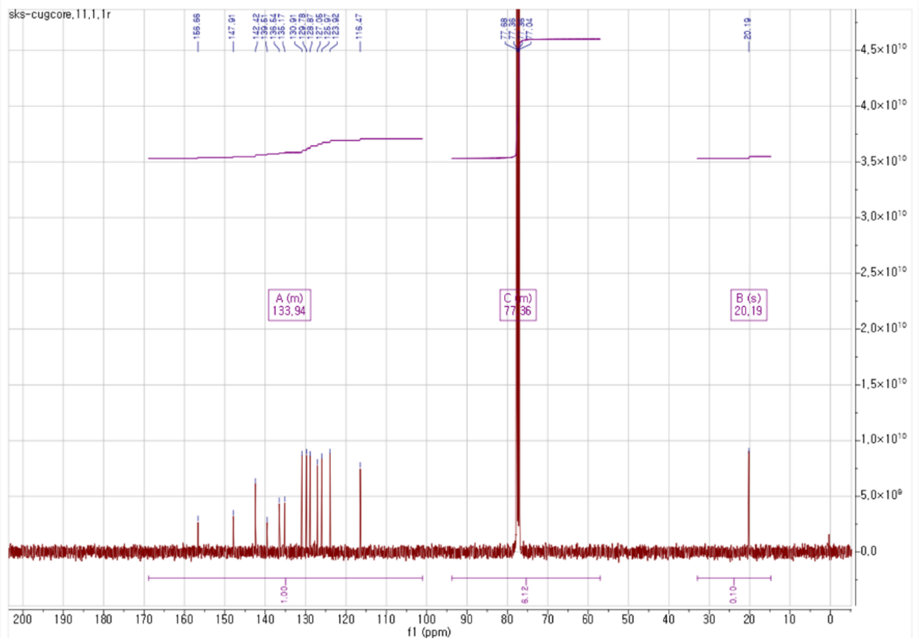


$^{13}C$ NMR, 100 MHz, $CDCl_3$

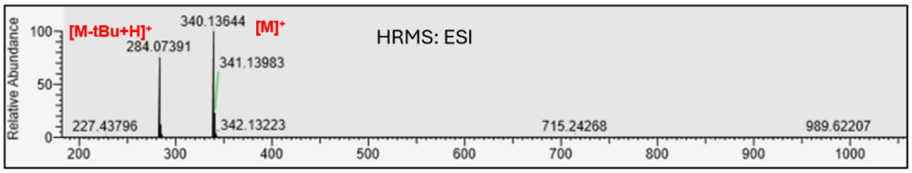


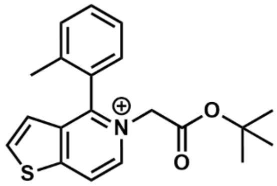

**10-p7**

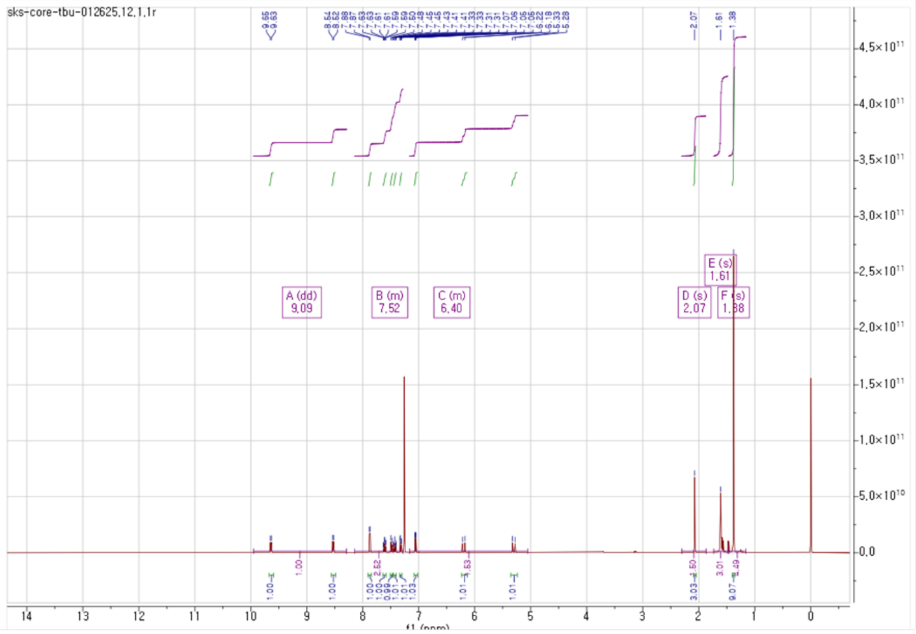

$^1$H NMR, 400 MHz, $CDCl_3$

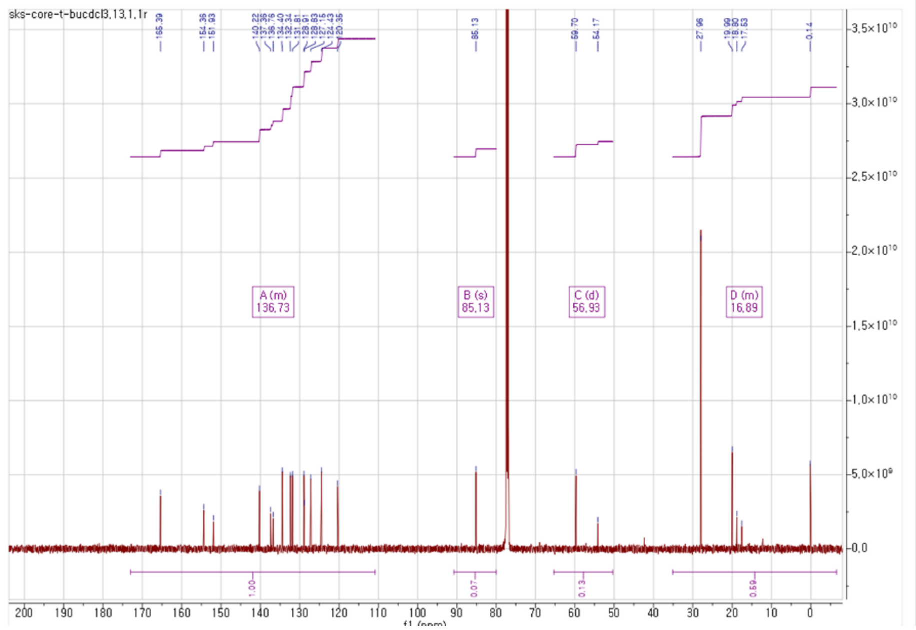

$^{13}$C NMR, 150 MHz, $CDCl_3$

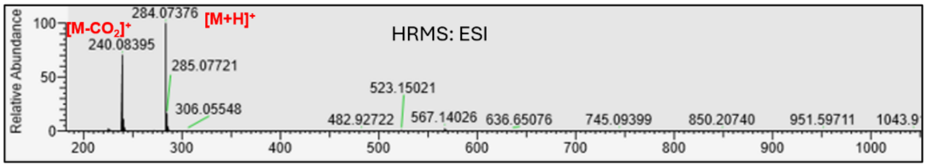


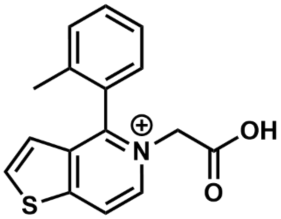

**10-p8**

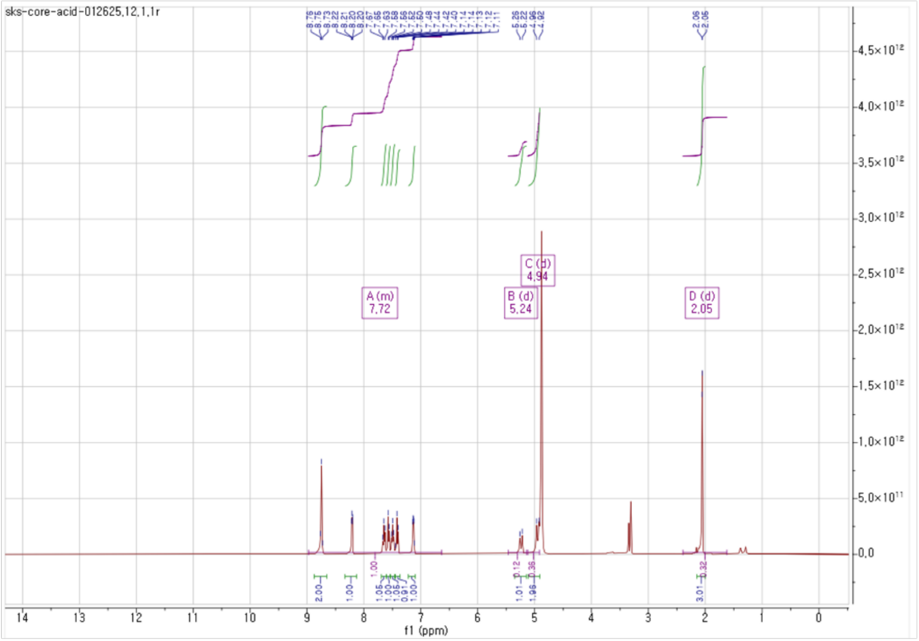

$^1$H NMR, 400 MHz, $CD_3OD$

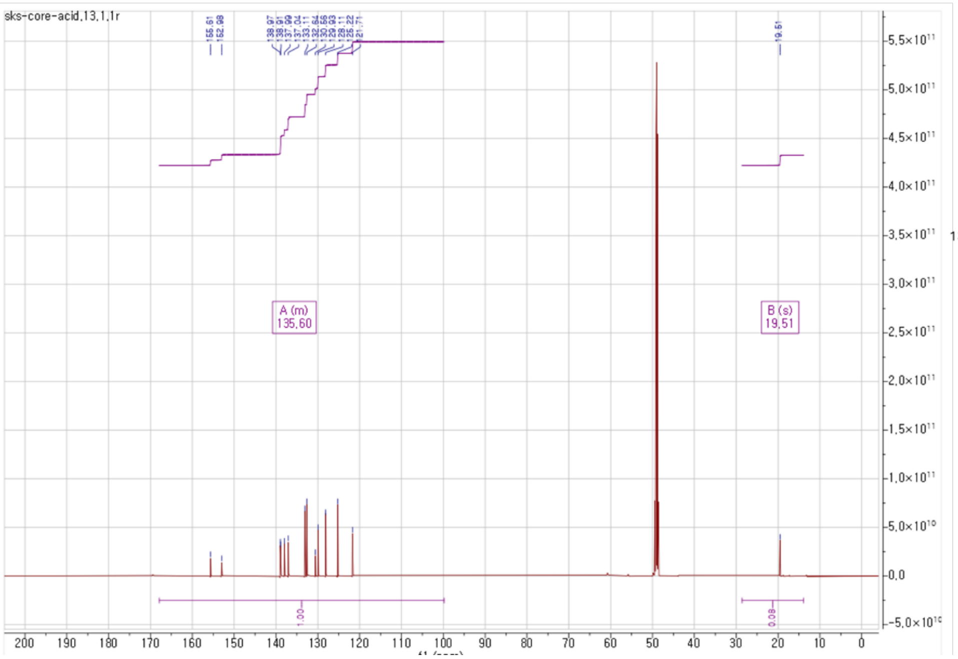

$^{13}$C NMR, 150 MHz, $CD_3OD$

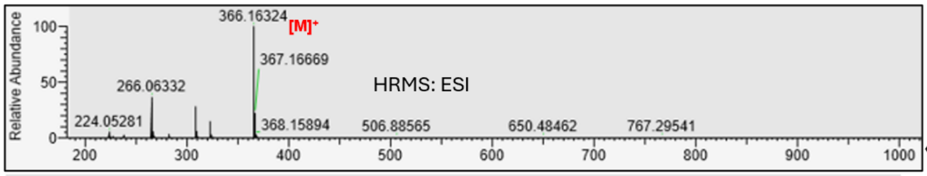


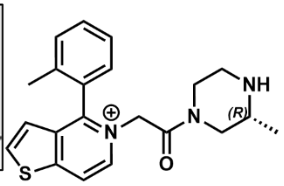


**10-p9**

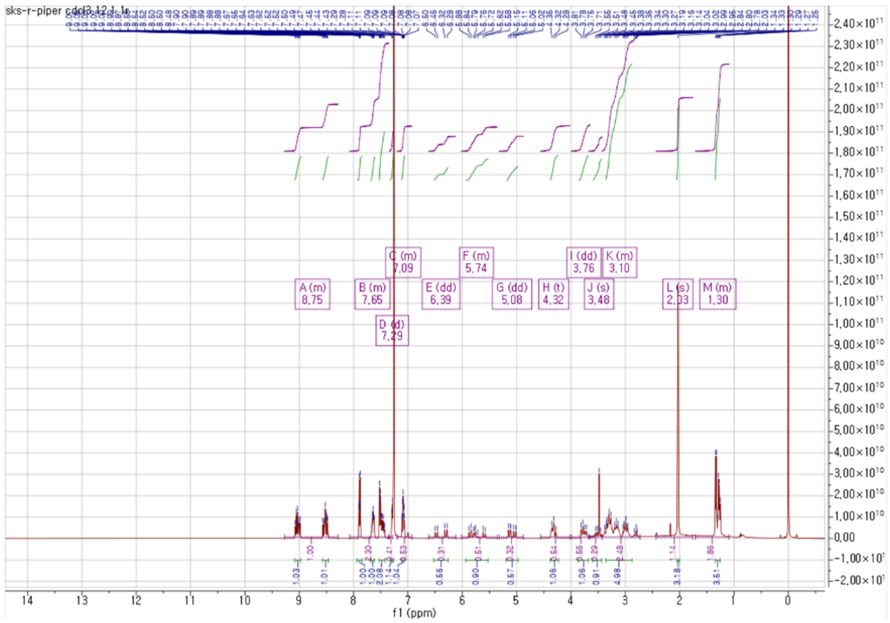


$^{1}H$ NMR, 400 MHz, $CDCl_3$

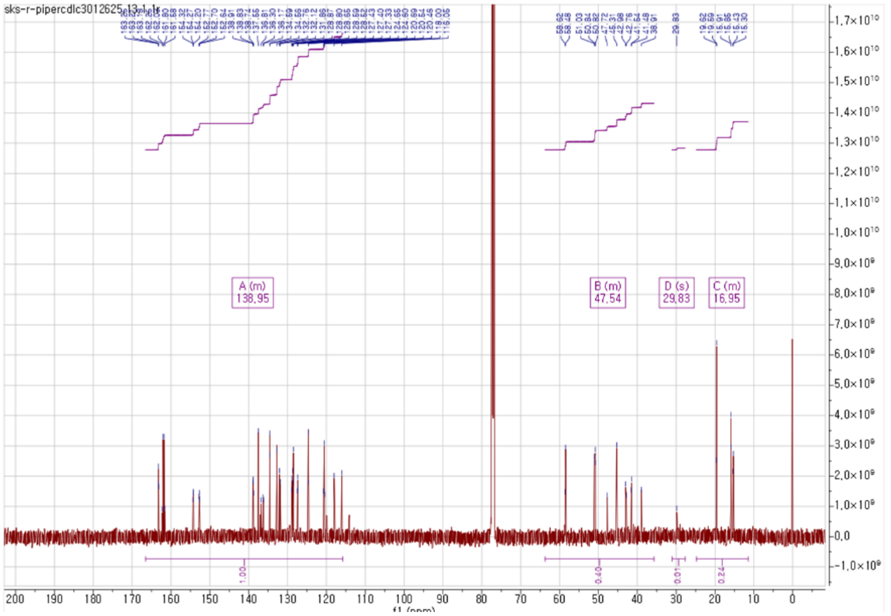


$^{13}C$ NMR, 150 MHz, $CDCl_3$

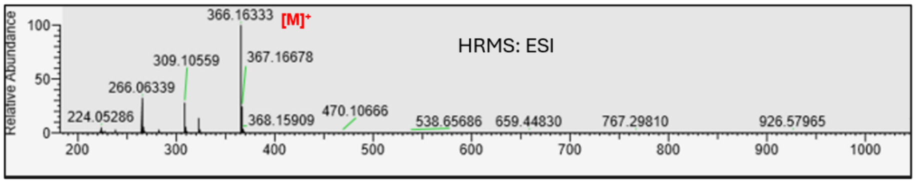


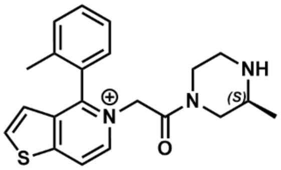


**10-p10**

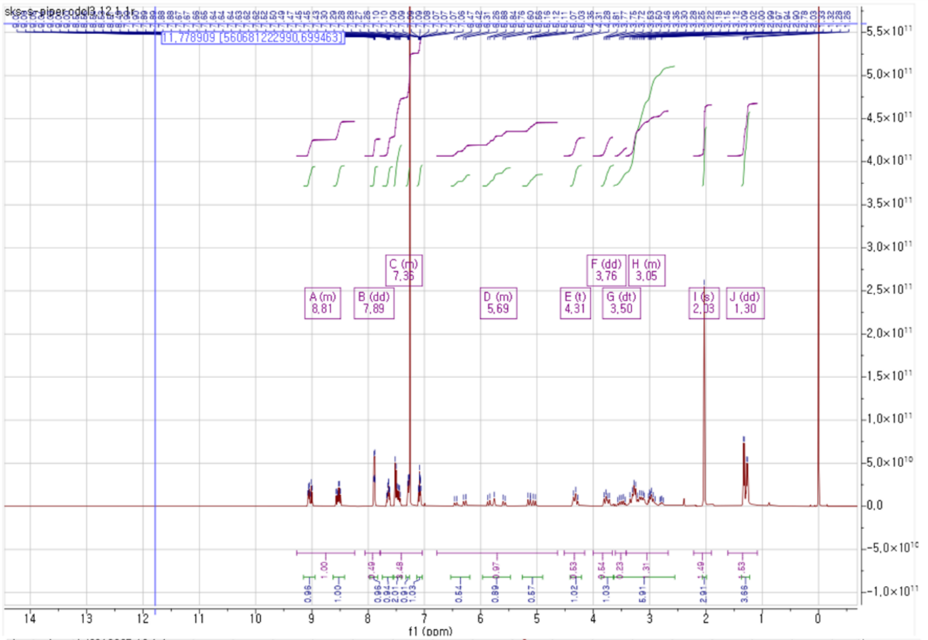


$^{1}H$ NMR, 400 MHz, $CDCl_3$

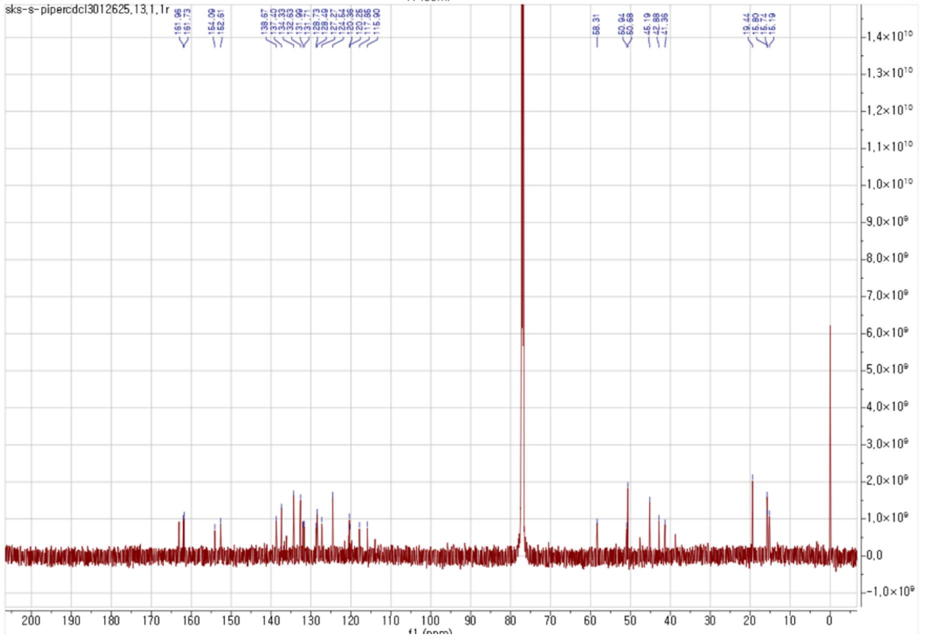


$^{13}C$ NMR, 150 MHz, $CDCl_3$

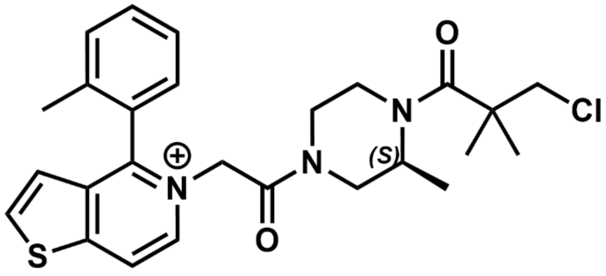

O
Cl
N
(S)
N
O
S


**10g**

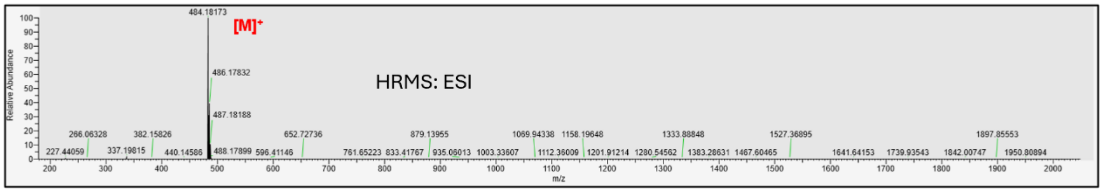

484.18173
[M]+
HRMS: ESI
486.17832
487.18188
Relative Abundance
m/z


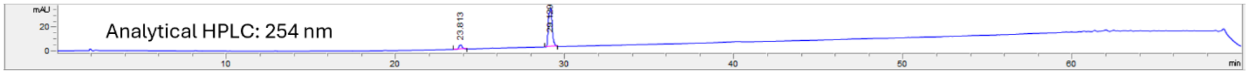

mAU
Analytical HPLC: 254 nm
23.813
min


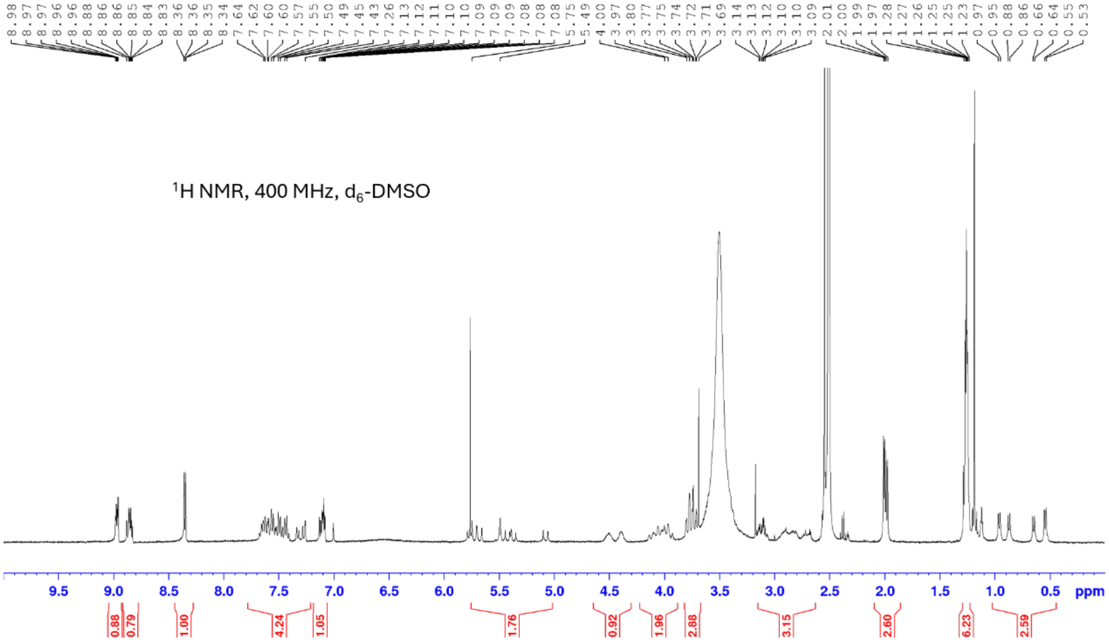

1H NMR, 400 MHz, d6-DMSO
ppm

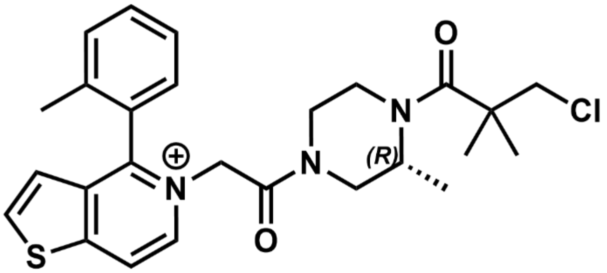

**10h**

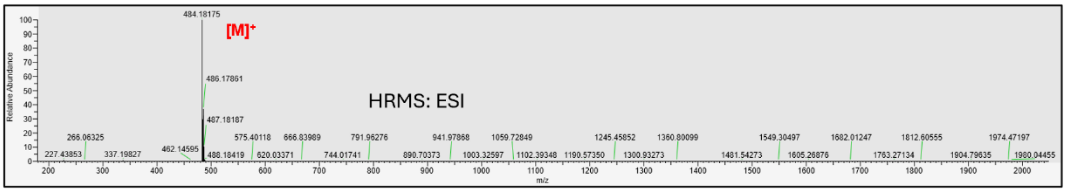

484.18175
[M]+
486.17861
HRMS: ESI
487.18187
Relative Abundance
m/z


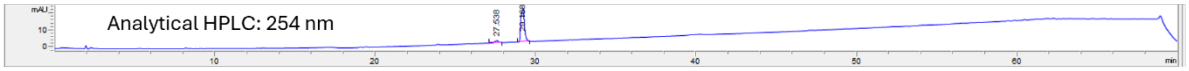

Analytical HPLC: 254 nm
mAU
27.538
min


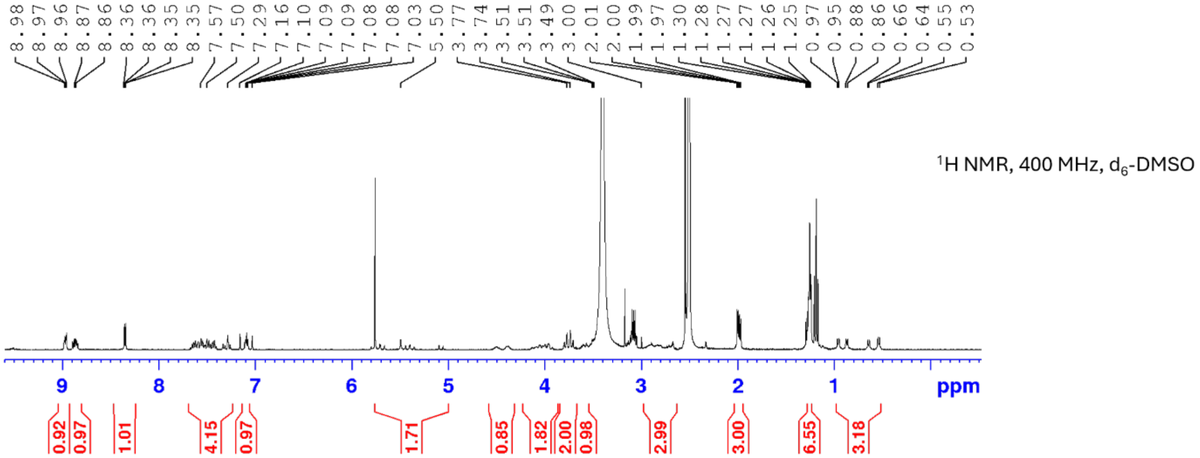

1H NMR, 400 MHz, d6-DMSO
ppm

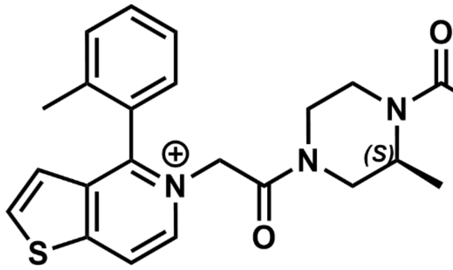

**10i**

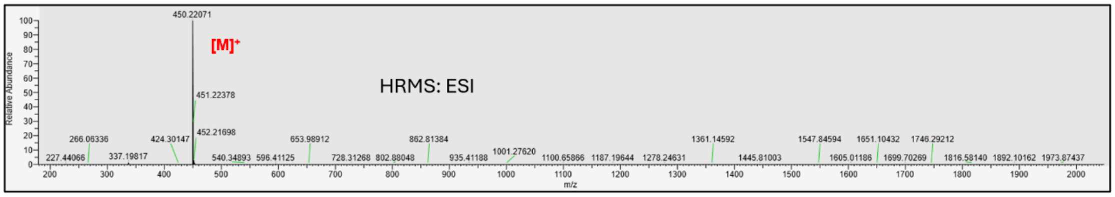


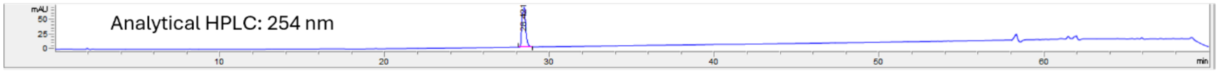


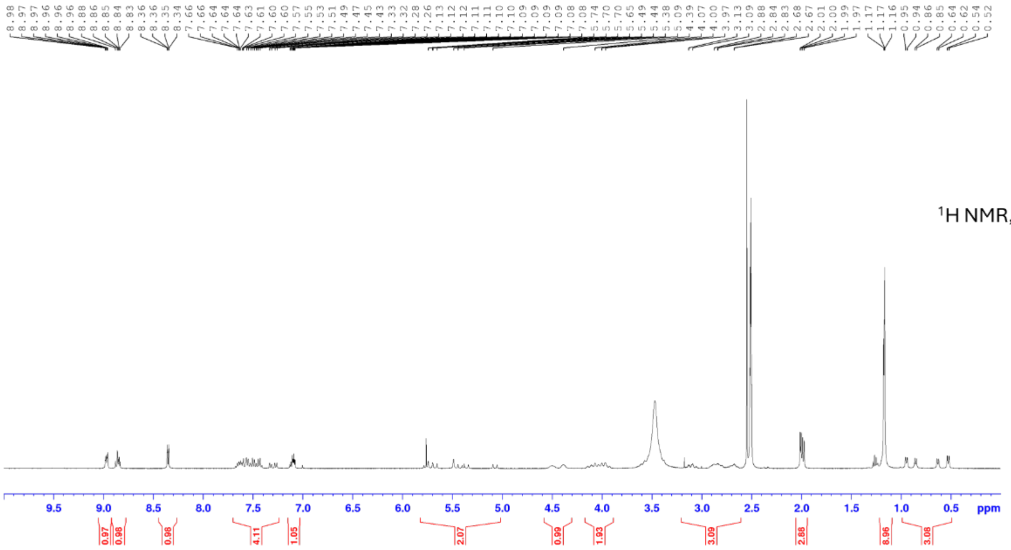


$^1$H NMR, 400 MHz, $d_6$-DMSO

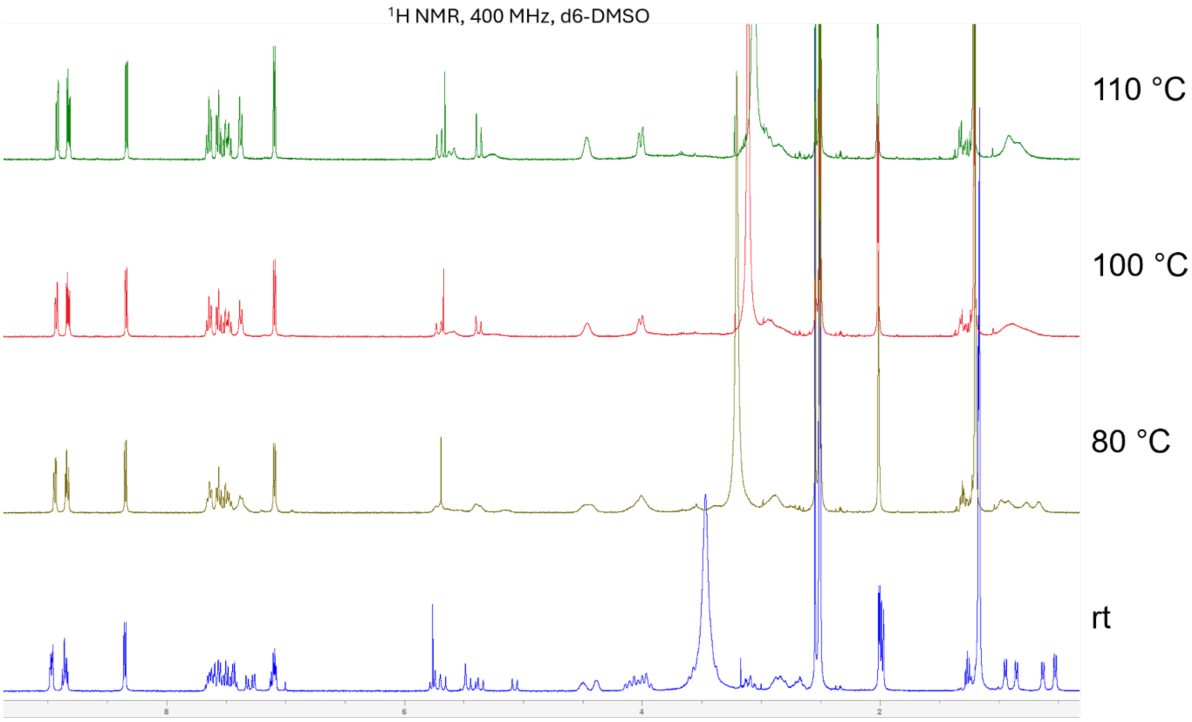


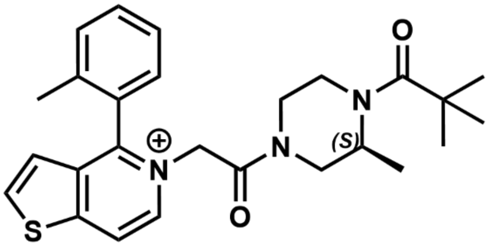


**10i**

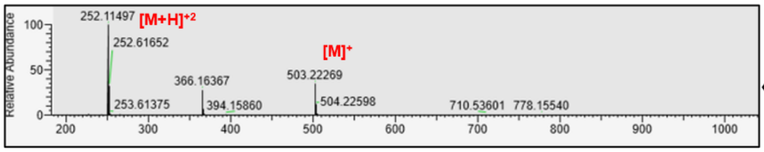


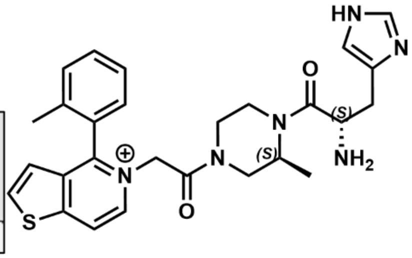


**10j**

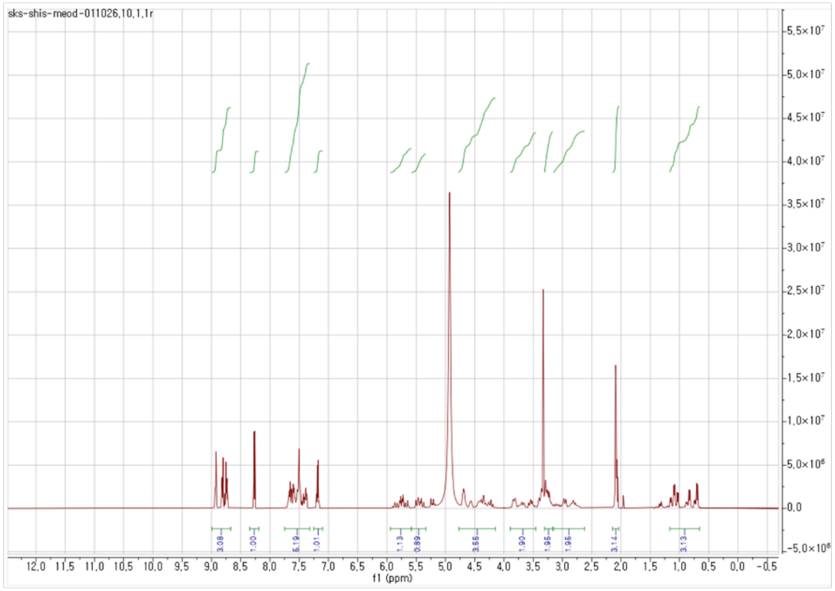


$^{1}$H NMR, 400 MHz, $CD_3OD$

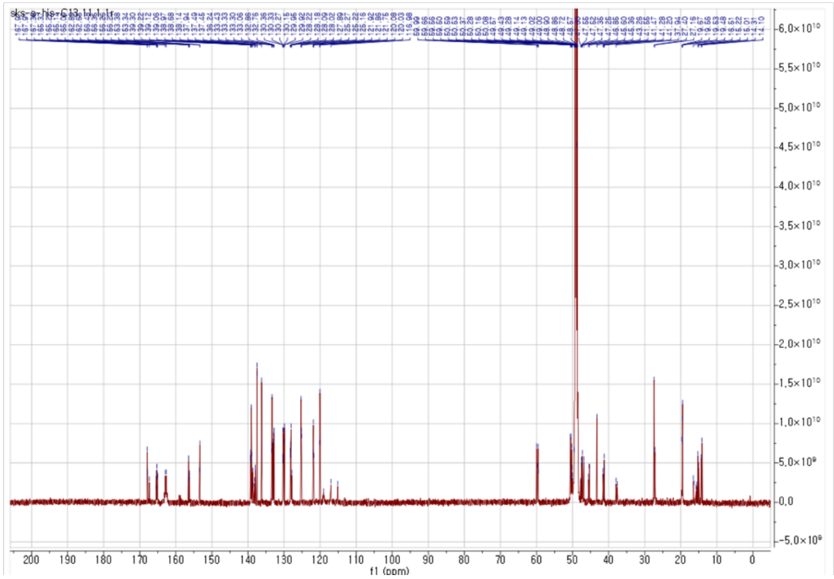


$^{13}$C NMR, 150 MHz, $CD_3OD$

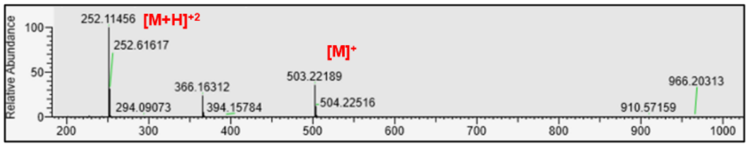


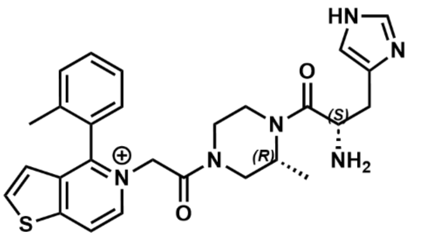


**10k**

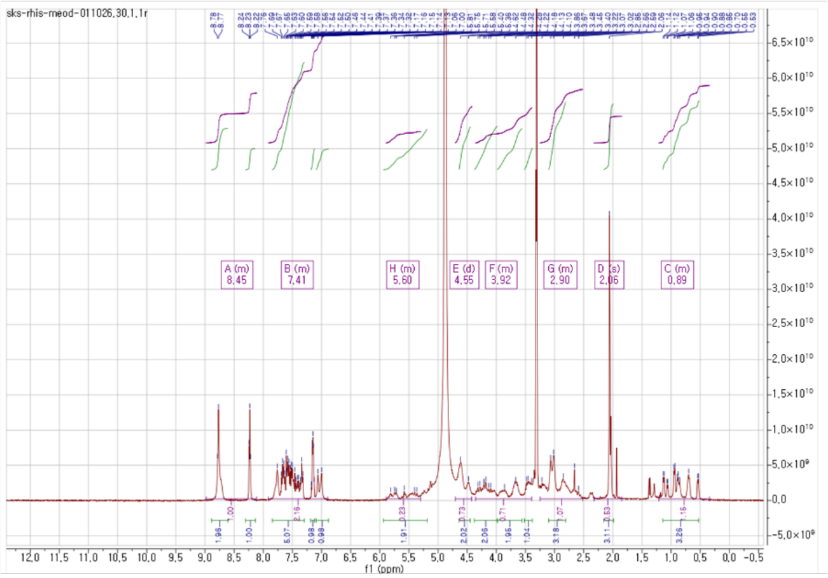


$^{1}$H NMR, 400 MHz, $CD_3OD$

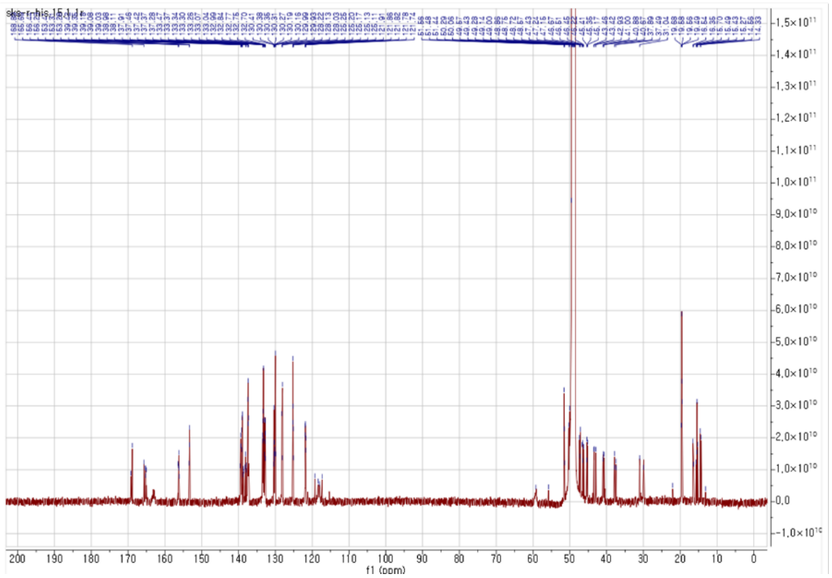


$^{13}$C NMR, 150 MHz, $CD_3OD$

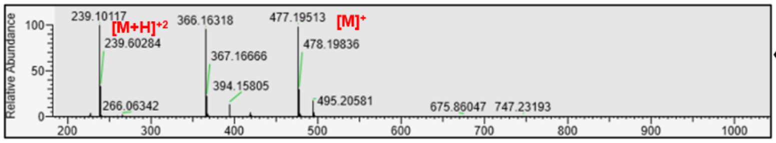


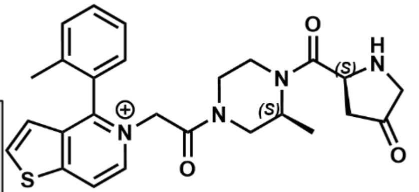

**10l**

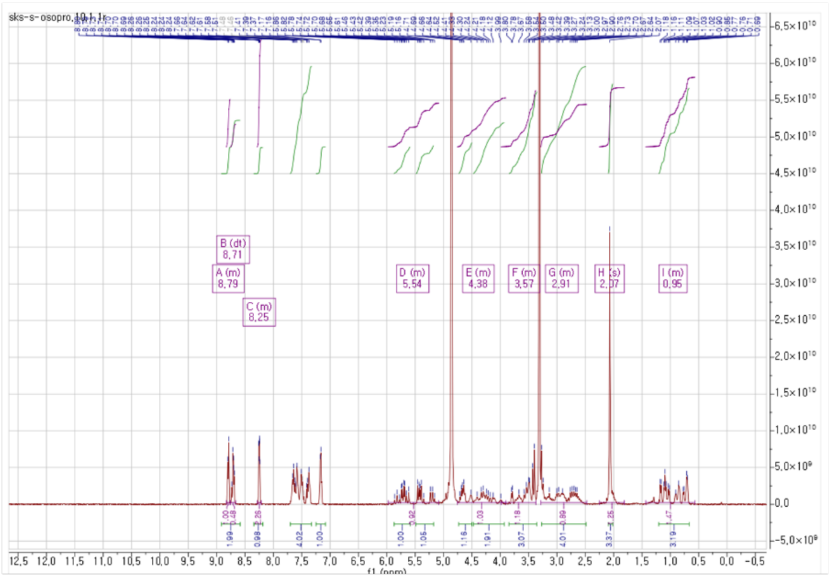

$^{1}H$ NMR, 400 MHz, $CD_3OD$

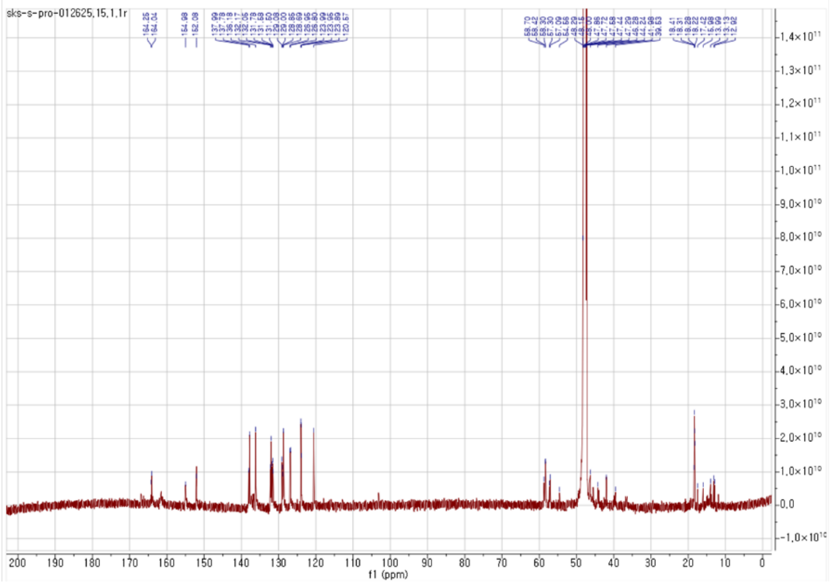

$^{13}C$ NMR, 150 MHz, $CD_3OD$

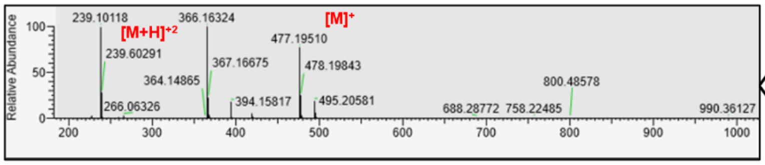


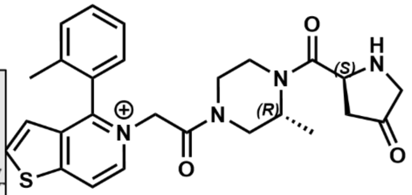

**10m**

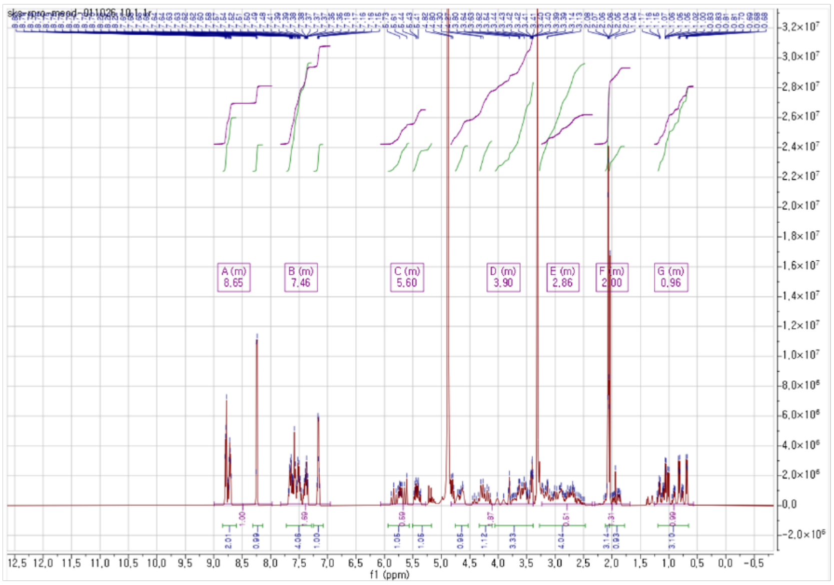


$^{1}H$ NMR, 400 MHz, $CD_3OD$

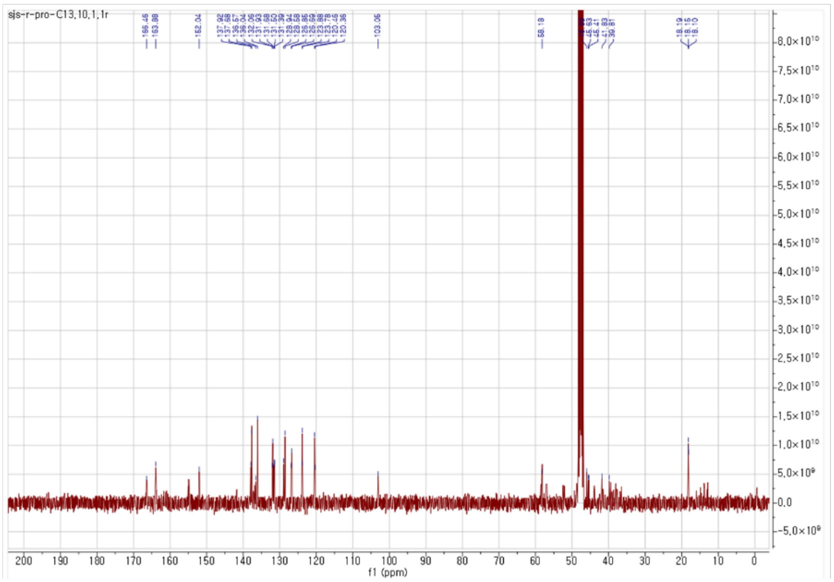


$^{13}C$ NMR, 150 MHz, $CD_3OD$

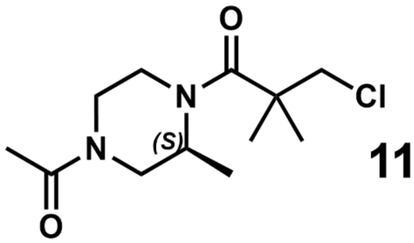

**11**

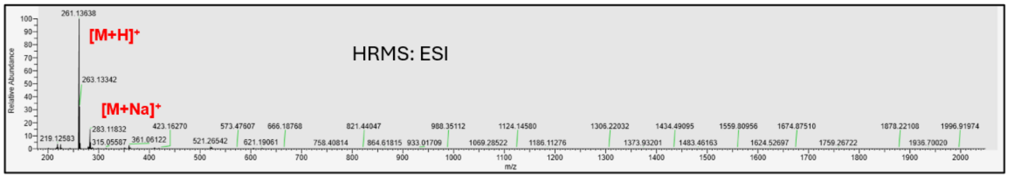


$^{1}$H NMR, 400 MHz, $d_6$-DMSO

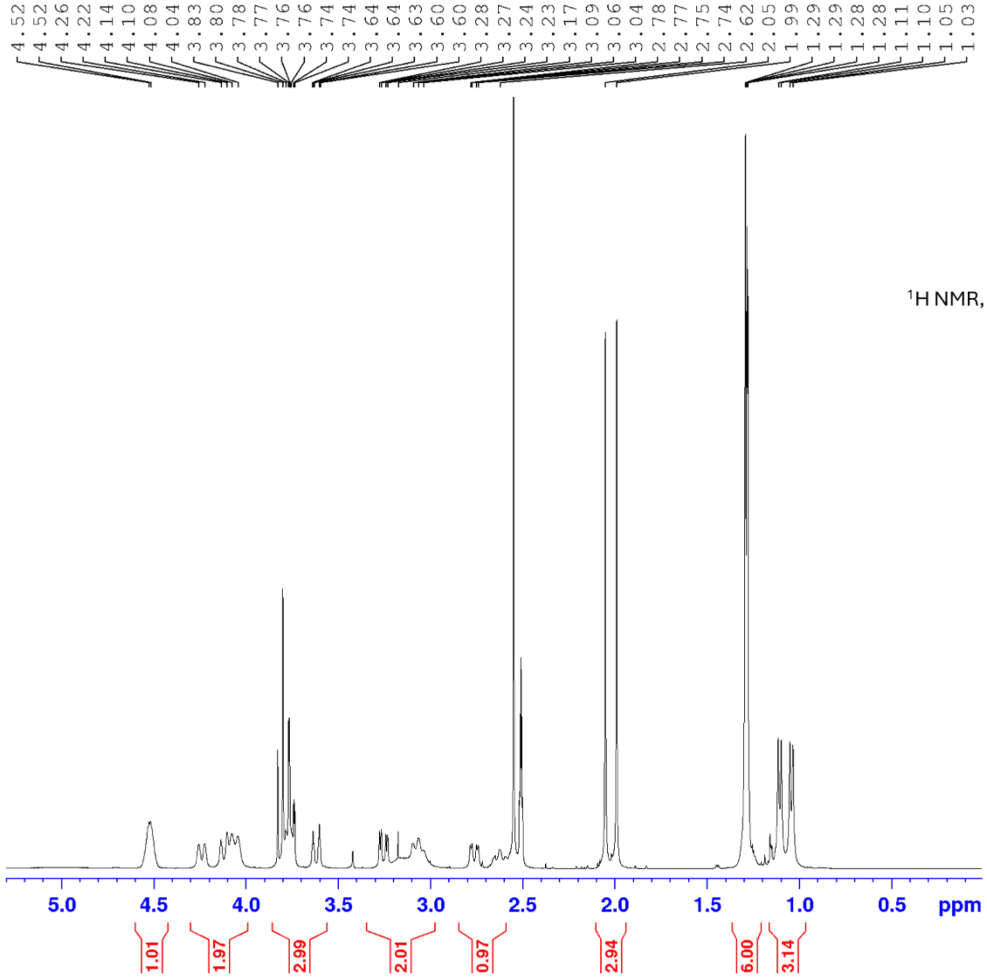

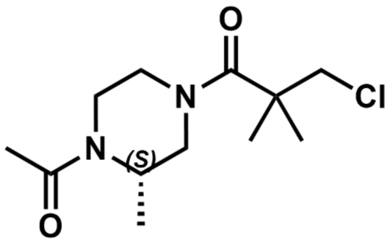

**12**

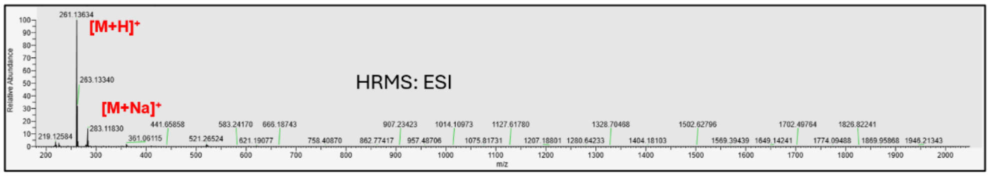


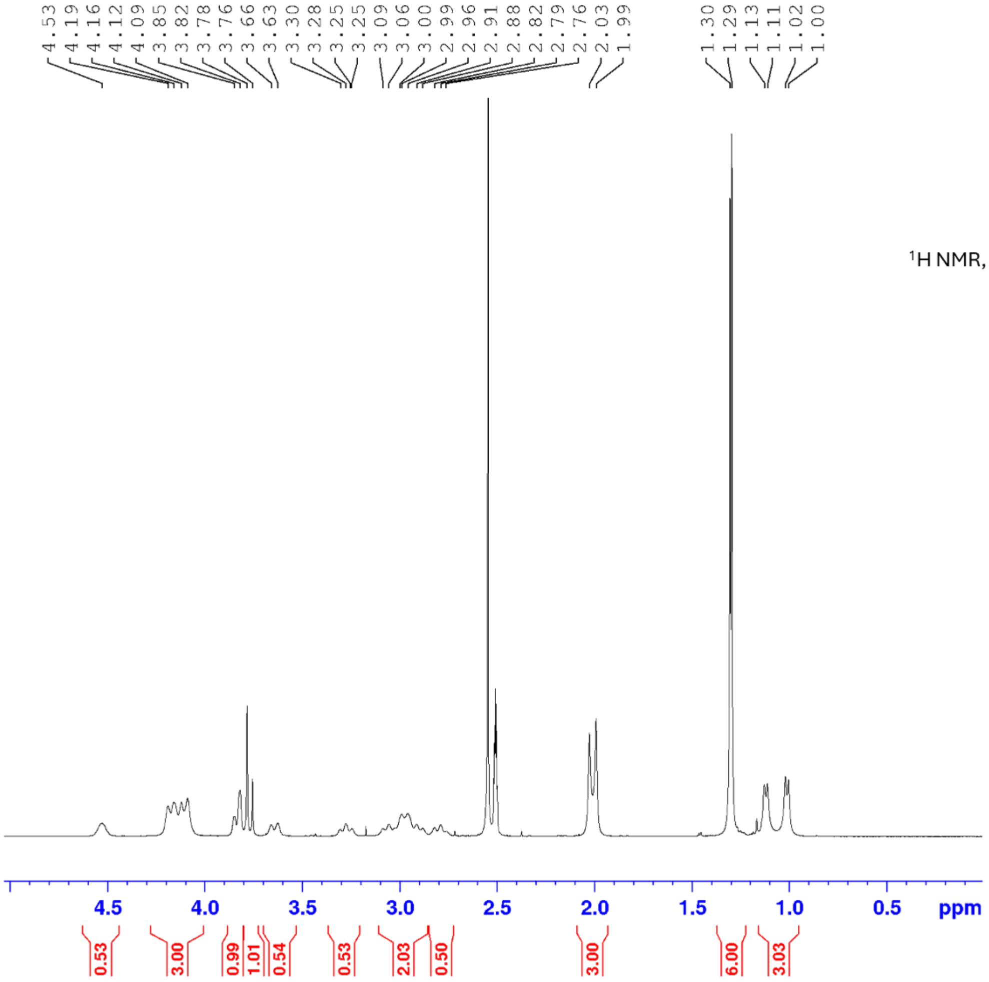


$^{1}H$ NMR, 400 MHz, $d_6$-DMSO

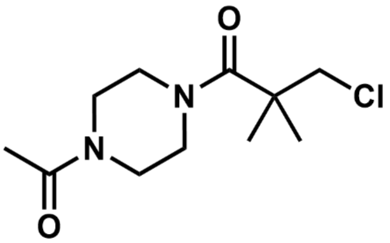

**13**

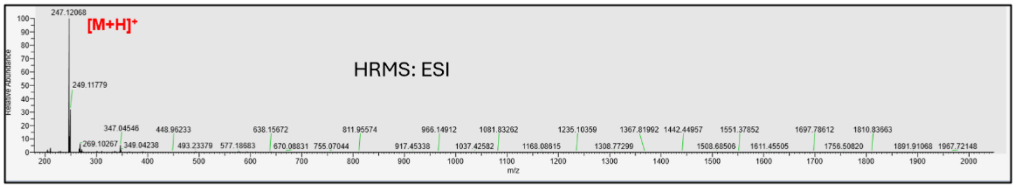
[M+H]+
HRMS: ESI
247.12068
249.11779
347.04546
448.96233
638.15672
811.95574
966.14912
1081.83262
1235.10359
1367.81992
1442.44957
1551.37852
1697.78612
1810.83663
269.10267
349.04238
493.23379
577.18683
670.08831
755.07044
917.45338
1037.42582
1168.08615
1308.77290
1508.68506
1611.45505
1756.50820
1891.91068
1967.72148
m/z
Relative Abundance

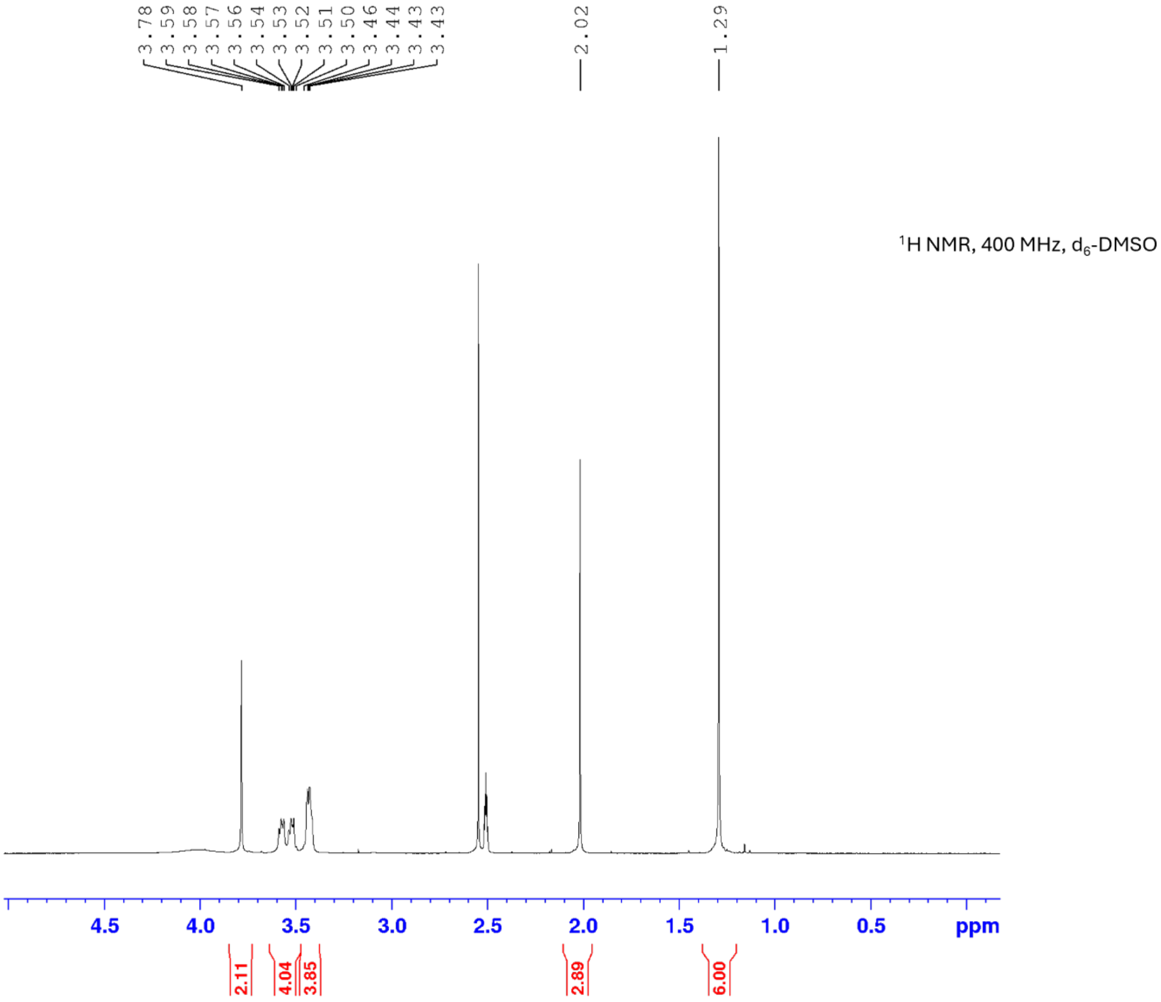
3.78
3.59
3.58
3.57
3.56
3.54
3.53
3.52
3.51
3.50
3.46
3.44
3.43
3.43
2.02
1.29
1H NMR, 400 MHz, d6-DMSO
4.5
4.0
3.5
3.0
2.5
2.0
1.5
1.0
0.5
ppm
2.11
4.04
3.85
2.89
6.00

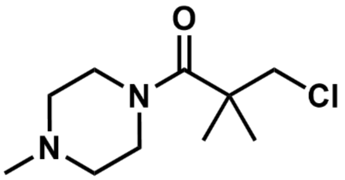

O
N
N
Cl


**14**

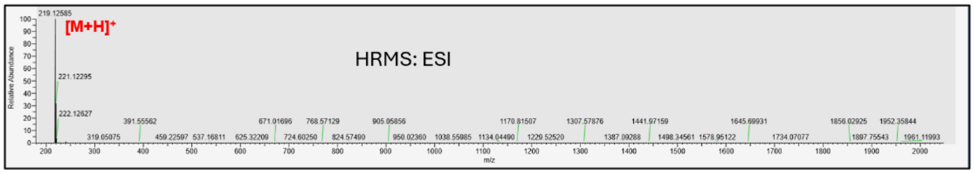

[M+H]+
HRMS: ESI
219.12585
221.12295
222.12627
m/z


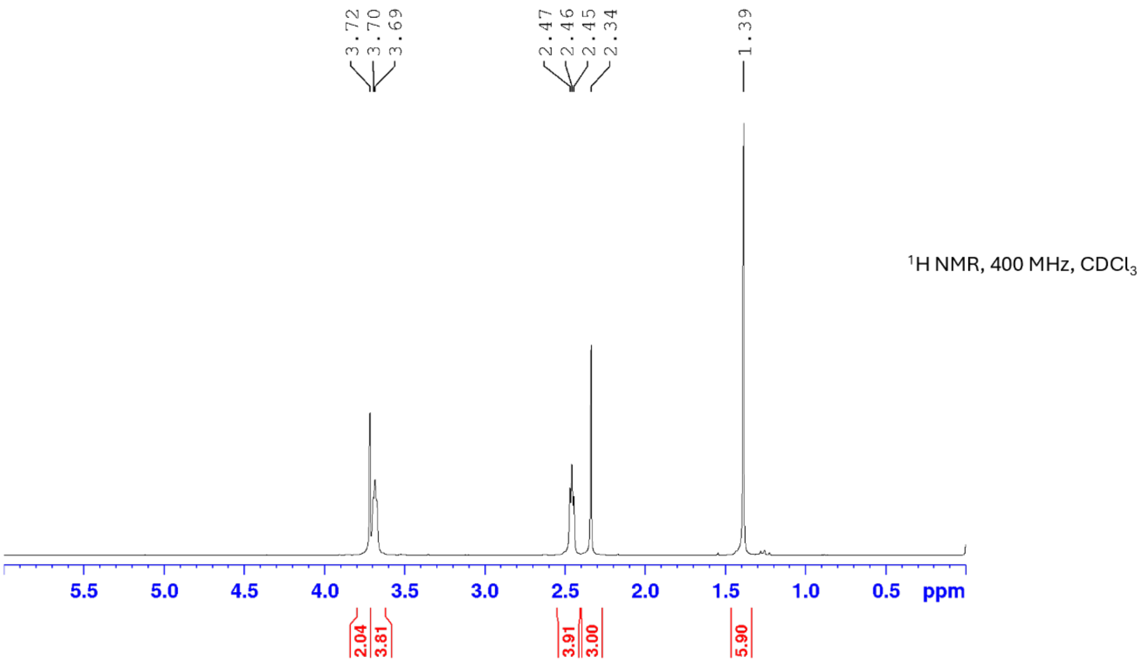

3.72
3.70
3.69
2.47
2.46
2.45
2.34
1.39
1H NMR, 400 MHz, CDCl3
ppm
2.04
3.81
3.91
3.00
5.90

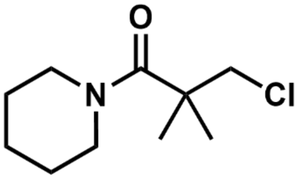


**15**

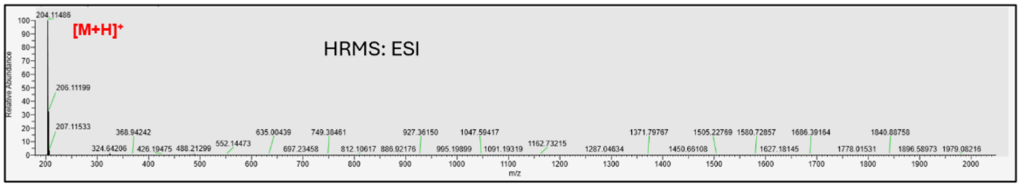


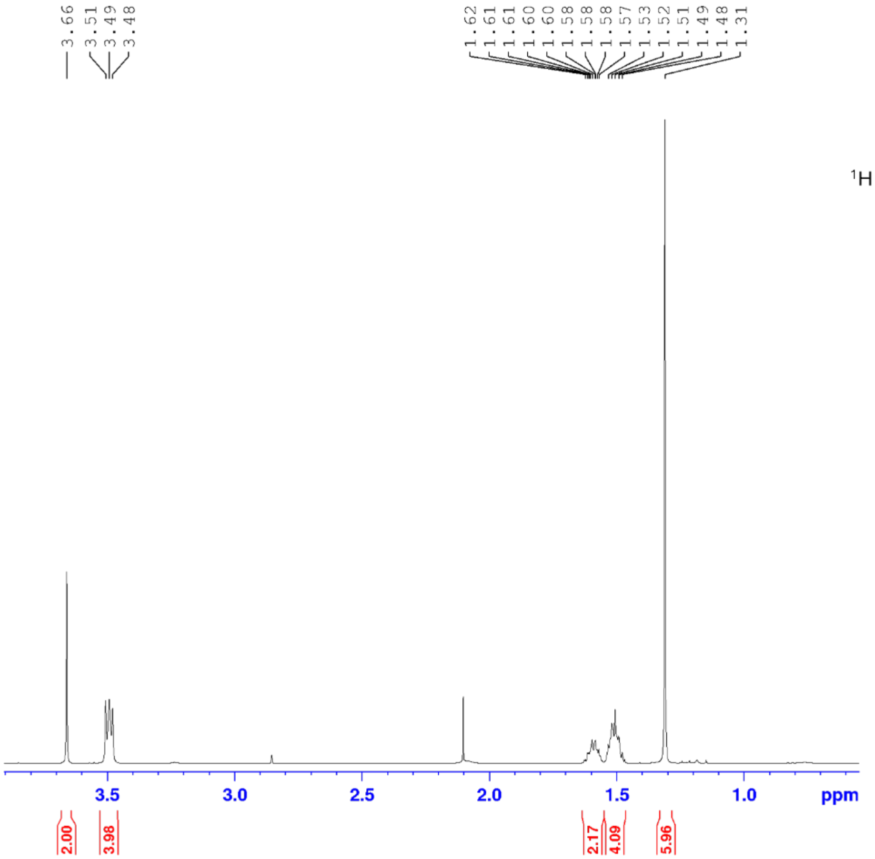


$^{1}H$ NMR, 400 MHz, $CDCl_3$

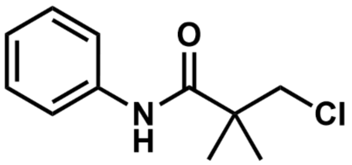
O
N
H
Cl

**16**

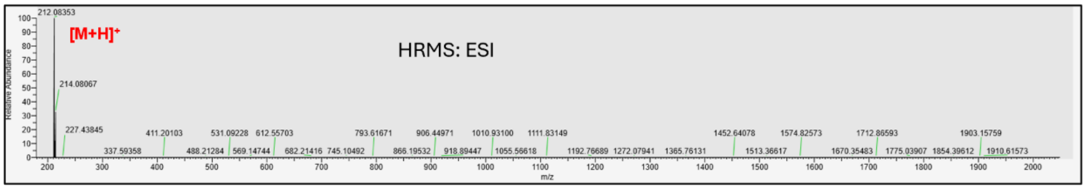
HRMS: ESI
[M+H]+
212.08353
214.08067
227.43845
337.59358
411.20103
488.21284
531.09228
569.14744
612.55703
682.21416
745.10492
793.61671
866.19532
906.44971
918.89447
1010.93100
1055.56618
1111.83149
1192.76689
1272.07941
1365.76131
1452.64078
1513.36617
1574.82573
1670.35483
1712.86593
1775.03907
1854.39612
1903.15759
1910.61573
Relative Abundance
m/z

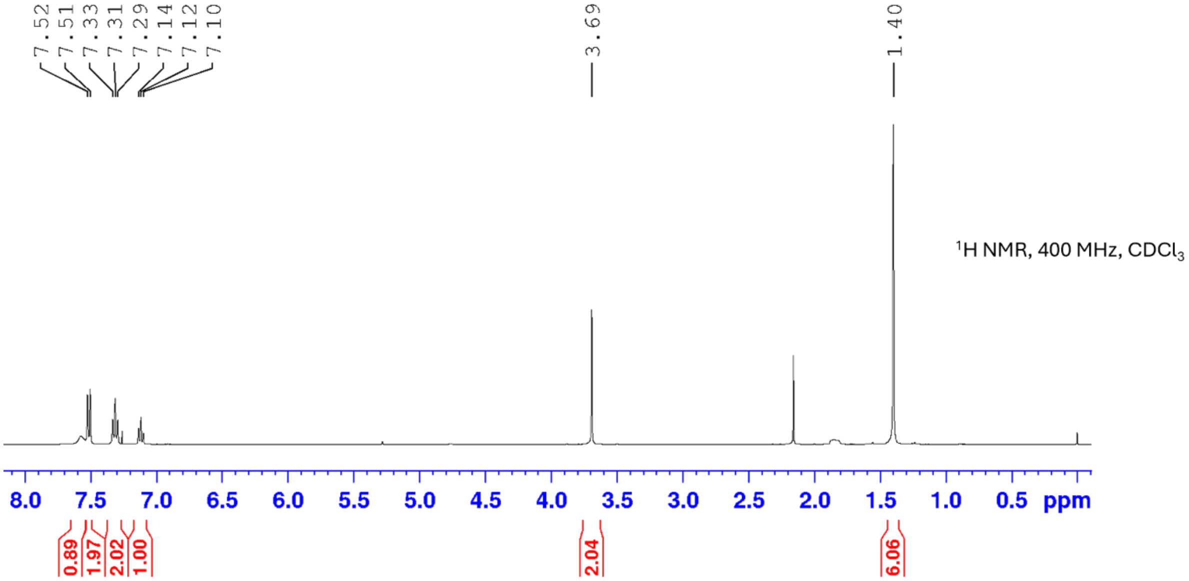
7.52
7.51
7.33
7.31
7.29
7.14
7.12
7.10
3.69
1.40
1H NMR, 400 MHz, CDCl3
8.0 7.5 7.0 6.5 6.0 5.5 5.0 4.5 4.0 3.5 3.0 2.5 2.0 1.5 1.0 0.5 ppm
0.89
1.97
2.02
1.00
2.04
6.06

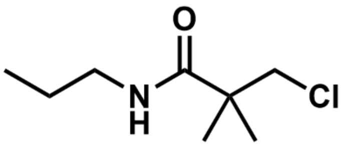

**17**

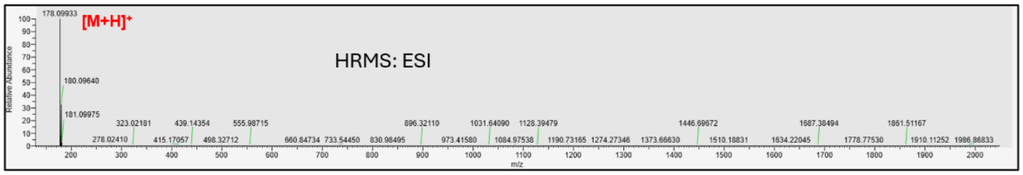
178.09933
[M+H]+
HRMS: ESI
180.09640
181.09975
Relative Abundance
m/z

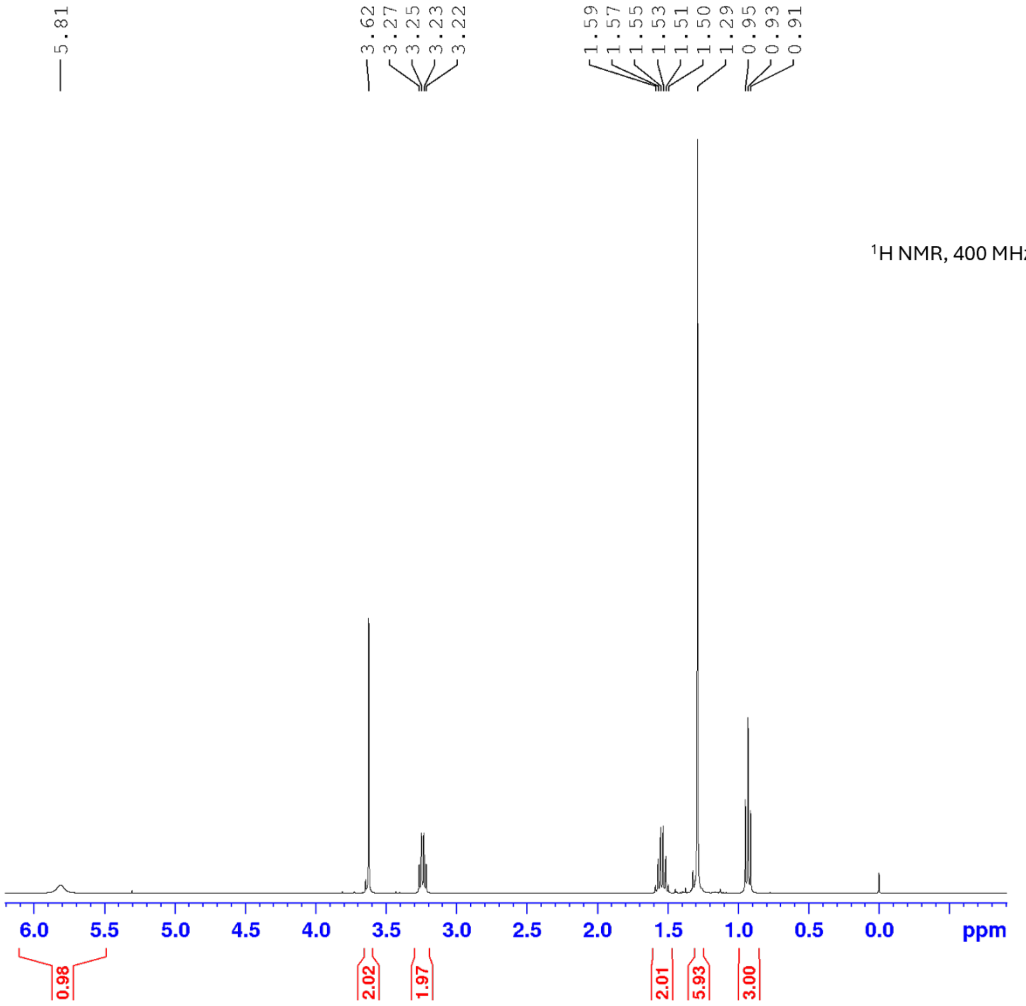
5.81
3.62
3.27
3.25
3.23
3.22
1.59
1.57
1.55
1.53
1.51
1.50
1.29
0.95
0.93
0.91
1H NMR, 400 MHz, CDCl3
ppm
0.98
2.02
1.97
2.01
5.93
3.00

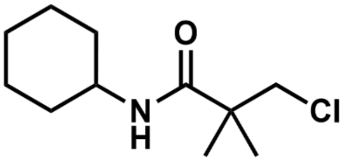

**18**

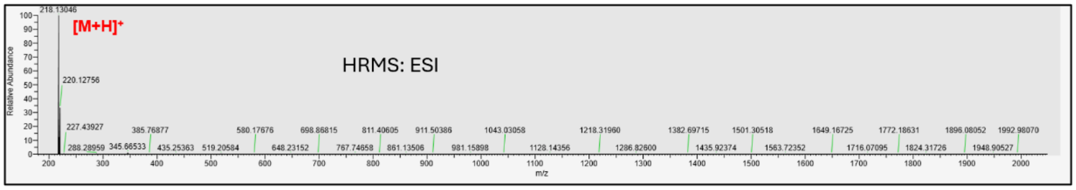


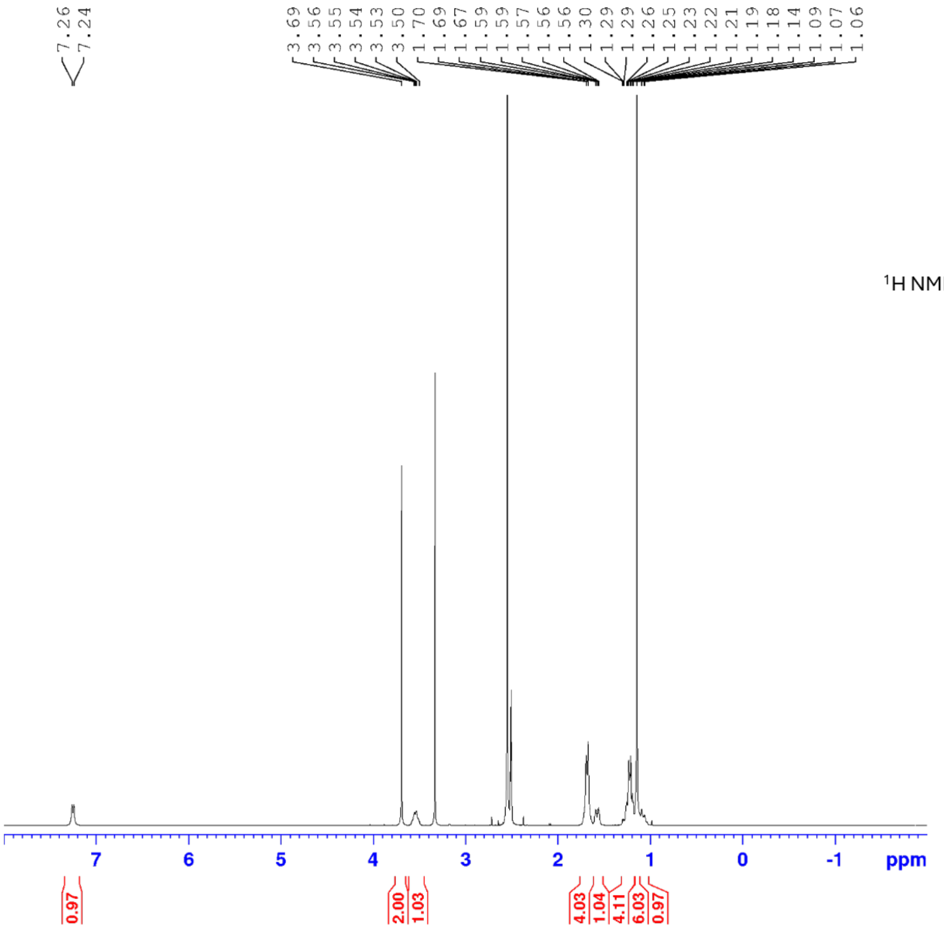


$^1$H NMR, 400 MHz, $d_6$-DMSO

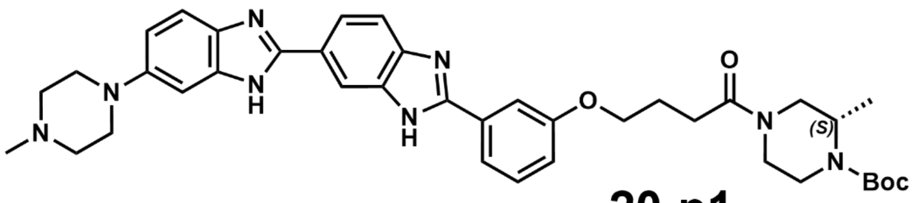

**20-p1**

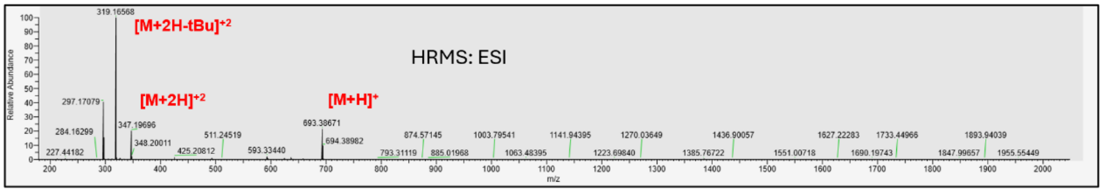

HRMS: ESI
[M+2H-tBu]+2
[M+2H]+2
[M+H]+
319.16568
297.17079
347.19696
693.38671
Relative Abundance
m/z


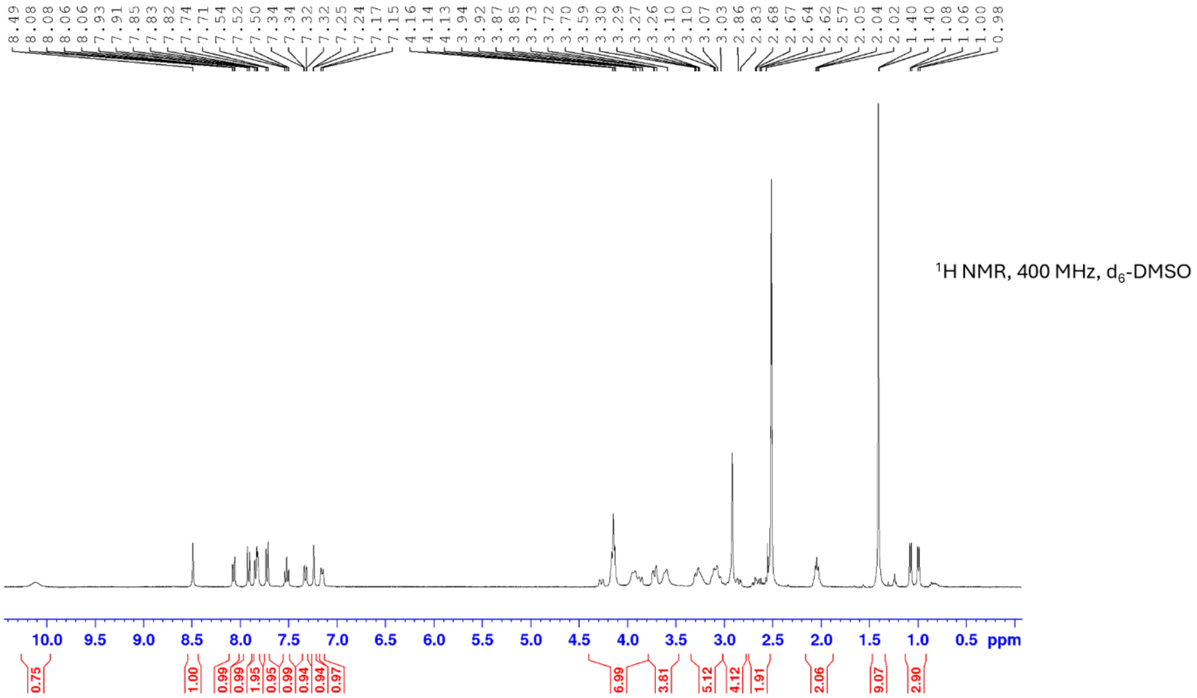

1H NMR, 400 MHz, d6-DMSO
ppm

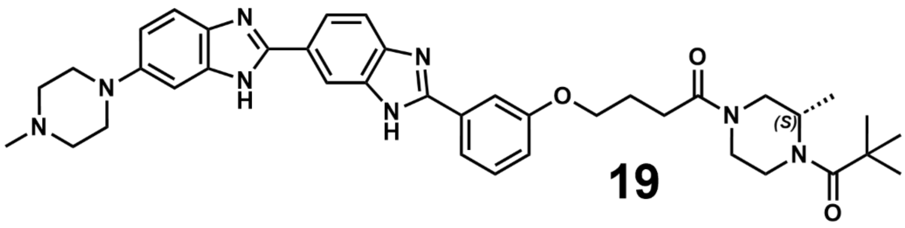

**19**

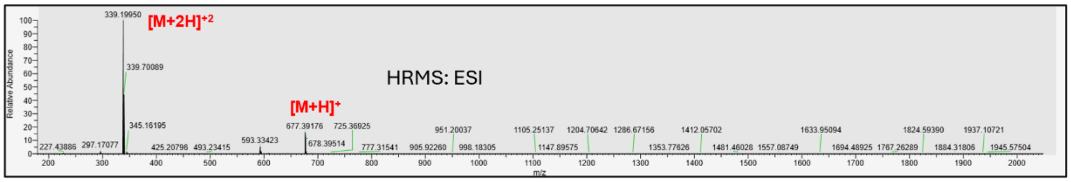


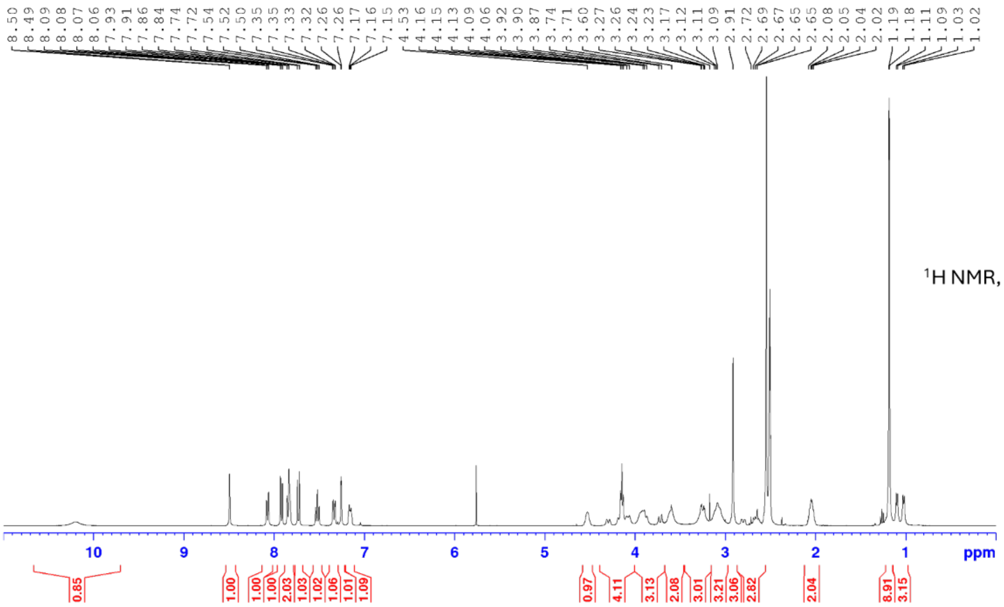


$^{1}$H NMR, 400 MHz, $d_6$-DMSO

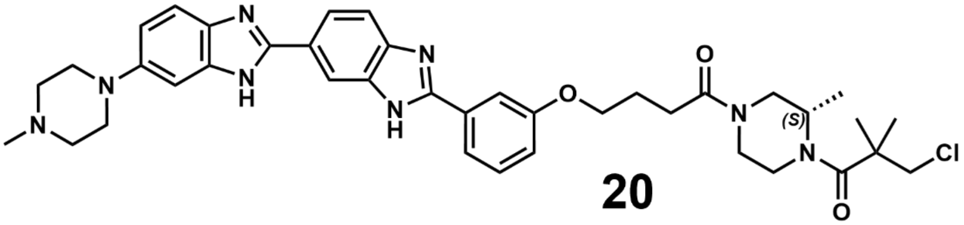

N
H
O
(S)
Cl


**20**

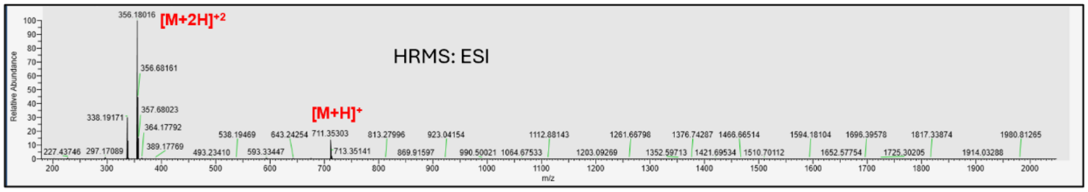

HRMS: ESI
[M+2H]+2
[M+H]+
356.18016
356.68161
357.68023
338.19171
364.17792
389.17769
711.35303
713.35141
Relative Abundance
m/z


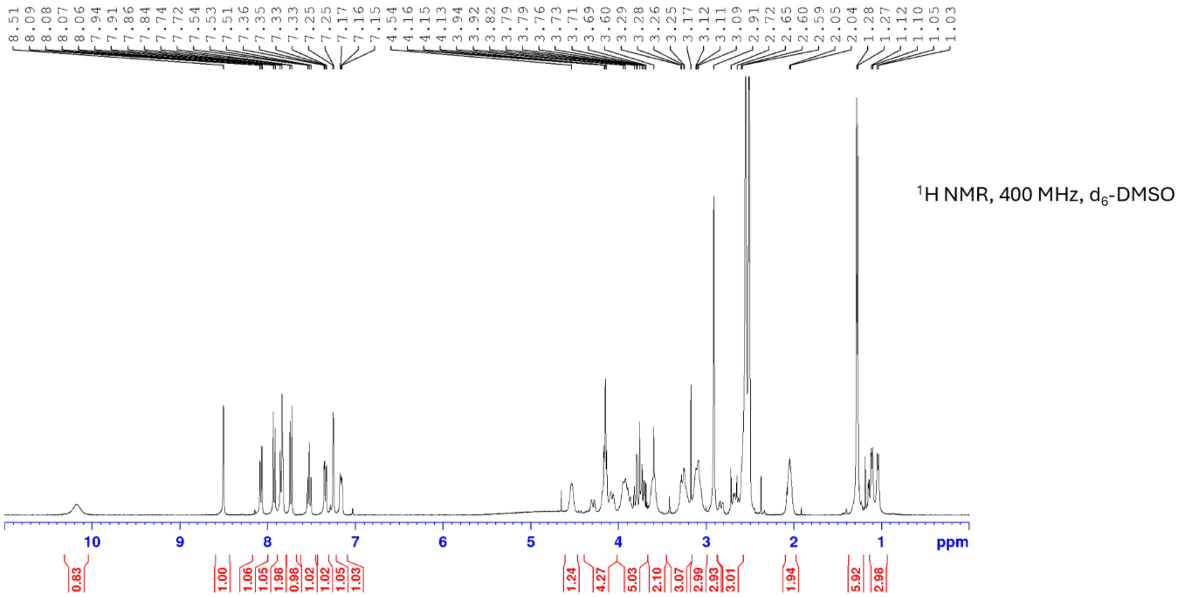

8.51 8.09 8.08 8.07 8.06 7.94 7.91 7.86 7.84 7.74 7.72 7.54 7.53 7.51 7.36 7.35 7.33 7.33 7.25 7.17 7.16 7.15 4.54 4.16 4.15 4.13 3.94 3.92 3.82 3.79 3.76 3.73 3.69 3.60 3.29 3.28 3.26 3.25 3.17 3.12 3.11 3.09 2.91 2.72 2.65 2.60 2.59 2.05 2.04 1.28 1.27 1.12 1.10 1.05 1.03
$^1$H NMR, 400 MHz, $d_6$-DMSO
ppm
0.83 1.00 1.06 1.05 1.98 0.98 1.02 1.02 1.05 1.03 1.24 4.27 5.03 2.10 3.07 2.99 2.93 3.01 1.94 5.92 2.98
